# No compelling evidence that preferences for facial masculinity track changes in women’s hormonal status

**DOI:** 10.1101/136549

**Authors:** Benedict C Jones, Amanda C Hahn, Claire I Fisher, Hongyi Wang, Michal Kandrik, Chengyang Han, Vanessa Fasolt, Danielle Morrison, Anthony J Lee, Iris J Holzleitner, Kieran J O’Shea, Craig Roberts, Anthony C Little, Lisa M DeBruine

## Abstract

Although widely cited as strong evidence that sexual selection has shaped human facial attractiveness judgments, evidence that preferences for masculine characteristics in men’s faces are related to women’s hormonal status is equivocal and controversial. Consequently, we conducted the largest ever longitudinal study of the hormonal correlates of women’s preferences for facial masculinity (N=584). Analyses showed no compelling evidence that preferences for facial masculinity were related to changes in women’s salivary steroid hormone levels. Furthermore, both within-subject and between-subject comparisons showed no evidence that oral contraceptive use decreased masculinity preferences. However, women generally preferred masculinized over feminized versions of men’s faces, particularly when assessing men’s attractiveness for short-term, rather than long-term, relationships. Our results do not support the hypothesized link between women’s preferences for facial masculinity and their hormonal status.

## Introduction

Exaggerated male sex-typical (i.e., masculine) characteristics in men have been proposed as cues of a strong immune system that would be inherited by offspring, but also linked to reduced willingness to invest time and other resources in personal relationships (Gangestad et al., 2004; Gildersleeve et al., 2014; Little et al., 2011; Perrett et al., 1998; Penton-Voak et al., 1999). Given this proposed trade off between the benefits and costs of choosing a masculine mate, researchers have hypothesized that women could maximize the benefits of their mate choices by mating with masculine men when fertile, while forming long-term relationships with relatively feminine men (Gangestad et al., 2004; Gildersleeve et al., 2014; Little et al., 2011; Penton-Voak et al., 1999).

Consistent with this hypothesis, some studies have reported that women show stronger preferences for masculine characteristics in men’s faces when in hormonal states associated with high fertility (e.g., during the ovulatory phase of the menstrual cycle and/or when not using hormonal contraceptives, Ditzen et al., 2017; Johnston et al., 2001; Little & Jones, 2012; Little et al., 2002; Little et al., 2013; Penton-Voak et al., 1999; Penton-Voak & Perrett, 2000; Roney & Simmons, 2008; Roney et al., 2011; Vaughn et al., 2010; Welling et al., 2007). These effects are widely cited as evidence that sexual selection has shaped women’s judgments of men’s facial attractiveness (Gangestad & Simpson, 2000; Grammer et al., 2003; Fink & Penton-Voak, 2002; Thornhill & Gangestad, 1999).

The claim that women’s preferences for facial masculinity are related to their hormonal status has been influential. However, it is also highly controversial (see Gildersleeve et al., 2014 and Wood et al., 2014 for meta-analyses drawing opposite conclusions about the robustness of hypothesized links between women’s masculinity preferences and hormonal status). In particular, recent work has highlighted four potentially serious methodological problems with research on the hormonal correlates of masculinity preferences.

First, sample sizes are usually small, meaning that studies are badly underpowered (Gangestad et al., 2016). For example, the mean sample size of within-subject studies reporting significant effects of hormonal status on facial masculinity preferences is 40 women (median = 34). Consequently, results from previous studies are difficult to interpret (Blake et al., 2016; Gangestad et al., 2016).

Second, hormonal status is typically assessed using self-reported menstrual cycle data (e.g., number of days since onset of last menses or number of days until expected date of next menses, Harris, 2013; Johnston et al., 2001; Little & Jones, 2012; Munoz-Reyes et al., 2014; Penton-Voak et al., 1999; Penton-Voak & Perrett, 2000; Scott et al., 2014; Zietsch et al., 2015). This method is imprecise and prone to bias (Blake et al., 2016; Gangestad et al., 2016; Harris, 2013).

Third, many studies use between-subject designs. Use of between-subject designs in this research is potentially problematic because, even with large samples, the substantial genetic contribution to individual differences in facial masculinity preferences (Zietsch et al., 2015) could obscure subtle effects of hormonal status. Thus, although several recent studies testing for possible effects of hormonal status on facial masculinity preferences have reported null results (Harris, 2013; Marcinkowska et al., 2016; Munoz-Reyes et al., 2014; Scott et al., 2014; Zietsch et al., 2015), it is noteworthy that these studies all used between-subject designs.

Fourth, studies using within-subject designs typically test women on only two occasions (Johnston et al., 2001; Little & Jones, 2012; Little et al., 2013; Penton-Voak et al., 1999; Roney et al., 2011). This limited approach may not adequately capture complex changes in hormonal status (see, e.g., Roney & Simmons, 2013).

The current study directly addressed all of these potentially serious methodological problems by recruiting 584 heterosexual women for a longitudinal (i.e., within-subject) study in which both women’s hormonal status and preferences for masculinity in men’s faces were repeatedly assessed (519 women completed at least 5 test sessions, 176 women completed at least 10 test sessions). Changes in women’s hormonal status were assessed by measuring steroid hormones from saliva samples and also by tracking within-subject changes in hormonal contraceptive use.

## Methods

### Participants

Five hundred and ninety-eight heterosexual white women who reported that they were either not using any form of hormonal contraceptive (i.e., had natural menstrual cycles) or were using the combined oral contraceptive pill were recruited for the study. Data from 14 of these women were excluded from the dataset because they reported hormonal contraceptive use inconsistently within a single block of test sessions. Thus, the final data set was 584 women (mean age=21.46 years, SD=3.09 years). Participants completed up to three blocks of test sessions (mean time between Block 1 and Block 2 = 230 days; mean time between Block 2 and Block 3 = 487 days). Each of the three blocks of test sessions consisted of five weekly test sessions. Table 1 shows how many women completed one, two, three, four, or five test sessions in Block 1, Block 2, and Block 3.

**Table 1.** The number of women who completed five, four, three, two, or one weekly test sessions in Block 1, Block 2, and Block 3.

|  | **5 sessions** | **4 sessions** | **3 sessions** | **2 sessions** | **1 session** |
| --- | --- | --- | --- | --- | --- |
| **Block 1** | 508 | 22 | 6 | 14 | 26 |
| **Block 2** | 184 | 3 | 3 | 1 | 4 |
| **Block 3** | 18 | 0 | 0 | 0 | 0 |

Forty-five women reported changing their hormonal contraceptive status between blocks during the study. Fifteen women reported changing from using the combined oral contraceptive pill to not using the combined oral contraceptive pill and 30 women reported changing from not using the combined oral contraceptive pill to using the combined oral contraceptive pill.

### Stimuli

The methods we used to manufacture stimuli to test women’s preferences for facial masculinity have been used in many previous studies (e.g., Harris, 2013; Johnston et al., 2001; Little & Jones, 2012; Marcinkowska et al., 2016; Munoz-Reyes et al., 2014; Penton-Voak et al., 1999; Penton-Voak & Perrett, 2000; Scott et al., 2014; Welling et al., 2007; Zietsch et al., 2015). Responses to stimuli manufactured using these methods predict women’s actual partner choices (DeBruine et al., 2006). They have also been shown to be very similar to responses to stimuli manufactured using other methods for manipulating sexually dimorphic characteristics in face images (DeBruine et al., 2006). We have made the stimuli from this study publicly available at osf.io/9b4y7.

First, we manufactured a female prototype (i.e., average) face by using specialist software (Tiddeman et al., 2001) to average the shape, color, and texture information from images of 50 young white women’s faces. A male prototype face was also manufactured in this way by averaging the shape, color, and texture information from images of 50 young white men’s faces.

Next, we randomly selected 10 images from the set of 50 individual male faces. We then created a feminized and a masculinized version of each of these 10 male images by adding or subtracting 50% of the linear (i.e., vector) differences in 2D shape between symmetrized versions of the female and male prototypes to (or from) each individual image. This process created 10 pairs of face images in total, with each pair consisting of a feminized and a masculinized version of one of the individual face images. Examples of these stimuli are shown in Figure 1. Note that our feminized and masculinized versions of faces differed in sexually dimorphic shape characteristics only (i.e., were matched in other regards, such as identity, color, and texture, Tiddeman et al., 2001).

**Figure 1.**
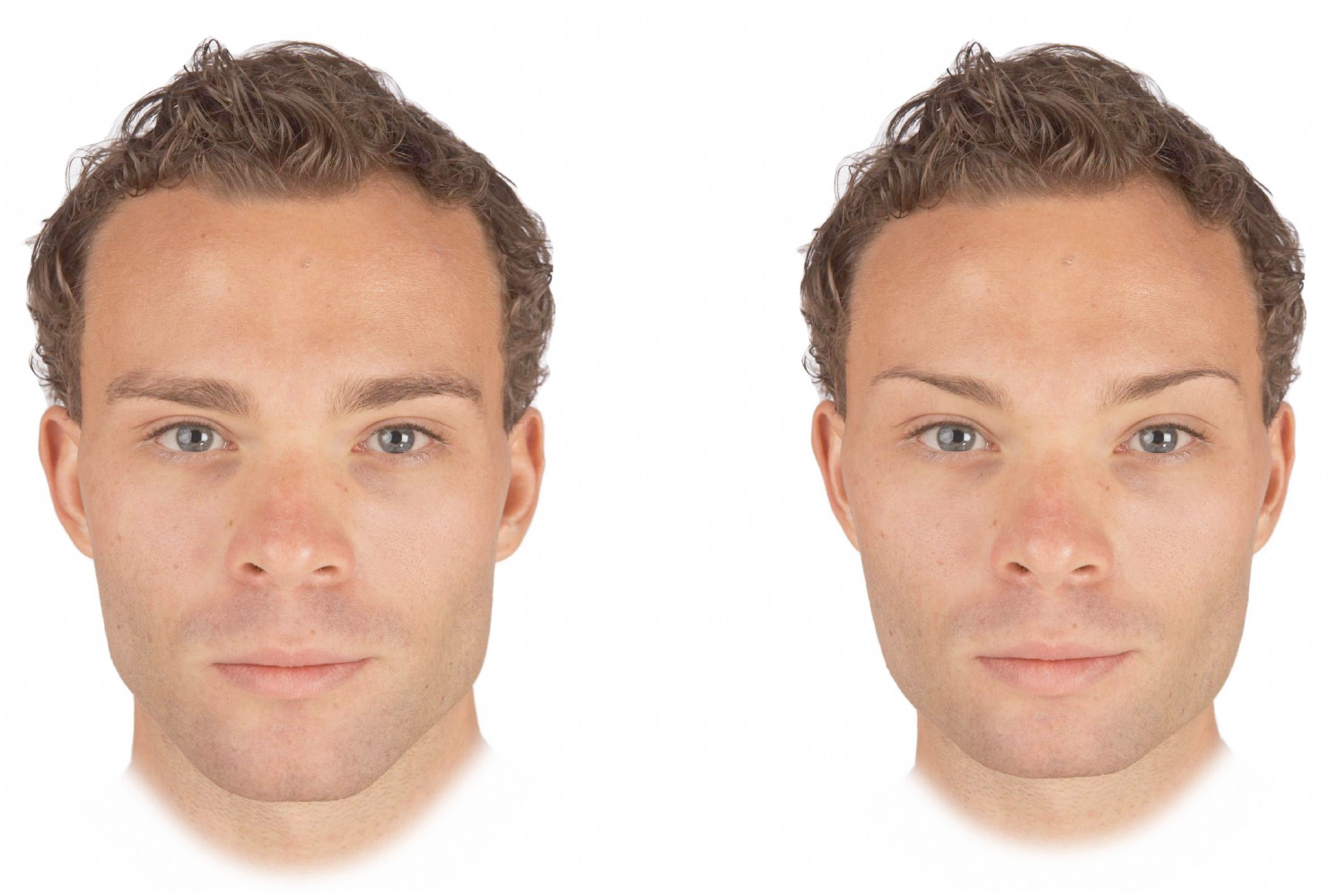
Examples of masculinized (left) and feminized (right) versions of men’s faces used to assess facial masculinity preferences in our study.

### Procedure

In each test session, women reported their current romantic partnership status (partnered or unpartnered), reported their hormonal contraceptive use status (using the combined oral contraceptive pill, not using any form of hormonal contraceptive), reported whether they were currently taking a scheduled break from the pill (and, if so, how many days into this scheduled break they were), provided a saliva sample, and completed two face preference tests (one assessing men’s attractiveness for a short-term relationship, the other assessing men’s attractiveness for a long-term relationship). Attractiveness of men for short-term relationships and long-term relationships were measured separately because hormonal status has previously been shown to influence women’s masculinity preferences when assessing men’s attractiveness for short-term, but not long-term, relationships (Little & Jones, 2012; Penton-Voak et al., 1999).

In the two face preference tests, women were shown the 10 pairs of male faces, each pair consisting of a masculinized and feminized version of a given individual. Women were instructed to select the more attractive face in each pair and to indicate the strength of that preference by choosing from the options “slightly more attractive”, “somewhat more attractive”, more attractive”, and “much more attractive”. This procedure has been used to assess masculinity preferences in previous studies (e.g., Zietsch et al., 2015).

In the short-term attractiveness test, women were told: “You are looking for the type of person who would be attractive in a short-term relationship. This implies that the relationship may not last a long time. Examples of this type of relationship would include a single date accepted on the spur of the moment, an affair within a long-term relationship, and possibility of a one-night stand.”

In the long-term attractiveness test, women were told: “You are looking for the type of person who would be attractive in a long-term relationship. Examples of this type of relationship would include someone you may want to move in with, someone you may consider leaving a current partner to be with, and someone you may, at some point, wish to marry (or enter into a relationship on similar grounds as marriage).”

Trial order within each test was fully randomized and the order in which the two face preference tests were completed in each test session was also fully randomized. Definitions of short-term and long-term relationships were taken from previous studies (Little & Jones, 2012; Penton-Voak et al., 2003).

Responses on the face preference test were coded using the following scale (higher scores indicate stronger masculinity preferences and the scale is centered on chance, i.e., zero):

0.5 to 3.5: masculinized face rated ‘slightly more attractive’ (=0.5), ‘somewhat more attractive’ (=1.5), ‘more attractive’ (=2.5) or ‘much more attractive’ (=3.5) than feminized face.

-0.5 to -3.5: feminized face rated ‘slightly more attractive’ (=-0.5), ‘somewhat more attractive’ (=-1.5), ‘more attractive’ (=-2.5) or ‘much more attractive’ (=- 3.5) than masculinized face.

Each woman’s average masculinity preference score was calculated separately for the short-term and long-term judgments for each test session. Higher scores indicate stronger masculinity preferences.

In each face preference test, the 10 trials assessing preferences for sexually dimorphic shape characteristics were interspersed among 30 filler trials assessing preferences for other facial traits.

### Saliva samples

Participants provided a saliva sample via passive drool (Papacosta & Nassis, 2011) in each test session. Participants were instructed to avoid consuming alcohol and coffee in the 12 hours prior to participation and avoid eating, smoking, drinking, chewing gum, or brushing their teeth in the 60 minutes prior to participation. Each woman’s test sessions took place at approximately the same time of day to minimize effects of diurnal changes in hormone levels (Veldhuis et al., 1988; Bao et al., 2003).

Saliva samples were frozen immediately and stored at -32°C until being shipped, on dry ice, to the Salimetrics Lab (Suffolk, UK) for analysis, where they were assayed using the Salivary 17β-Estradiol Enzyme Immunoassay Kit 1-3702 (M=3.30 pg/mL, SD=1.27 pg/mL, sensitivity=0.1 pg/mL, intra-assay CV=7.13%, inter-assay CV=7.45%), Salivary Progesterone Enzyme Immunoassay Kit 1-1502 (M=148.55 pg/mL, SD=96.13 pg/mL, sensitivity=5 pg/mL, intra-assay CV=6.20%, inter-assay CV=7.55%), Salivary Testosterone Enzyme Immunoassay Kit 1-2402 (M=87.66 pg/mL, SD=27.19 pg/mL, sensitivity<1.0 pg/mL, intra-assay CV=4.60%, inter-assay CV=9.83%), and Salivary Cortisol Enzyme Immunoassay Kit 1-3002 (M=0.23 μg/dL, SD=0.16 μg/dL, sensitivity<0.003 μg/dL, intra-assay CV=3.50%, inter-assay CV=5.08%). Only hormone levels from women not using hormonal contraceptives were used in analyses (values given above are for these women only).

Hormone levels more than three standard deviations from the sample mean for that hormone or where Salimetrics indicated levels were outside their sensitivity range were excluded from the dataset (~1% of hormone measures were excluded for these reasons). The descriptive statistics given above do not include these excluded values. Values for each hormone were centered on their subject-specific means to isolate effects of within-subject changes in hormones. They were then scaled so the majority of the distribution for each hormone varied from -.5 to .5 to facilitate calculations in the linear mixed models. Since hormone levels were centered on their subject-specific means, women with only one value for a hormone could not be included in analyses considering hormone levels.

### Analyses

Linear mixed models were used to test for possible effects of hormonal status on women’s facial masculinity preferences. Analyses were conducted using R version 3.3.2 (R Core Team, 2016), with lme4 version 1.1-13 (Bates et al., 2014) and lmerTest version 2.0-33 (Kuznetsova et al., 2013). The dependent variable was masculinity preference score, which was centered on chance. The relationship context for which women had judged men’s attractiveness was effect-coded (short-term=+0.5 and long-term=-0.5) and included as an independent variable in all analyses. Random slopes were specified maximally following Barr et al. (2013) and Barr (2013). Full model specifications and full results for each analysis are given in our Supplemental Information. Data files and analysis scripts are publicly available at osf.io/9b4y7.

## Results

### General preferences and relationship-context effect

Significant intercepts in all analyses indicated that women generally preferred masculinized to feminized versions of men’s faces. Masculinity preferences were also significantly stronger in the short-term than long-term relationship context in all analyses. The one exception was in the analyses described under Hypothesis 4. There, the relationship context effect was not significant, probably because these analyses were less powerful than our other analyses. Full results for these effects are given in our Supplemental Information.

### Hypothesis 1. Do facial masculinity preferences track changes in measured steroid hormone levels in women not using hormonal contraceptives?

The fertile phase of the menstrual cycle is characterized by the combination of high estradiol and low progesterone (Gangestad & Haselton, 2015; Puts et al., 2013). Additionally, some previous studies have suggested that changes in women’s masculinity preferences are positively correlated with changes in estradiol (Roney & Simmons, 2008; Roney et al., 2011) and negatively correlated with changes in progesterone (Jones et al., 2005; Puts, 2006). We therefore used linear mixed models to test for possible effects of estradiol, progesterone, and their interaction on women’s facial masculinity preferences. Masculinity preference scores could range from -3.5 to 3.5 (0 indicated no preference; higher scores indicated stronger masculinity preferences). This analysis included all women who were not using any form of hormonal contraceptive when tested (N=351). The specific models we used to test for hormonal correlates of within-woman changes in masculinity preferences are identical to those that we have used elsewhere to test for hormonal correlates of disgust sensitivity (Jones et al., 2017a) and sexual desire (Jones et al., 2017b). No effects involving hormone levels were significant in this analysis (all ts<0.88, all ps>.38), suggesting that women’s preferences for facial masculinity are not related to their hormonal status.

We conducted additional analyses to test for previously reported effects of testosterone (Welling et al., 2007) and cortisol (Ditzen et al., 2017) on masculinity preferences, and for hypothesized effects of estradiol-to-progesterone ratio on mating-related behavior (Eisenbruch et al., 2015). These analyses also showed no evidence that women’s preferences for masculine men were related to their hormone levels (see Supplemental Information).

At the suggestion of a reviewer, we also tested for an interaction between the effects of testosterone and cortisol (see Supplemental Information). The rationale for testing this interaction was that some research suggests that behavioral effects of testosterone are more pronounced when cortisol is low (the Dual Hormone Hypothesis, see Mehta & Prasad, 2015). Although there was a significant interaction between testosterone and cortisol (beta=0.51, SE=0.21, t=2.39, p=.018, 95% CIs=0.09, 0.93), it indicated that women’s masculinity preferences were strongest when both testosterone and cortisol were high. Since this is not the pattern of results predicted by the Dual Hormone Hypothesis, was not an a priori prediction, and was the only significant hormone effect in multiple tests for possible effects of endogenous hormones on masculinity preferences, we suggest that it is likely to be a false positive.

A reviewer also asked that we repeat each of the analyses described above controlling for effects of test session order on masculinity preferences. Doing so did not alter the patterns of results (i.e., no non-significant effects became significant and no significant effects became non-significant). These analyses are reported in our Supplemental Information.

### Hypothesis 2. Do women not using hormonal contraceptives show stronger facial masculinity preferences than women using the combined oral contraceptive pill?

Studies reporting that women not using hormonal contraceptives show stronger facial masculinity preferences than do women using hormonal contraceptives have been interpreted as converging evidence that women’s hormonal status influences their facial masculinity preferences (Little et al., 2013). To investigate this issue in our data set, we first used linear mixed models to compare the facial masculinity preferences of women using the combined oral contraceptive pill (N=212) and women not using any form of hormonal contraceptive (N=326). This analysis included all women who had reported either no use of hormonal contraceptives throughout the study or use of the combined oral contraceptive pill throughout the study (responses from women who changed contraceptive status during the study are reported under Hypothesis 4). Although there was a significant effect of oral contraceptive use in this analysis (beta=0.12, SE=0.04, t=2.75, p=.006, 95% CIs= 0.03, 0.20), the effect was such that women using the combined oral contraceptive pill showed *stronger* masculinity preferences (M=0.47, SEM=0.03) than did women not using any form of hormonal contraceptive (M=0.35, SEM=0.03). Note that stronger masculinity preferences in women using the combined oral contraceptive pill is the opposite pattern of results to what would be expected if fertility had the hypothesized positive effect on women’s masculinity preferences.

Stronger masculinity preferences in women using hormonal contraceptives have been reported in one other study (Cobey et al., 2015). We suggest that these between-group differences reflect effects of lifestyle and/or personality factors that are correlated with contraceptive use, rather than hormonal effects.

### Hypothesis 3. Do facial masculinity preferences of women using the combined oral contraceptive pill change when they are taking inactive pills?

In women using the combined oral contraceptive pill, fertility-linked hormone levels are affected when women are not taking active pills (i.e., the scheduled ‘hormone-free interval’ or ‘break’) during their monthly cycle of oral contraceptive use (van Heusden & Fauser, 2002). If women’s masculinity preferences are influenced by their hormonal status, one would then expect women’s facial masculinity preferences to change during this scheduled break. To investigate this possibility, we used linear mixed models to compare the facial masculinity preferences of women (N=173) using the combined oral contraceptive pill when they were taking active pills versus when they were taking a scheduled break from taking active pills. Note that not all women using the combined oral contraceptive pill were tested during a scheduled break. No effects involving the scheduled break were significant (both absolute ts<0.64, both ps>.52).

### Possible moderating role of partnership status

Some previous research has suggested that the magnitude of hormone-linked changes in women’s masculinity preferences is moderated by their partnership status (i.e., whether or not they had a romantic partner, Penton-Voak et al., 1999). Thus, we repeated each of the analyses described above including partnership status and all possible interactions between partnership status and the other predictors (see Supplemental Information). These additional analyses also showed no evidence that women’s salivary steroid hormone levels were related to their facial masculinity preferences or that oral contraceptive use decreased masculinity preferences.

### Hypothesis 4. Do facial masculinity preferences change when women start or stop using the combined oral contraceptive pill?

During the course of the current study, 45 women changed their hormonal contraceptive use by either switching from using no hormonal contraceptive to using the combined oral contraceptive pill, or vice versa. There was a mean time of 360 days (SD=282 days, range=56 to 1113 days) between test sessions where women were using no hormonal contraceptives and those where they were using the combined oral contraceptive pill. A previous study of 18 women’s facial masculinity preferences reported that women’s preferences for masculinity in men’s faces decreased when women started using oral contraceptives (Little et al., 2013). We therefore used linear mixed models to compare the facial masculinity preferences of these women when they were using the combined oral contraceptive pill and when they were using no form of hormonal contraceptive. Our analysis controlled for the direction of change in women’s oral contraceptive use (i.e., whether they changed from using no form of hormonal contraceptive to using the combined oral contraceptive pill, N=30; or vice versa, N=15). The effect of oral contraceptive use was not significant (beta=0.08, SE=0.05, t=1.57, p=.12, 95% CIs = -0.02, 0.17). Note that women’s masculinity preferences tended to be stronger when they were using the combined oral contraceptive pill (although not significantly so), suggesting that a lack of power did not prevent detection of the hypothesized weaker masculinity preferences when women are using the combined oral contraceptive pill.

Because changes in oral contraceptive use could be associated with a change in partnership status, we repeated this analysis controlling for possible effects of changes in women’s partnership status (see Supplemental Information). This additional analysis also did not show any evidence that using the combined oral contraceptive pill weakened women’s masculinity preferences.

### Preferences for additional facial traits

Some previous studies have tested for effects of hormonal status on other aspects of women’s face preferences, such as preferences for femininity in women’s faces, facial symmetry, facial averageness, and apparent facial health (reviewed in Jones et al., 2008). Consequently, we also tested for effects of hormonal status on women’s preferences for these facial characteristics.

All male face preferences were assessed in the same short-term and longterm blocks with trial order fully randomized. All female face preferences were tested in a separate block, again with trial order fully randomized. The order in which women completed the short-term male attractiveness, long-term male attractiveness, and female attractiveness preference tasks in each test session was fully randomized. Femininity in women’s faces was manipulated using identical methods to those that were used to manipulate masculinity in men’s faces. Methods used to manipulate facial symmetry, facial averageness, and apparent facial health are reported in Quist et al. (2012), Jones et al. (2007), and Wincenciak et al. (2015), respectively.

Analyses of these preferences using the same type of models we used to test for effects of hormonal status on masculinity preferences also showed no clear evidence that face preferences were consistently related to women’s hormonal status. Notably, we did not replicate putative effects of ovarian hormones on women’s preferences for symmetry or apparent health previously reported for women not using hormonal contraceptives (reviewed in Jones et al., 2008). Full results, along with the data, analysis files, and stimuli, are publicly available at osf.io/9b4y7. These full results include a significant negative effect of cortisol on preferences for male facial symmetry and a significant negative effect of progesterone on preferences for male facial averageness. Neither of these results were a priori predictions, so we suggest they should be treated as preliminary findings.

## Discussion

Collectively, our analyses showed no compelling evidence that changes in women’s salivary hormone levels are associated with their facial masculinity preferences or that the combined oral contraceptive pill decreases women’s masculinity preferences^1^. This was despite having a much larger sample size, having tested participants more often, and having used more reliable measures of hormonal status (e.g., measurements of multiple steroid hormones from saliva samples) than previous studies. Thus, the current study presents evidence against the popular and influential hypothesis that changes in women’s facial masculinity preferences track changes in their hormonal status (Ditzen et al., 2017; Johnston et al., 2001; Little & Jones, 2012; Little et al., 2013; Penton-Voak et al., 1999; Penton-Voak & Perrett, 2000). Analyses of preferences for other facial traits (symmetry, averageness, apparent health) that some previous research had suggested may track changes in hormonal status also showed no compelling evidence for consistent effects of hormonal status on face preferences. Although we did observe a significant negative effect of cortisol on preferences for male facial symmetry and a significant negative effect of progesterone on preferences for male facial averageness, these findings were not predicted a priori and should be treated as preliminary. Indeed, given that symmetry and averageness are correlated in faces (see Jones et al., 2007), it is unclear why steroid hormones would have different effects on preferences for these facial characteristics.

A crucial piece of the rationale for predicting hormone-linked changes in women’s preferences for facial masculinity is the claim that facial masculinity is a cue of men’s heritable immunocompetence (Penton-Voak et al., 1999). Our null results for hormonal status and facial masculinity preferences add to a growing body of evidence calling this assumption into question (Lee et al., 2014; Scott et al., 2014). Rather than functioning as a cue of men’s immunocompetence, men’s facial masculinity may primarily function as a cue of their intrasexual competitiveness (reviewed in Puts, 2010).

Although we find no evidence that women’s masculinity preferences are linked to their hormonal status, our analyses do suggest that women show stronger preferences for masculine facial characteristics when assessing men’s attractiveness for short-term relationships than when assessing men’s attractiveness for long-term relationships. Although this pattern of results is consistent with the proposal that perceived costs associated with choosing a masculine mate cause women’s preferences for masculinity in long-term partners to be weaker than preferences for masculinity in short-term partners (Little et al., 2011), we emphasize here that the effect of relationship context on masculinity preference was small.

In summary, and by contrast with previous research using smaller samples and less precise measures of hormonal status, our analyses show no compelling evidence for links between women’s hormonal status and preferences for facial masculinity. These results highlight the importance of employing large sample sizes and rigorous assessments of hormonal status (e.g., measures of salivary hormone levels) to test hypotheses concerning links between hormonal status and mate preferences.

## Acknowledgments

This research was supported by European Research Council grants awarded to BCJ (OCMATE) and LMD (KINSHIP). We thank Sean Murphy, Ruben Arslan, Hans IJzerman, Steve Gangestad, and Lawrence Barsalou for helpful discussion and comments, and thank the Editor and two reviewers for thoughtful and constructive comments.

## Male Facial Masculinity

BC Jones et al. ()

- Overview
- Hypothesis 1

∘ Descriptive stats: data_hormones

■ Preferences
■ Hormones
∘ Analyses H1: Hormones

■ E + P + E^*^P:
■ E + P + EPratio:
■ T + C:
■ T + C + T^*^C:
∘ Analyses H1s: Hormones (+ session order)

■ E + P + E^*^P: (+ session order)
■ E + P + EPratio: (+ session order)
■ T + C: (+ session order)
■ T + C + T^*^C: (+ session order)
∘ Descriptive stats: data_hormones_partner
∘ Analyses H1p: Hormones (+ partnership status)

■ E + P + E^*^P: (+ partnership status)

■ E + P + E*P: (single women only to interpret interaction)
■ E + P + E*P: (partnered women only to interpret interaction)
■ E + P + EPratio: (+ partnership status)
■ T + C: (+ partnership status)
■ T + C + T^*^C: (+ partnership status)
∘ Analyses H1ps: Hormones (+ session order, + partnership status)

■ E + P + E^*^P: (+ session order, + partnership status)

■ E + P + E^*^P: (single women only to interpret interaction)
■ E + P + E^*^P: (partnered women only to interpret interaction)
■ E + P + EPratio: (+ session order, + partnership status)
■ T + C: (+ session order, + partnership status)
■ T + C + T^*^C: (+ session order, + partnership status)
- Hypothesis 2

∘ Descriptive stats: data_between
∘ Analyses H2: Pill
∘ Analyses H2p: Pill (+ partnership status)

■ Pill (single women only to interpret interaction)
■ Pill (partnered women only to interpret interaction)
- Hypothesis 3

∘ Descriptive stats: data_pillbreak
∘ Analyses H3: Pill-break
∘ Analyses H3p: Pill-break (+ partnership status)

■ Pill break (single women only to interpret interaction)
■ Pill break (partnered women only to interpret interaction)
- Hypothesis 4

∘ Descriptive stats: data_pill_switchers

■ Interval between pill use and non-use testing blocks
∘ Analyses H4: Pill-switch
∘ Analyses H4p: Pill-switch (+ partnership status change)

## Overview

This supplemental information contains the R code for data analysis of **male facial masculinity** preferences reported in the manuscript (data are publicly available at https://osf.io/9b4y7/ (https://osf.io/9b4y7/)).

*Hypothesis 1*. Do preferences track changes in measured steroid hormone levels in women not using hormonal contraceptives?

Women reporting no use of hormonal contraceptives and for whom at least two test sessions with valid hormone levels are available.

*Hypothesis 2*. Do women not using hormonal contraceptives show stronger preferences than women using the combined oral contraceptive pill?

Women reporting use of the combined oral contraceptive pill or no use of hormonal contraceptives across all test sessions.

*Hypothesis 3*. Do preferences of women using the combined oral contraceptive pill change when they are taking inactive pills?

Women who were tested during a scheduled break from use of the combined oral contraceptive pill.

*Hypothesis 4*. Do preferences change when women start or stop using the combined oral contraceptive pill?

Women who switched from using no hormonal contraceptive to using the combined oral contraceptive pill (or vice versa) between blocks of test sessions.

~~~
## Loading tidyverse: ggplot2
## Loading tidyverse: tibble
## Loading tidyverse: tidyr
## Loading tidyverse: readr
## Loading tidyverse: purrr
## Loading tidyverse: dplyr
~~~

~~~
## Conflicts with tidy packages ----------------------------------------------
~~~

~~~
## filter(): dplyr, stats
## lag():    dplyr, stats
~~~

~~~
## Loading required package: Matrix
~~~

~~~
##
## Attaching package: ‘Matrix’
~~~

~~~
## The following object is masked from ‘package:tidyr’:
##
##     expand
~~~

~~~
##
## Attaching package: ‘lmerTest’
~~~

~~~
## The following object is masked from ‘package:lme4’:
##
##     lmer
~~~

~~~
## The following object is masked from ‘package:stats’:
##
##     step
~~~

~~~
##
## Attaching package: ‘lubridate’
~~~

~~~
## The following object is masked from ‘package:base’:
##
##     date
~~~

~~~
*# load data frames from Jones_hormones_data.Rmd*
data_hormones <- readRDS(“data_hormones.Rda”)
data_hormones_partner <- readRDS(“data_hormones_partner.Rda”)
data_between <- readRDS(“data_between.Rda”)
data_between_partner <- readRDS(“data_between_partner.Rda”)
data_pillbreak <- readRDS(“data_pillbreak.Rda”)
data_pillbreak_partner <- readRDS(“data_pillbreak_partner.Rda”)
data_pill_switchers <- readRDS(“data_pill_switchers.Rda”)
data_pill_switchers_partner <- readRDS(“data_pill_switchers_partner.Rda”)
~~~

~~~
*# filter only sexual dimorphism manipulations for male faces in the analyses below*
theManip <- “sexdim”
theFaceSex <- “men”
confint_method <- “Wald” *# c(“profile”, “Wald”, “boot”)*
~~~

~~~
*# calculate standard errors*
se <- **function**(x, na.rm = FALSE) {
   **if** (na.rm) {
     the.SE <- sqrt(var(x,na.rm=TRUE)/length(na.omit(x)))
   } **else** {
     the.SE <- sqrt(var(x,na.rm=FALSE)/length(x))
}
 **return**(the.SE)
}
*# short summaries for lmerTest*
mySummary <- **function**(lmer_summary) {
 coefTable <- lmer_summary$coefficients %>%
   round(3) %>%
   as.data.frame() %>%
   rownames_to_column()
 **if** (ncol(coefTable)>5) {
   coefTable <- coefTable %>%
    mutate(
     sig = ifelse(.[6]<.001, “***”,
           ifelse(.[6]<.01, “**”,
           ifelse(.[6]<.05, “*”,
           ifelse(.[6]<.10, “+”, ““)))))
  }
 **return**(list(lmer_summary$ngrps, kable(coefTable)))
}
~~~

### Hypothesis 1

Do preferences track changes in measured steroid hormone levels in women not using hormonal contraceptives?

For tests of effects of endogenous hormones. Women reporting no use of hormonal contraceptives and for whom at least two test sessions with valid hormone levels are available (all relationship statuses).

#### Descriptive stats: data_hormones

##### Preferences

~~~
*# create mean DV for all ratings by oc_id*
stats_overall <- filter(data_hormones, face_sex==theFaceSex, manip==theManip)
%>%
 group_by(oc_id) %>%
 summarise(
   overall_rating.c = mean(rating.c)
 ) %>%
 ungroup() %>%
 group_by() %>%
 summarise(
   context=“overall”,
   n = n_distinct(oc_id),
   mean_dv = mean(overall_rating.c),
   sd_dv = sd(overall_rating.c),
   se_dv = se(overall_rating.c)
 )
*# create mean DV splitting by context*
stats_context <- filter(data_hormones, face_sex==theFaceSex, manip==theManip) %>%
 group_by(oc_id, context) %>%
 summarise(
   context_rating.c = mean(rating.c)
 ) %>%
 group_by(context) %>%
 summarise(
   n = n_distinct(oc_id),
   mean_dv = mean(context_rating.c),
   sd_dv = sd(context_rating.c),
   se_dv = se(context_rating.c)
 )
rbind(stats_overall, stats_context)
~~~

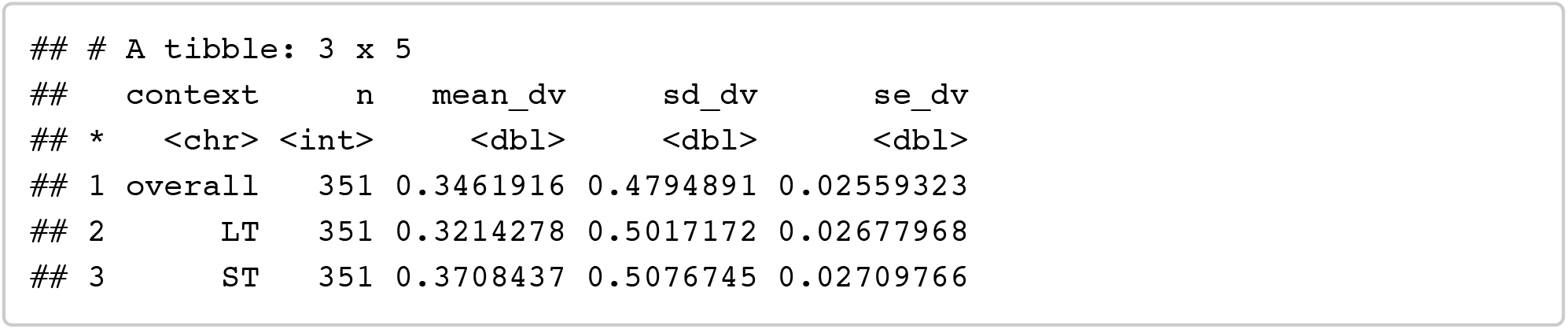

#### Hormones

~~~
filter(data_hormones, face_sex==theFaceSex, manip==theManip) %>%
   group_by() %>%
   summarise(
      mean_prog = mean(prog, na.rm = TRUE),
      sd_prog =sd(prog, na.rm = TRUE),
      se_prog =se(prog, na.rm = TRUE),
      mean_estr = mean(estr, na.rm = TRUE),
      sd_estr =sd(estr, na.rm = TRUE),
      se_estr =se(estr, na.rm = TRUE),
      mean_test = mean(test, na.rm = TRUE),
      sd_test =sd(test, na.rm = TRUE),
      se_test =se(test, na.rm = TRUE),
      mean_cort = mean(cort, na.rm = TRUE),
      sd_cort =sd(cort, na.rm = TRUE),
      se_cort =se(cort, na.rm = TRUE)
  ) %>% gather(“stat”, “value”, 1:length(.)) %>%
    mutate(value = round(value, 4)) %>%
    separate(stat, c(“stat”, “hormone”)) %>%
    spread(stat, value)
~~~

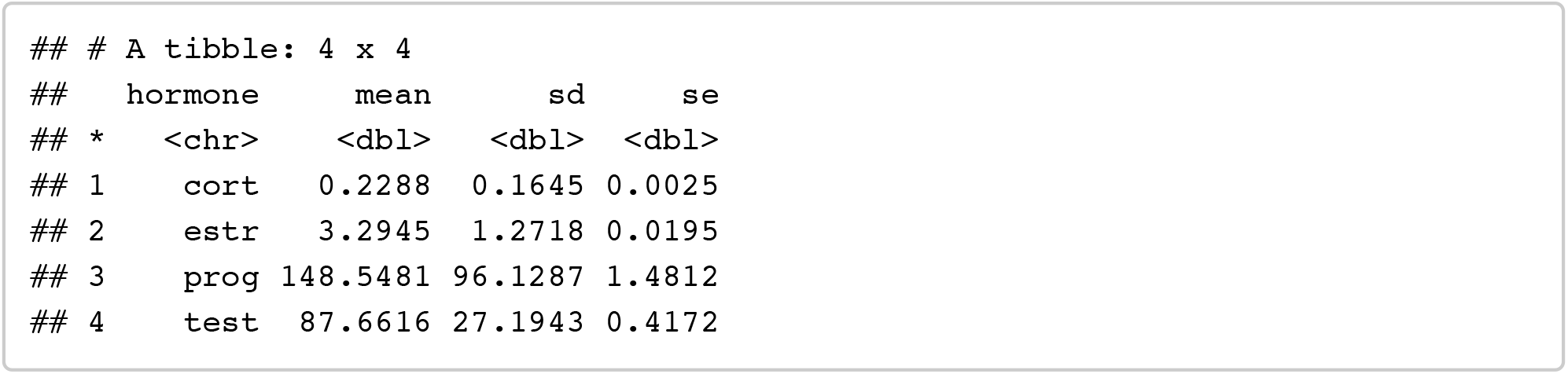

#### Analyses H1: Hormones

##### E + P + E^*^P

Testing for effects of estradiol, progesterone, and their interaction on preferences

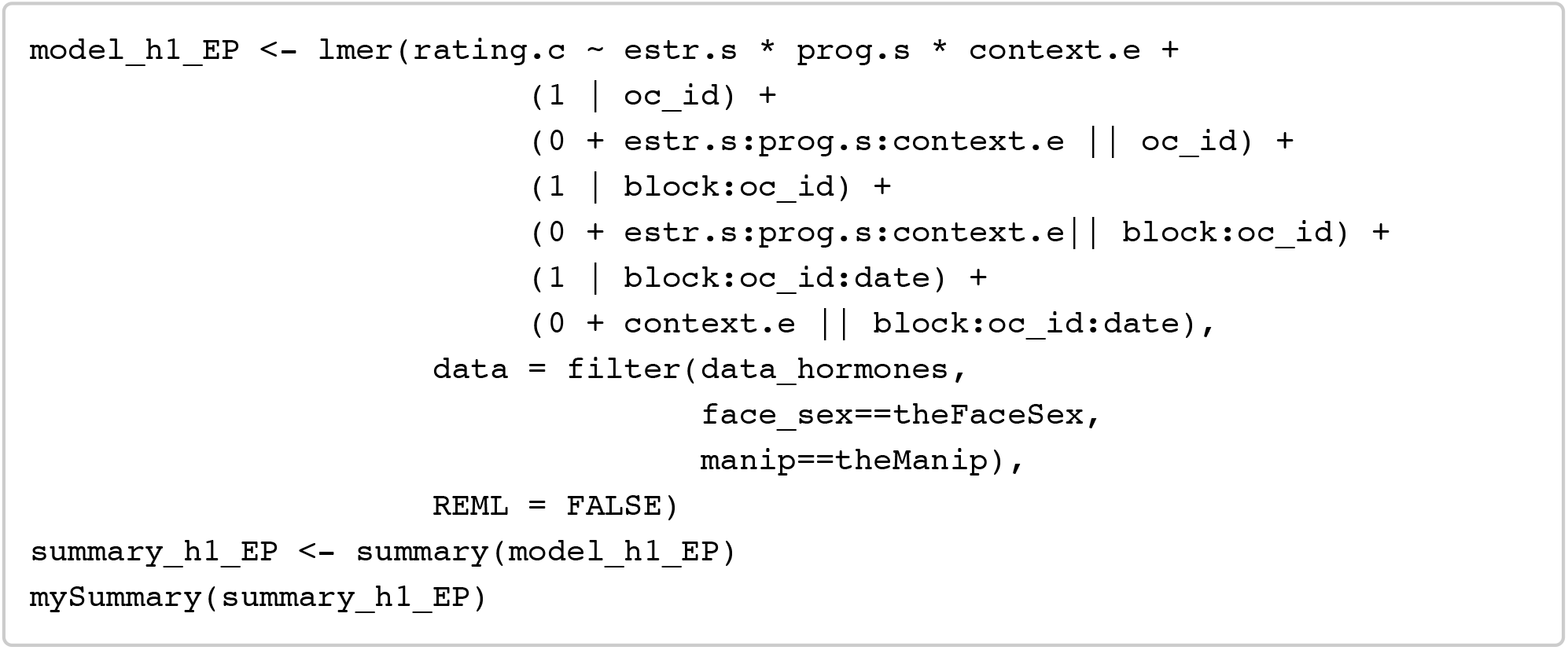

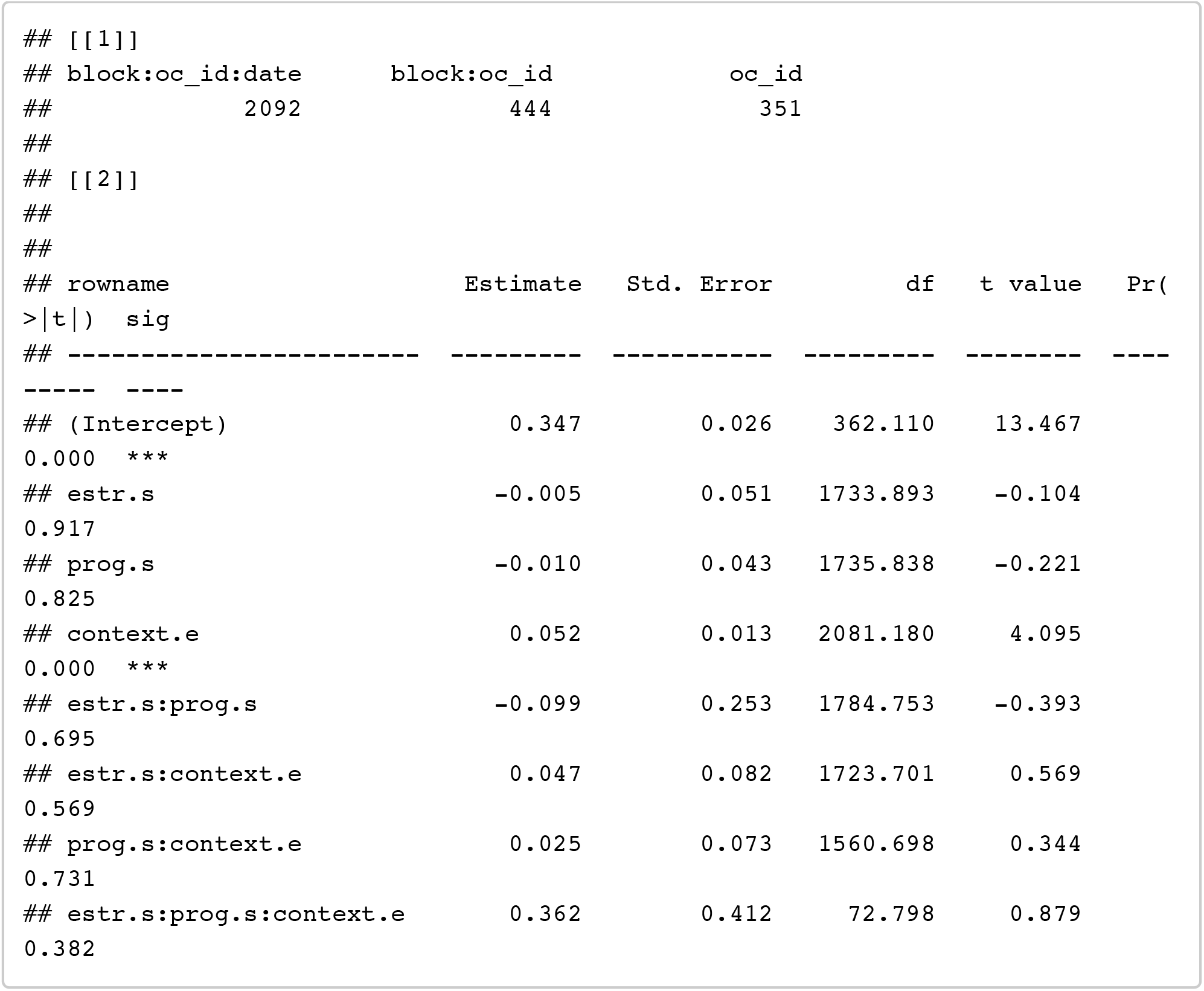

~~~
confint(model_h1_EP, method = confint_method) %>% as.data.frame() %>% rownames _to_column() %>% filter(!is.na(‘2.5 %’))
~~~

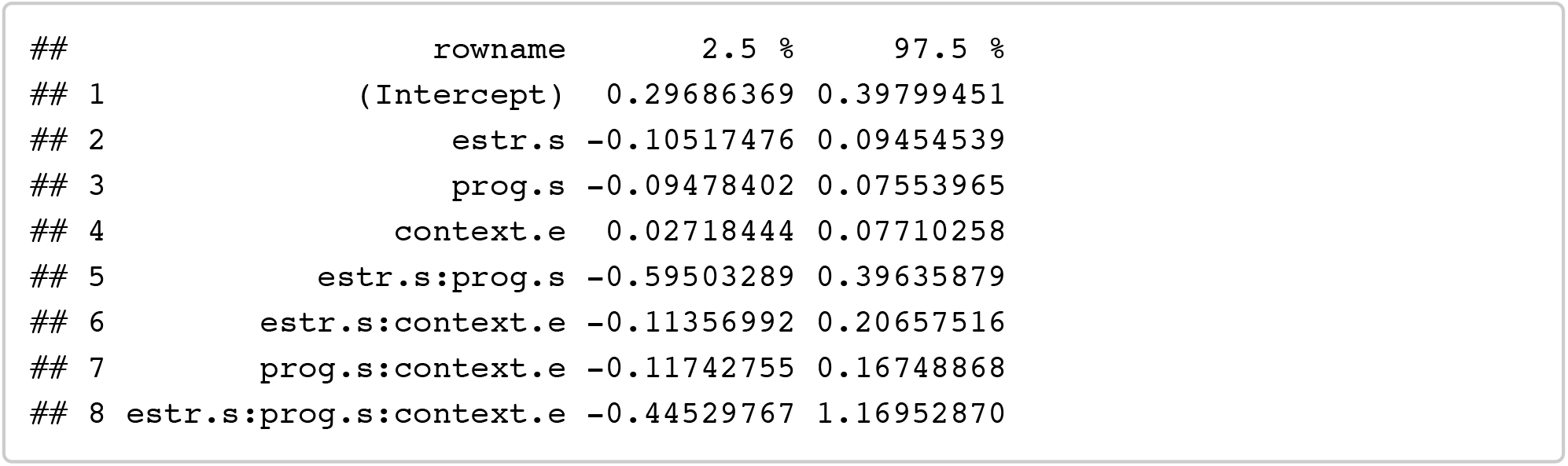

##### E + P + EPratio

Testing for effects of estradiol, progesterone, and their ratio on preferences

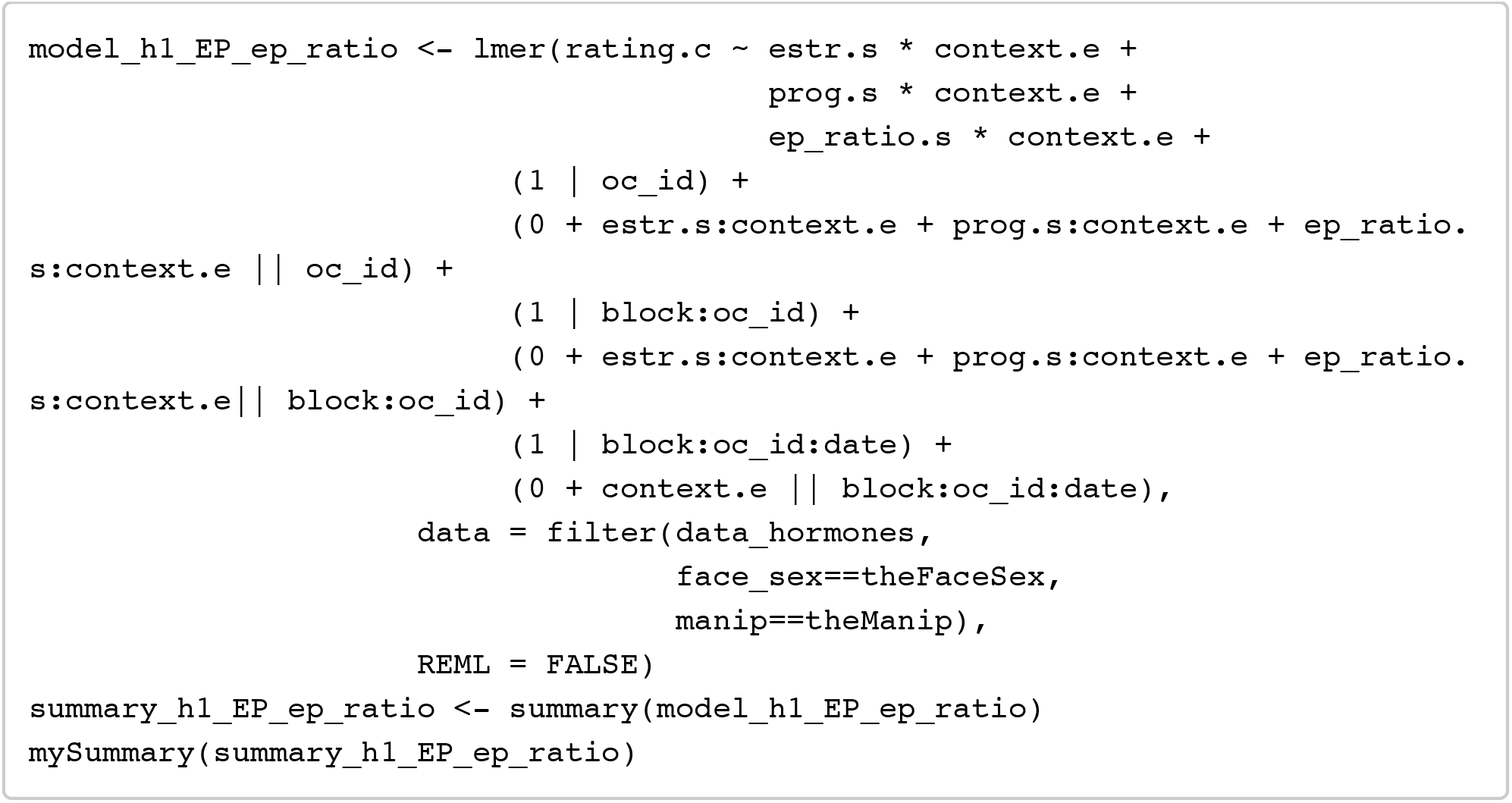

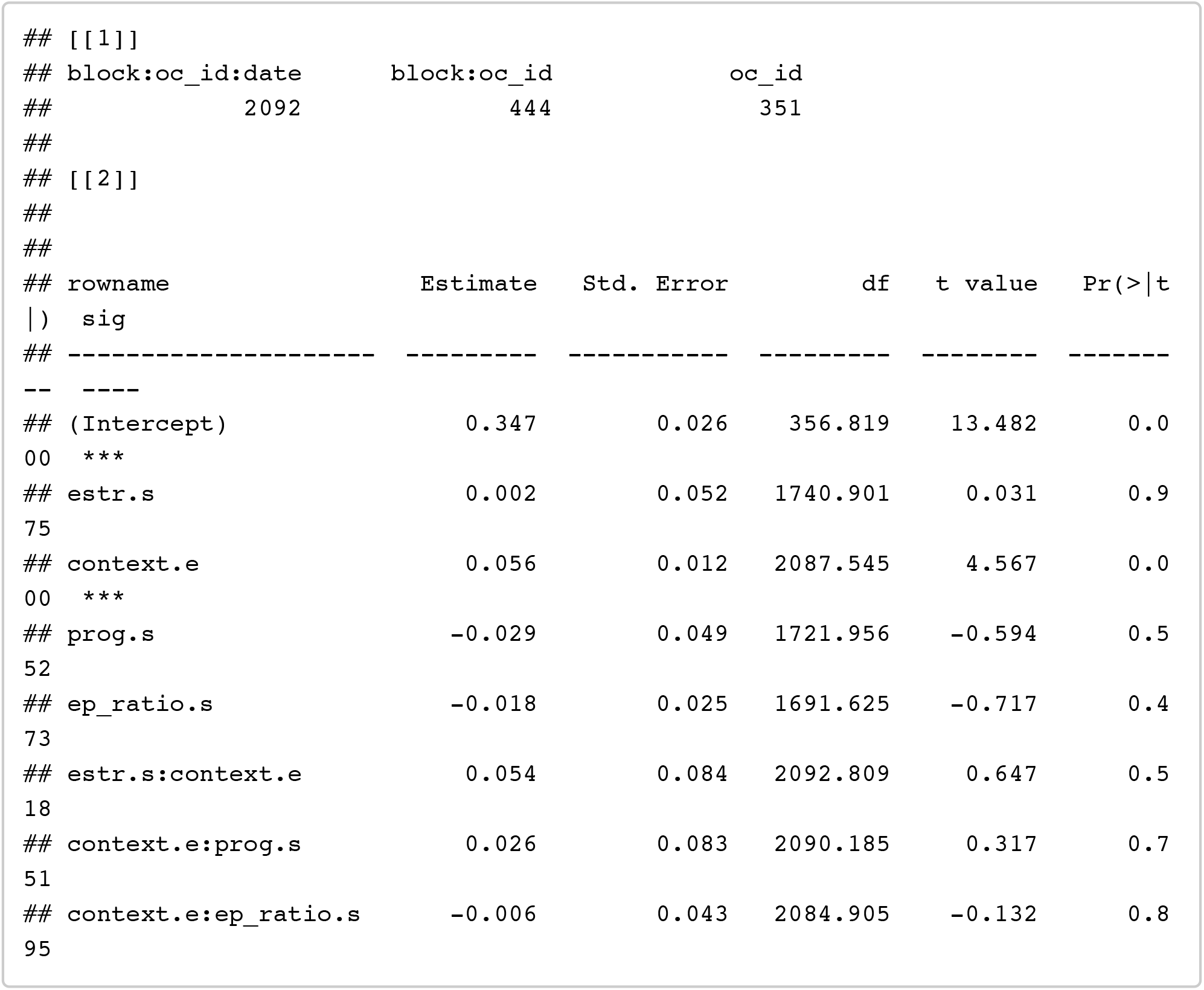

~~~
confint(model_h1_EP_ep_ratio, method = confint_method) %>% as.data.frame() %>% rownames_to_column() %>% filter(!is.na(‘2.5 %’))
~~~

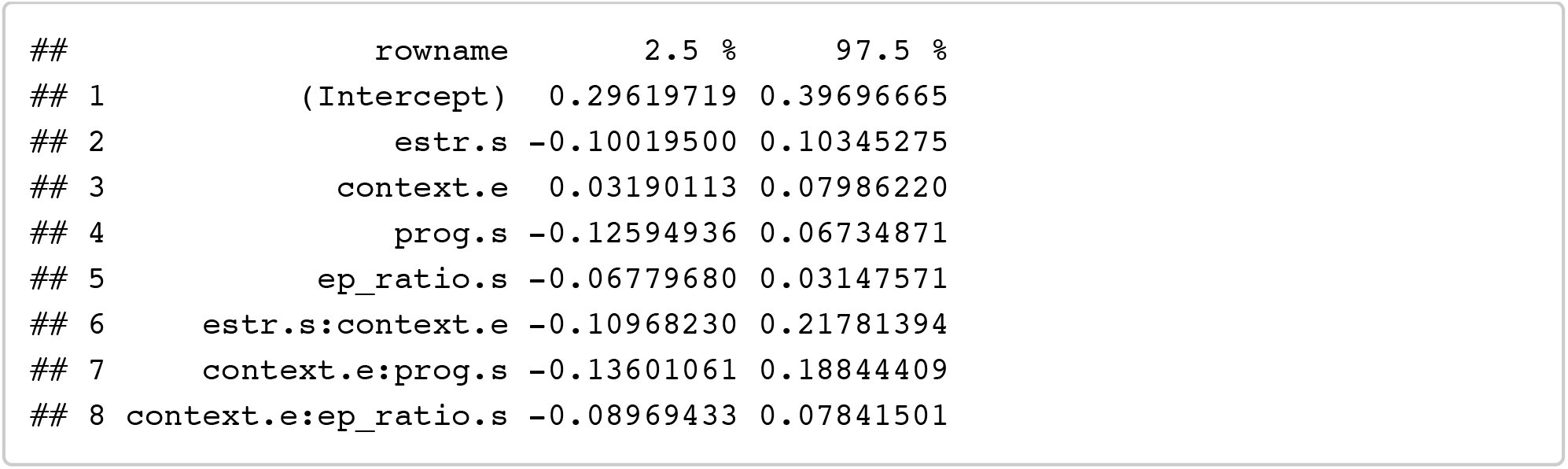

##### T + C

Testing for effects of testosterone and coritsol on preferences

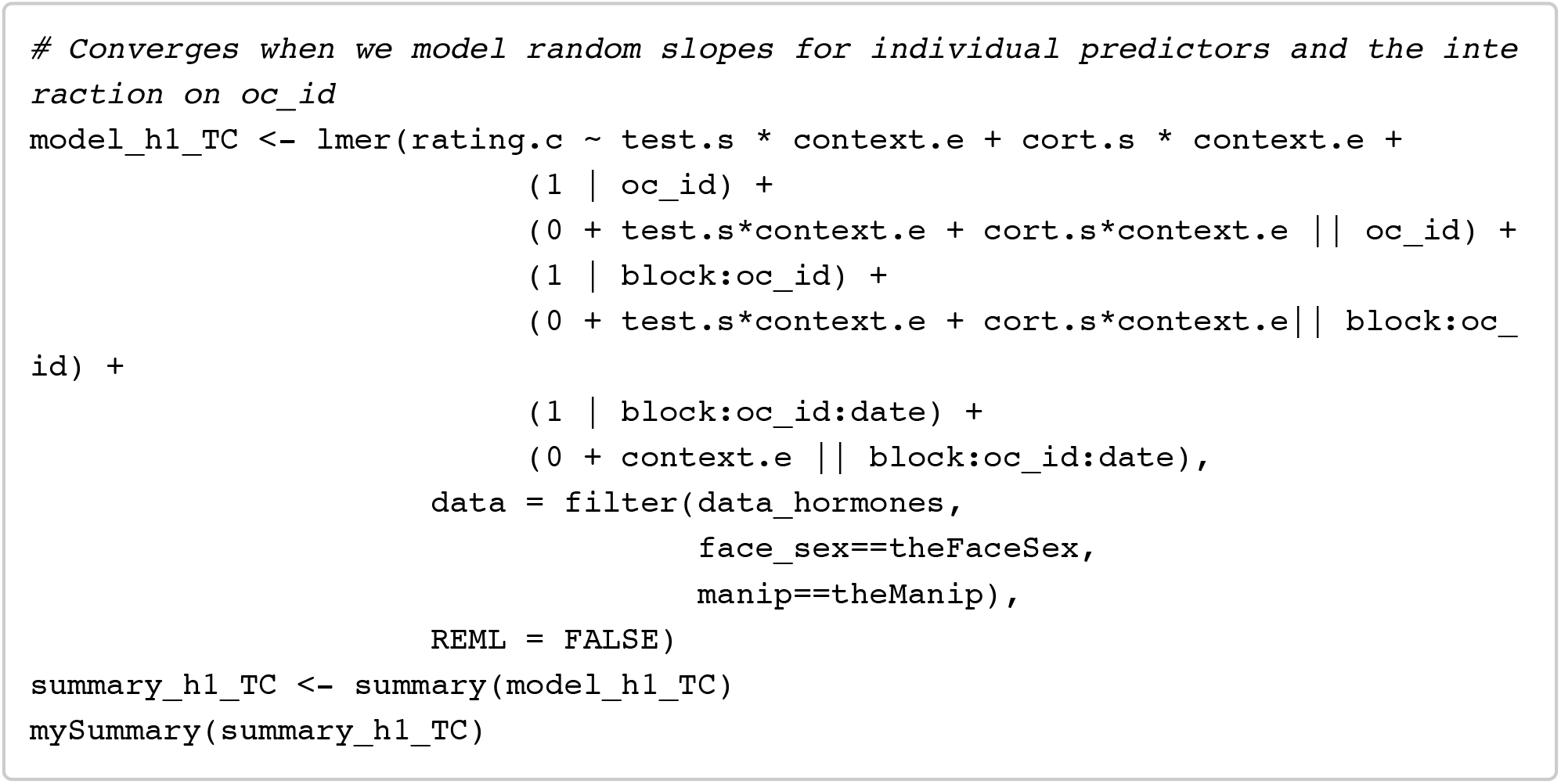

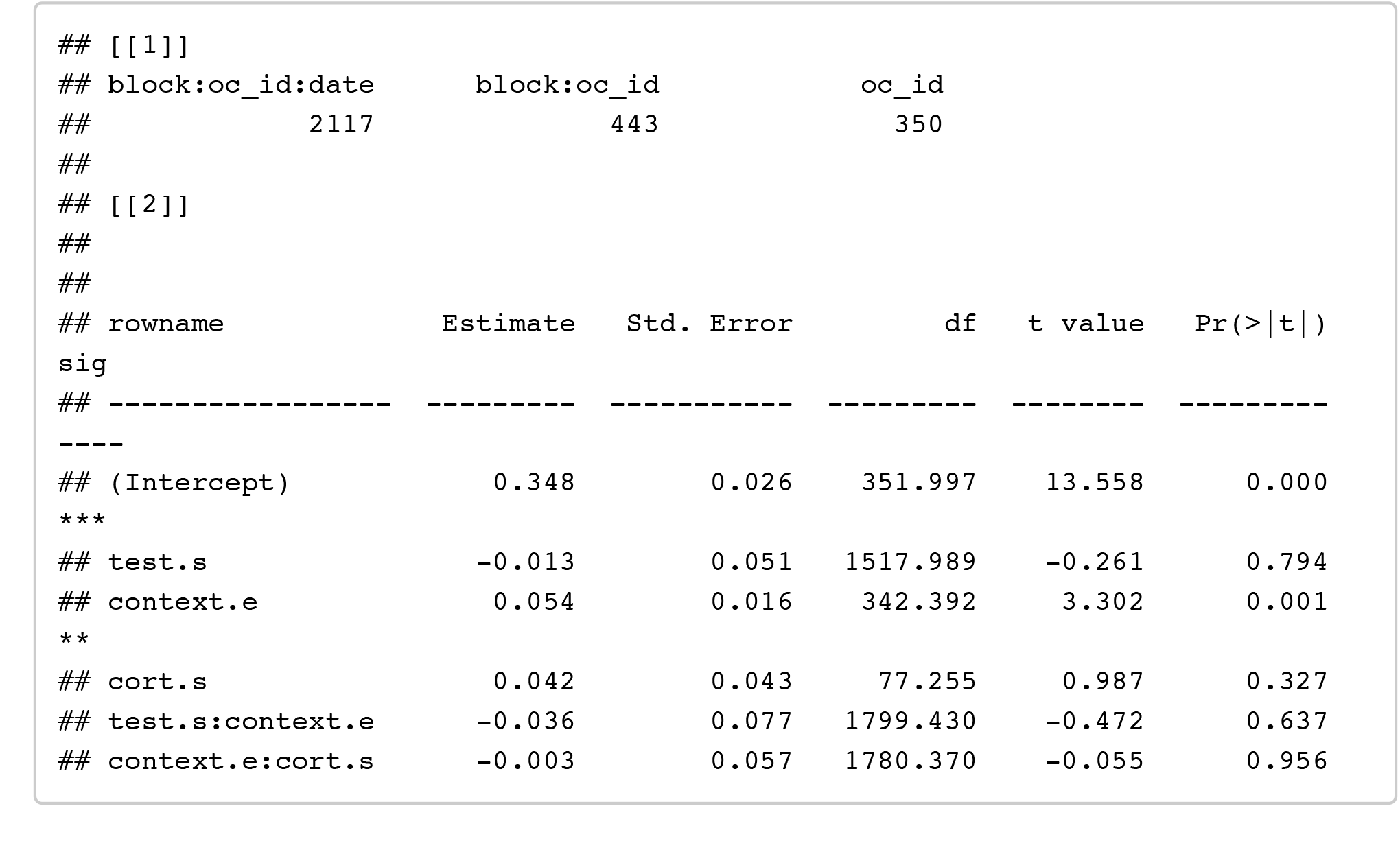

~~~
confint(model_h1_TC, method = confint_method) %>% as.data.frame() %>% rownames _to_column() %>% filter(!is.na(‘2.5 %’))
~~~

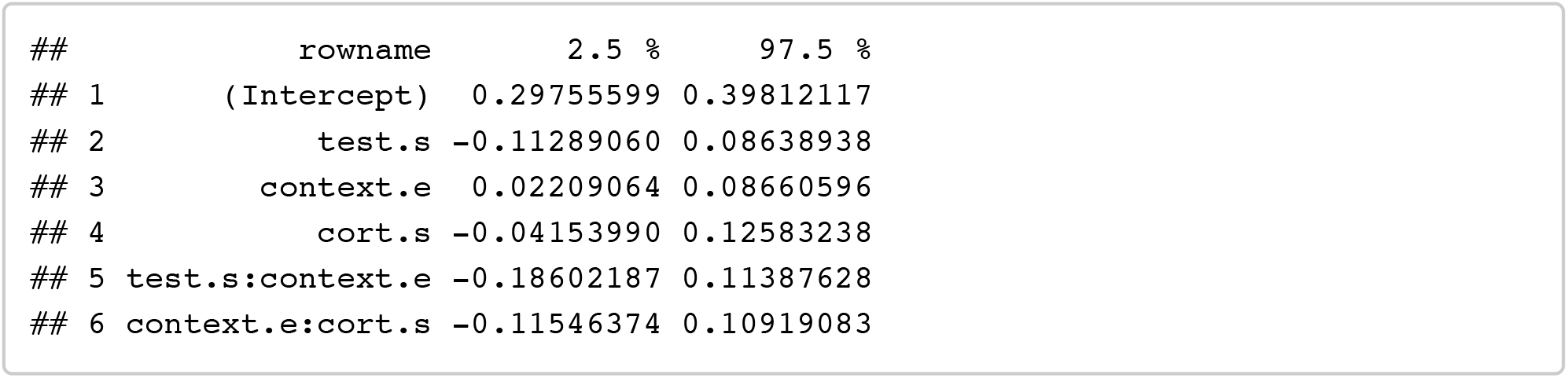

##### T + C + T^*^C

Testing for effects of testosterone and coritsol plus their interaction on preferences

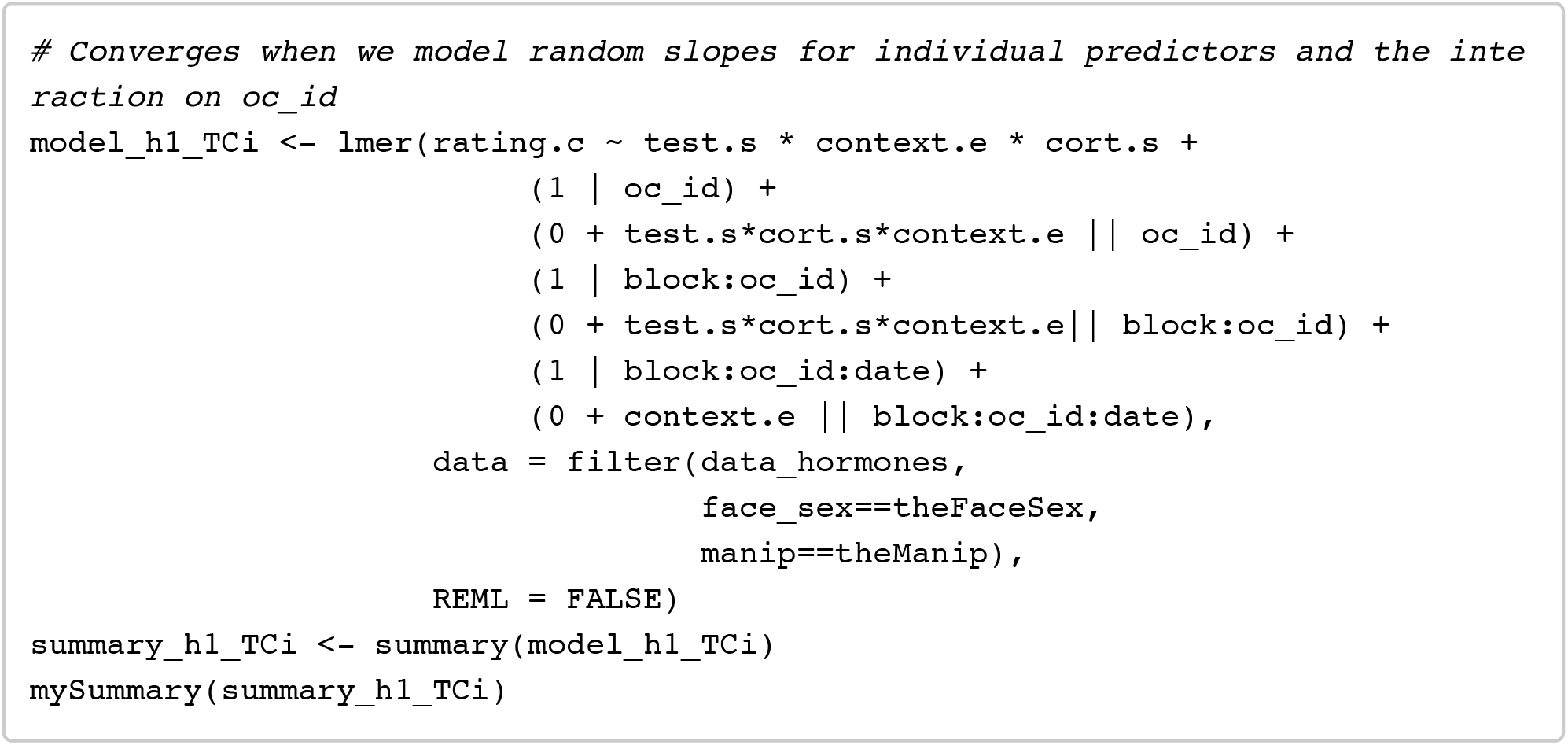

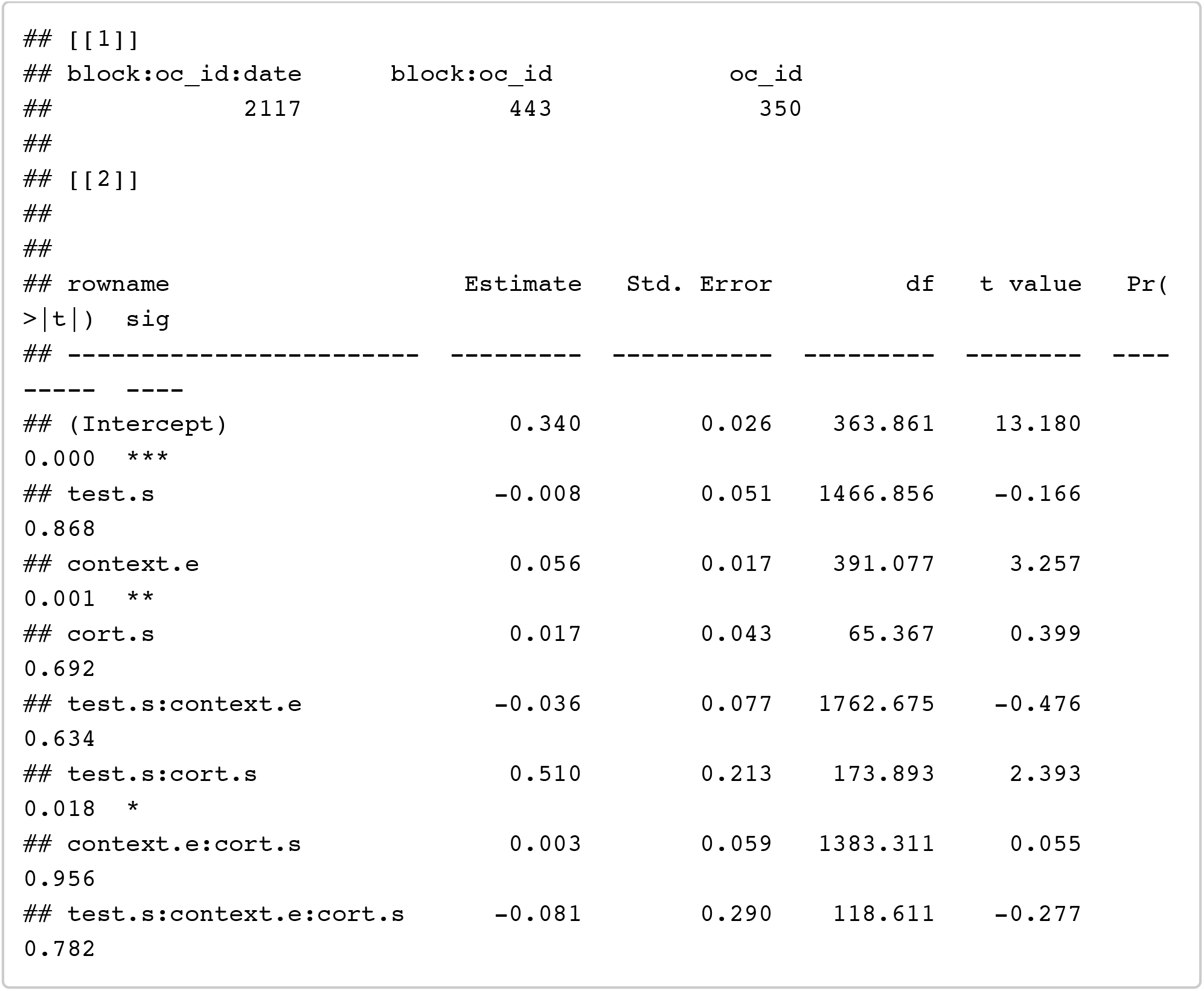

~~~
confint(model_h1_TCi, method = confint_method) %>% as.data.frame() %>% rowname s_to_column() %>% filter(!is.na(‘2.5 %’))
~~~

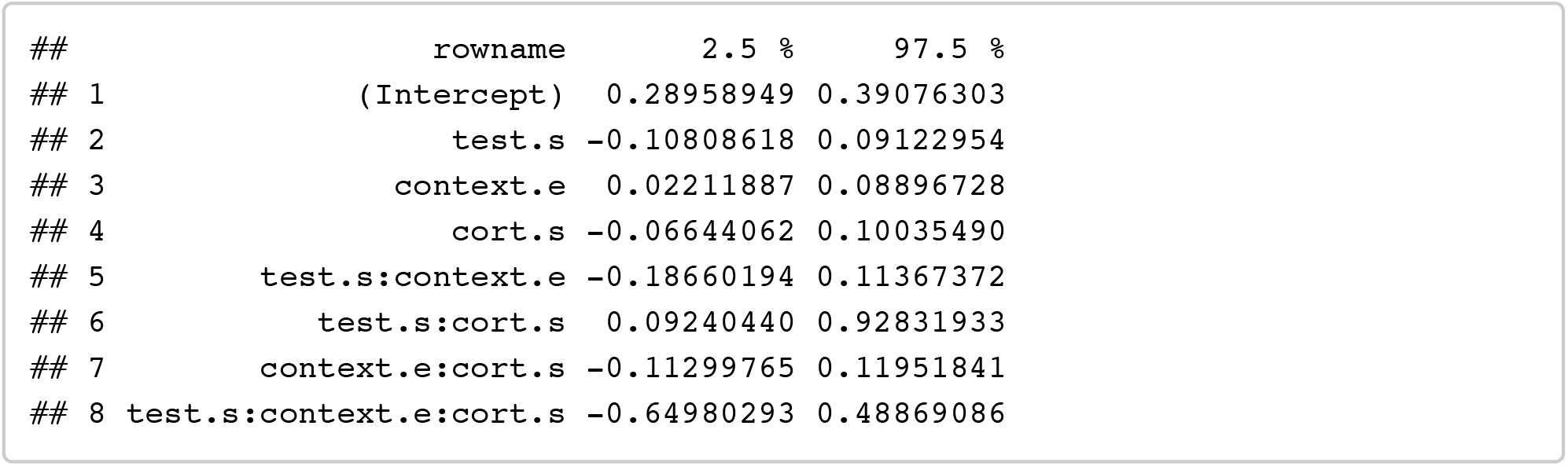

#### Analyses H1s: Hormones (+ session order)

##### E + P + E^*^P: (+ session order)

Testing for effects of estradiol, progesterone, and their interaction on preferences

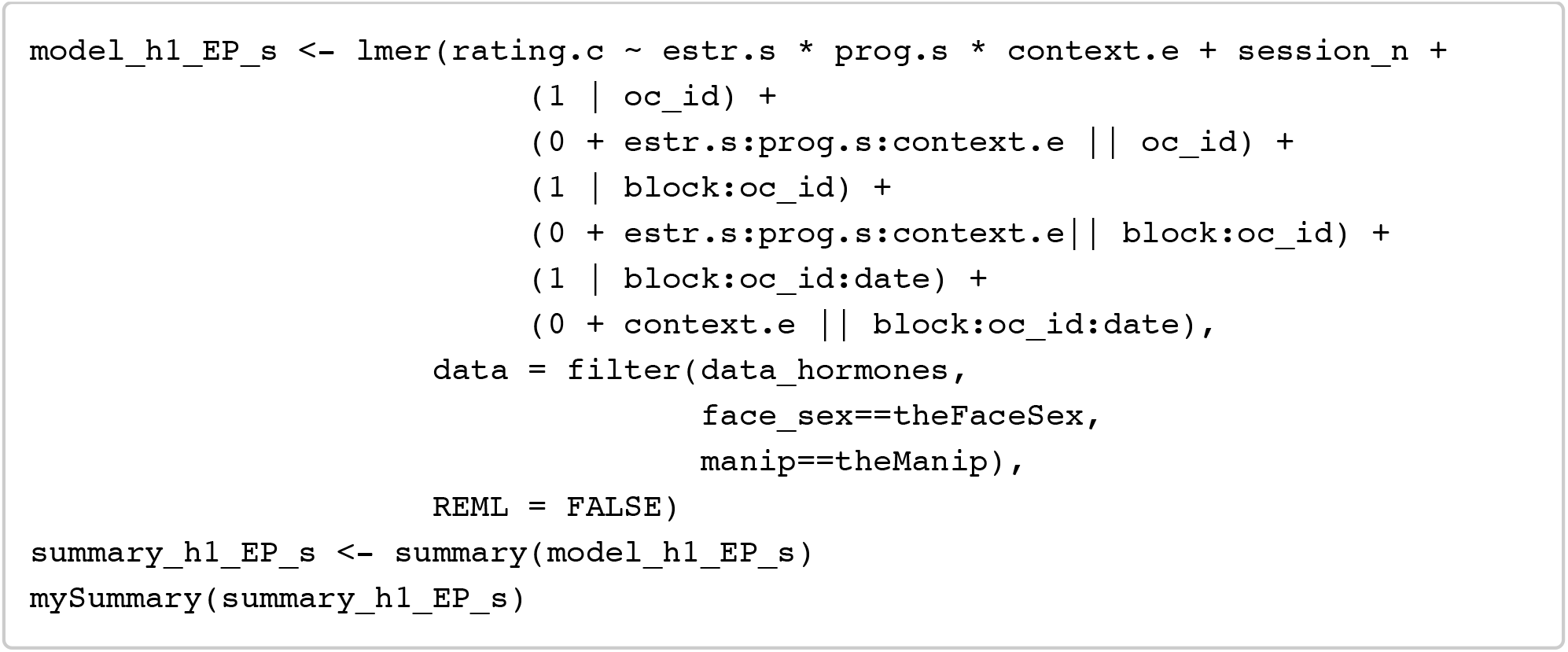

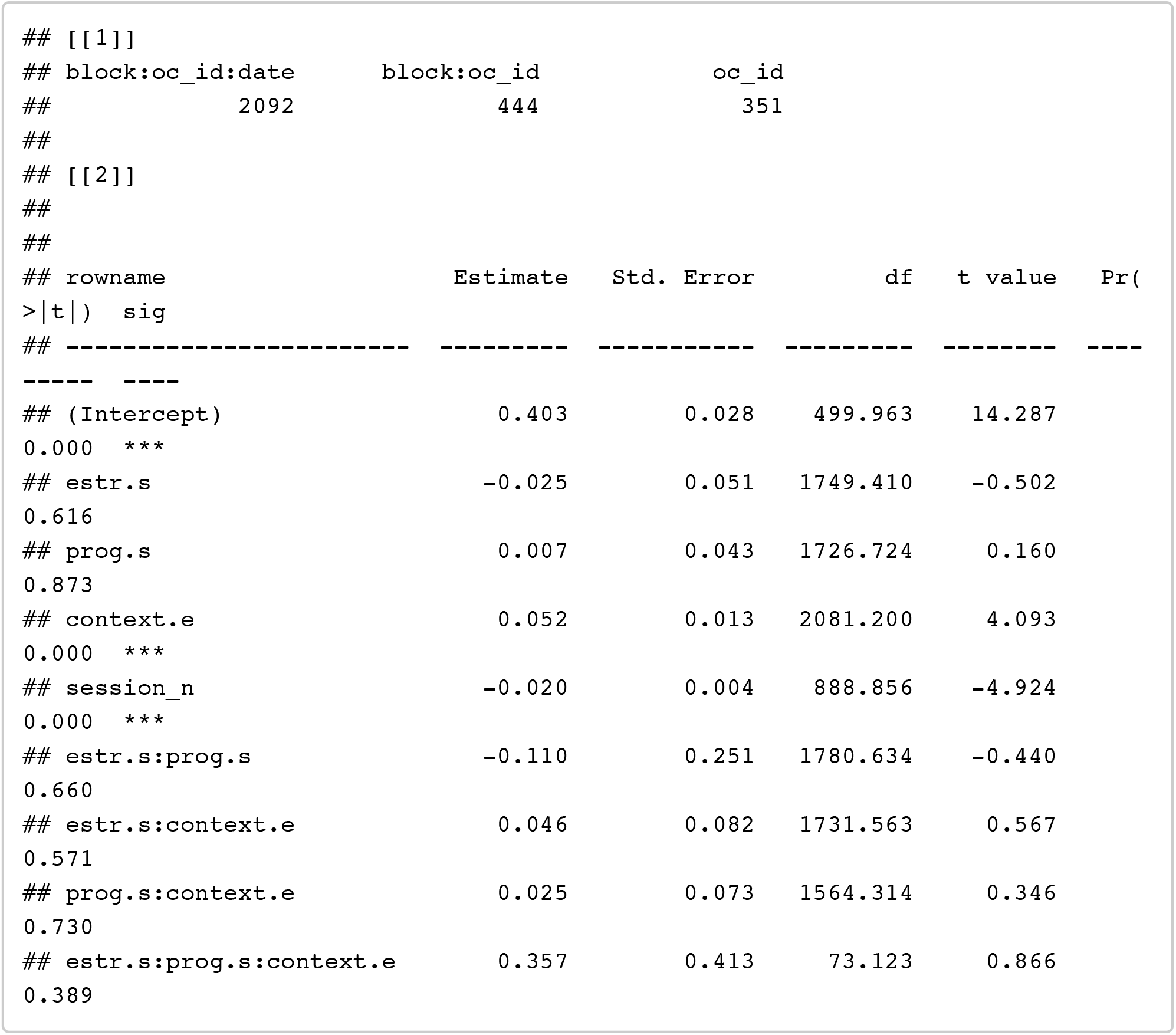

~~~
confint(model_h1_EP_s, method = confint_method) %>% as.data.frame() %>% rownam es_to_column() %>% filter(!is.na(‘2.5 %’))
~~~

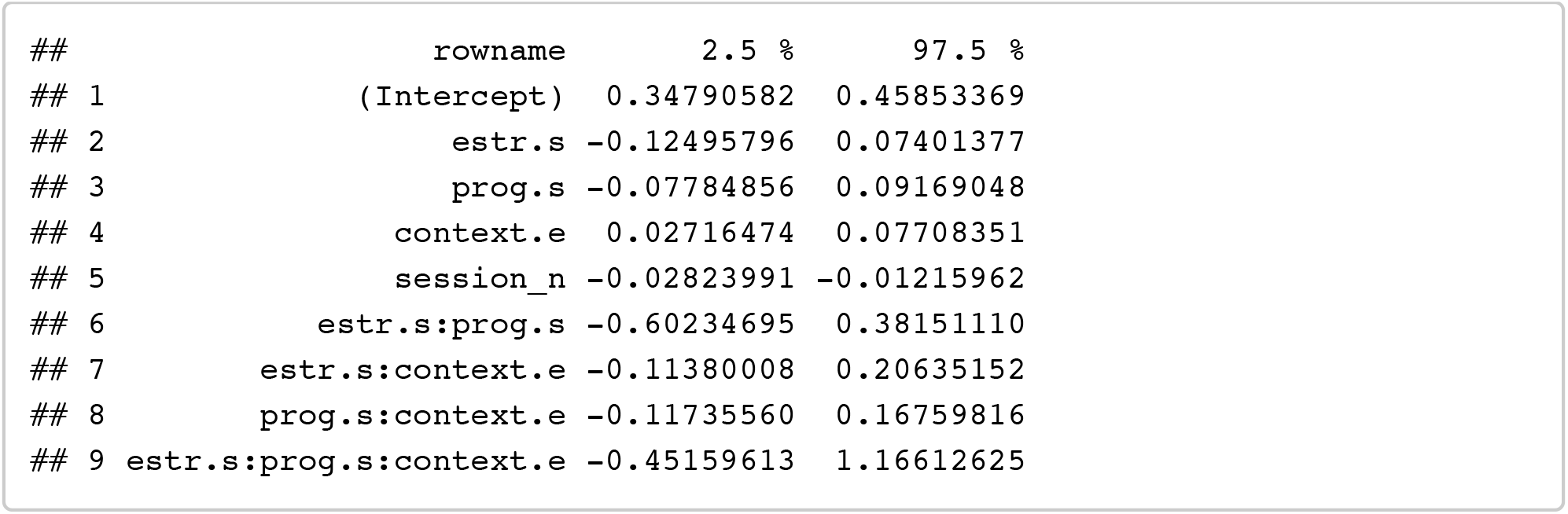

##### E + P + EPratio: (+ session order)

Testing for effects of estradiol, progesterone, and their ratio on preferences

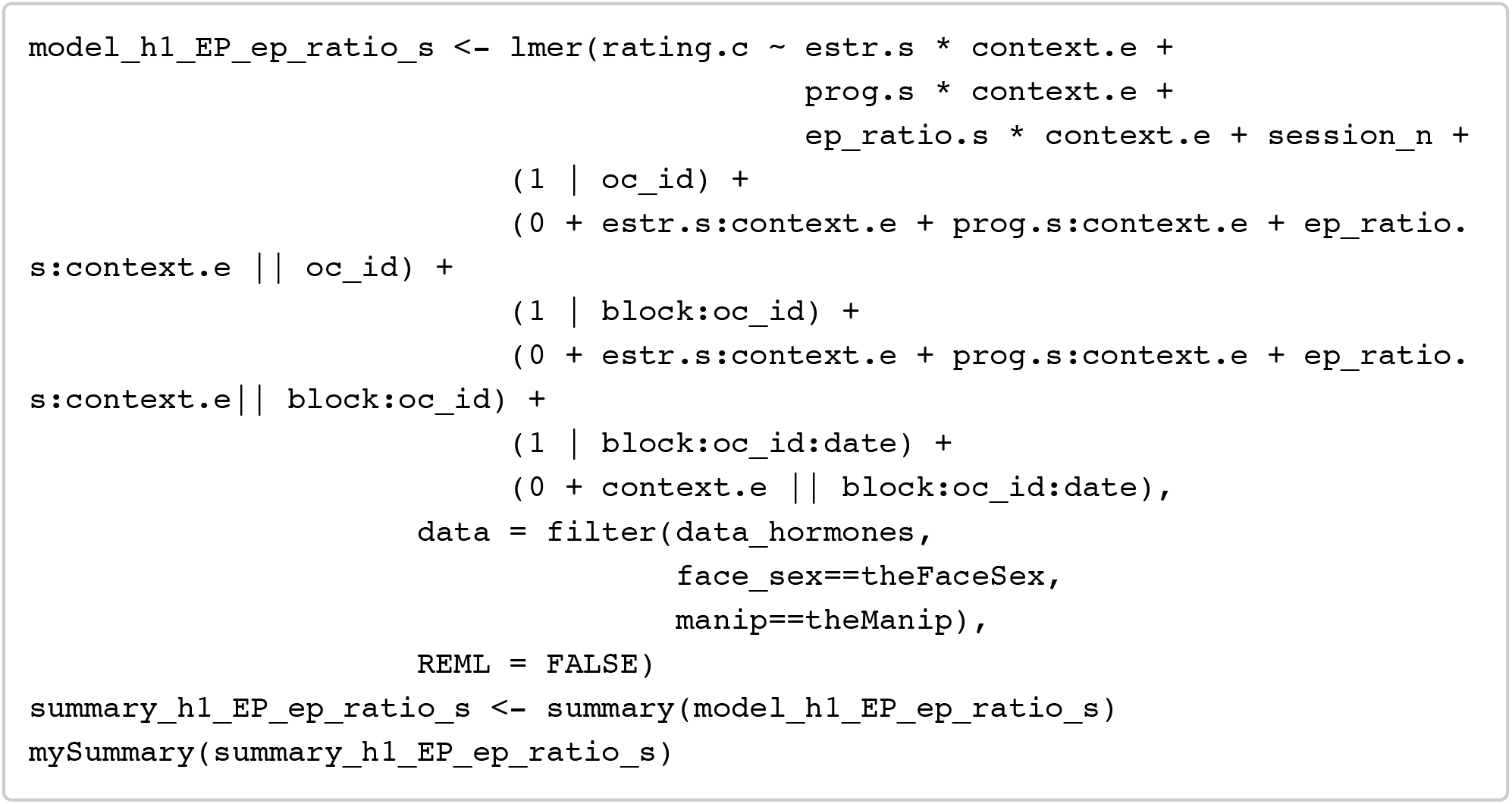

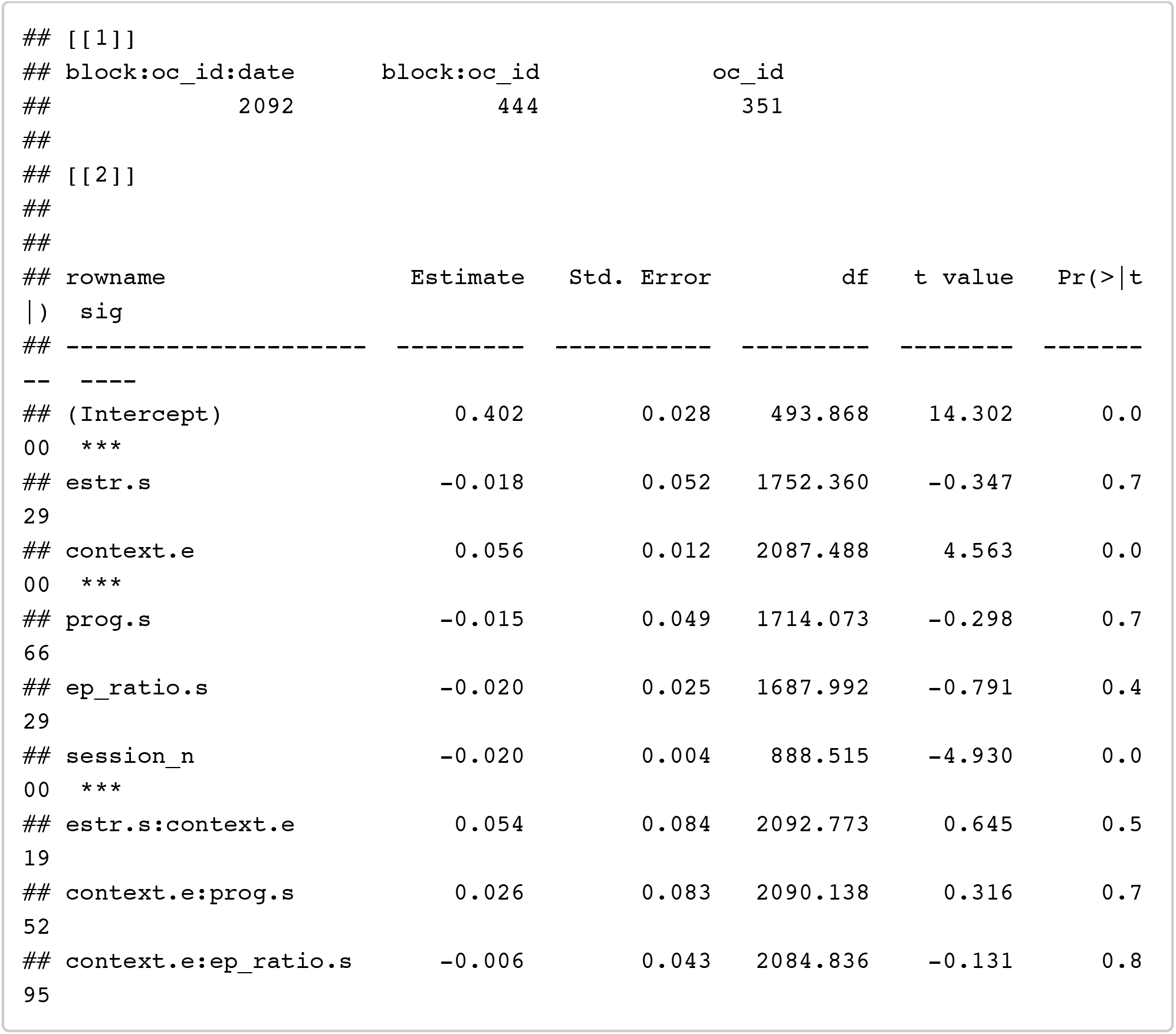

~~~
confint(model_h1_EP_ep_ratio_s, method = confint_method) %>% as.data.frame() % >% rownames_to_column() %>% filter(!is.na(‘2.5 %’))
~~~

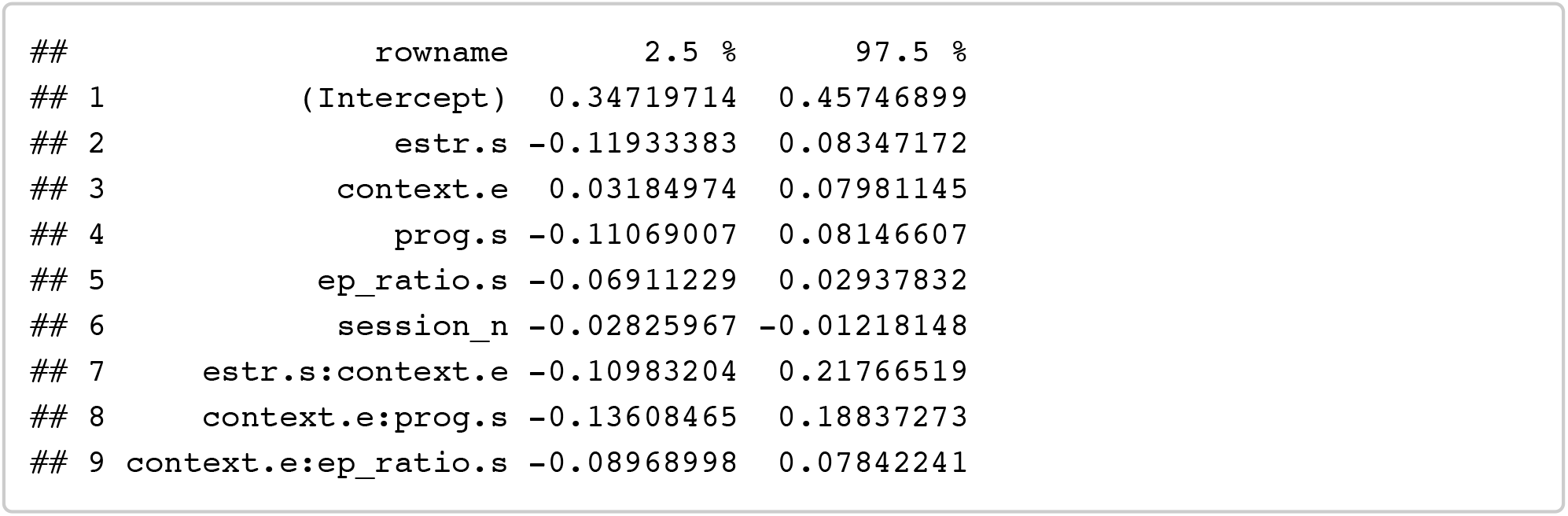

##### T + C: (+ session order)

Testing for effects of testosterone and coritsol on preferences

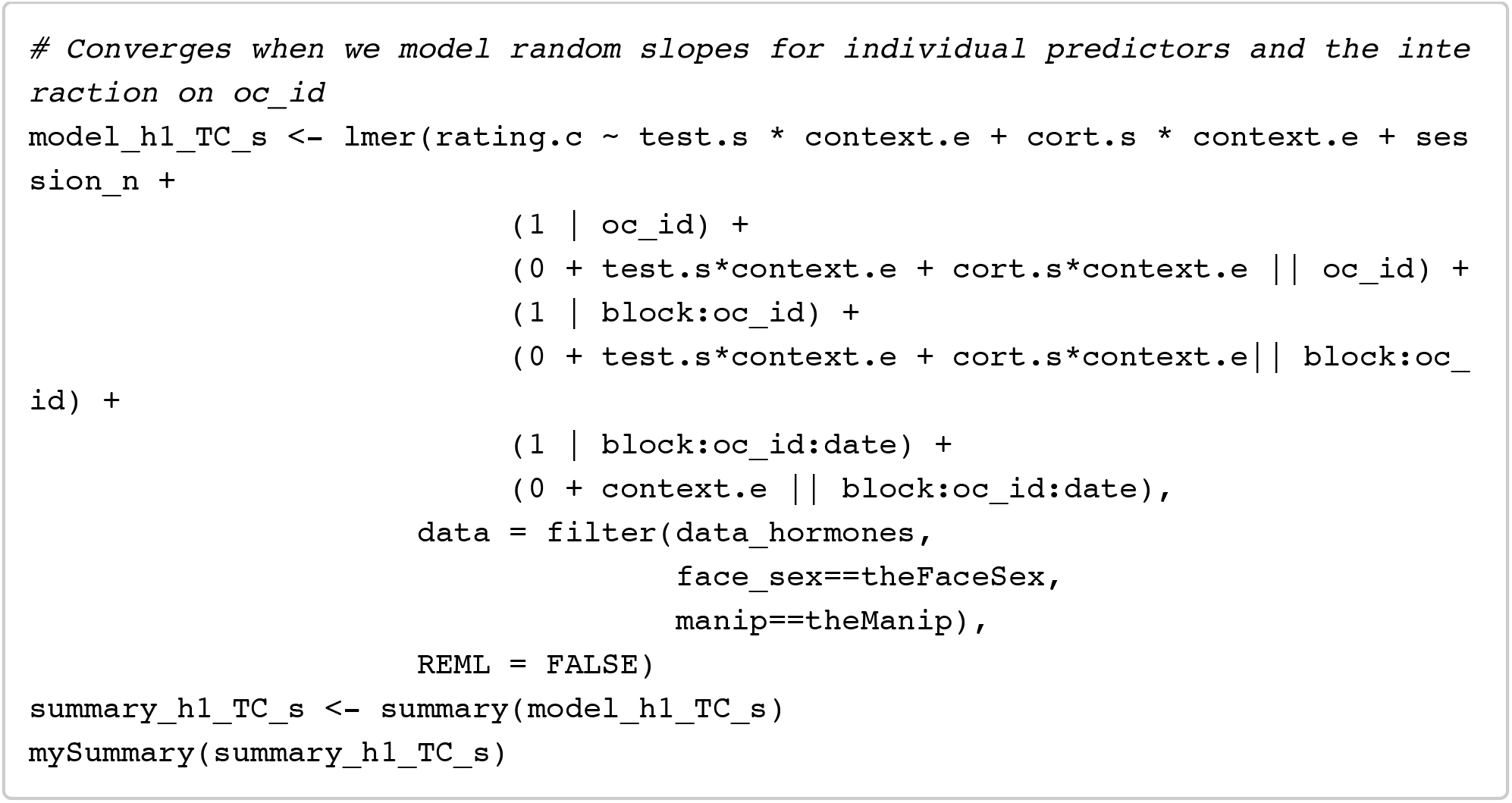

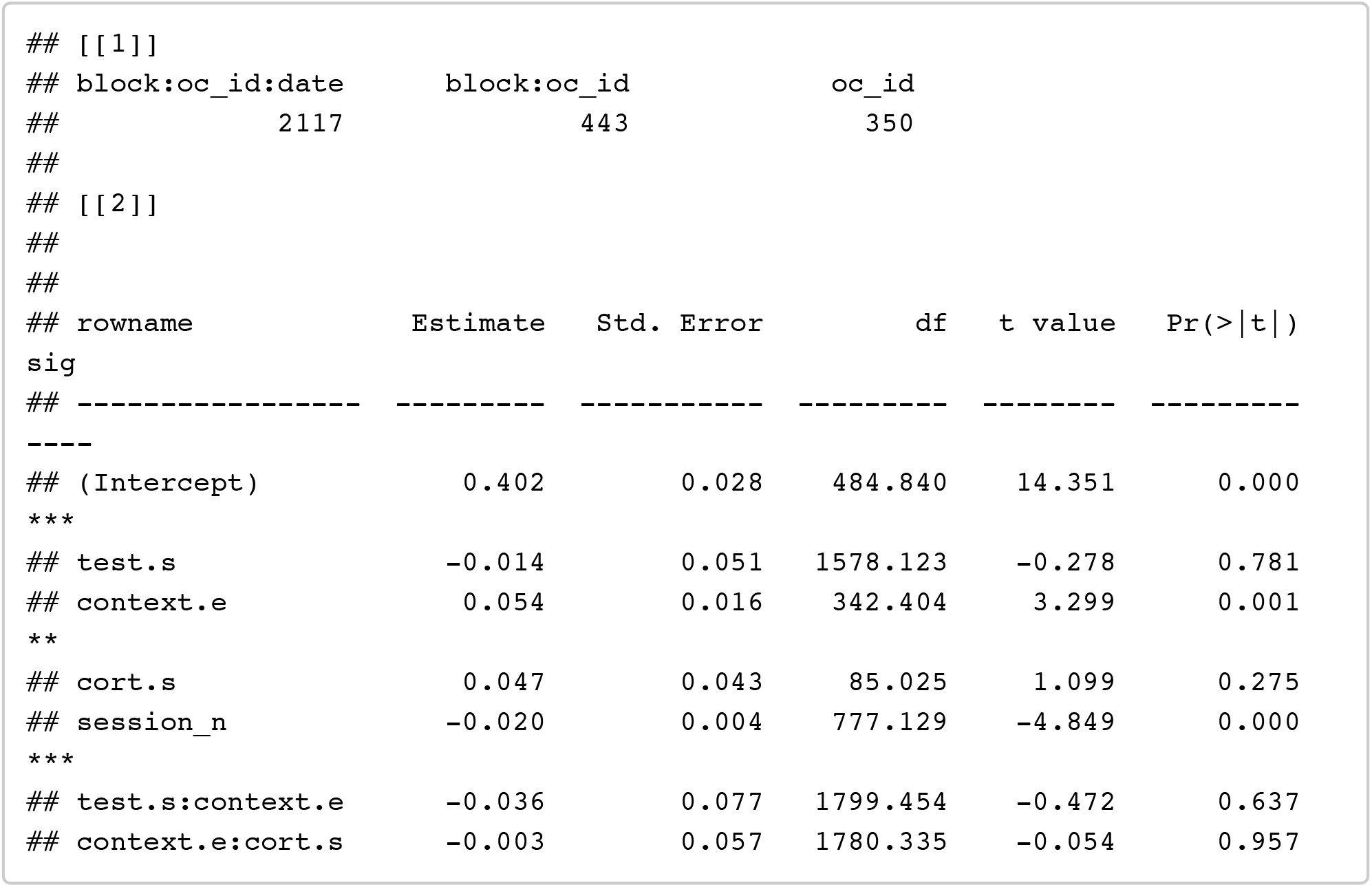

~~~
confint(model_h1_TC_s, method = confint_method) %>% as.data.frame() %>% rownam es_to_column() %>% filter(!is.na(‘2.5 %’))
~~~

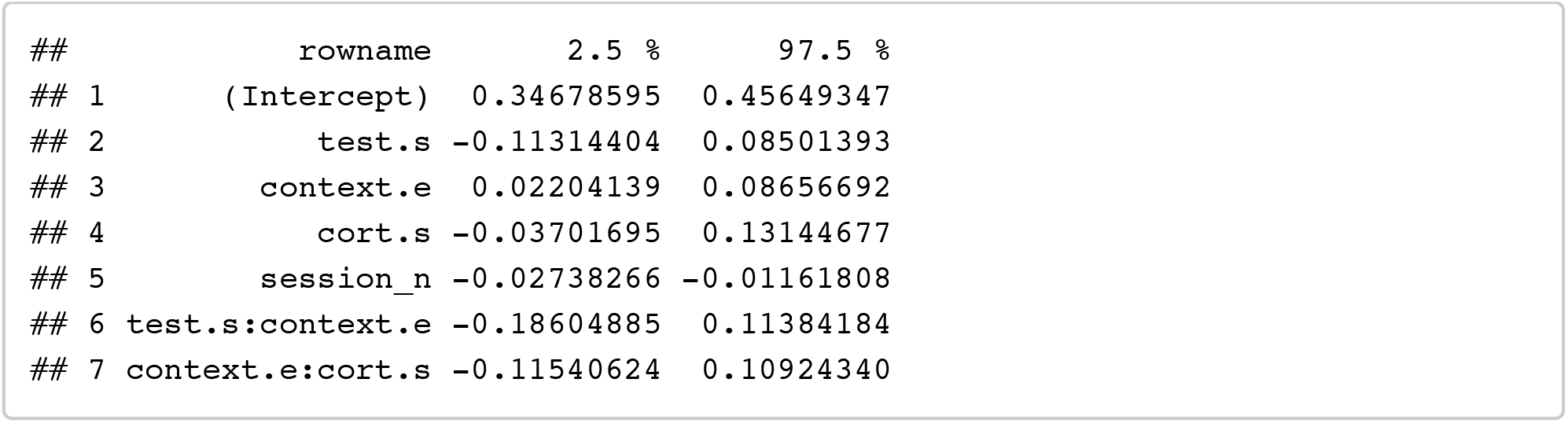

##### T + C + T^*^C: (+ session order)

Testing for effects of testosterone and coritsol plus their interaction on preferences

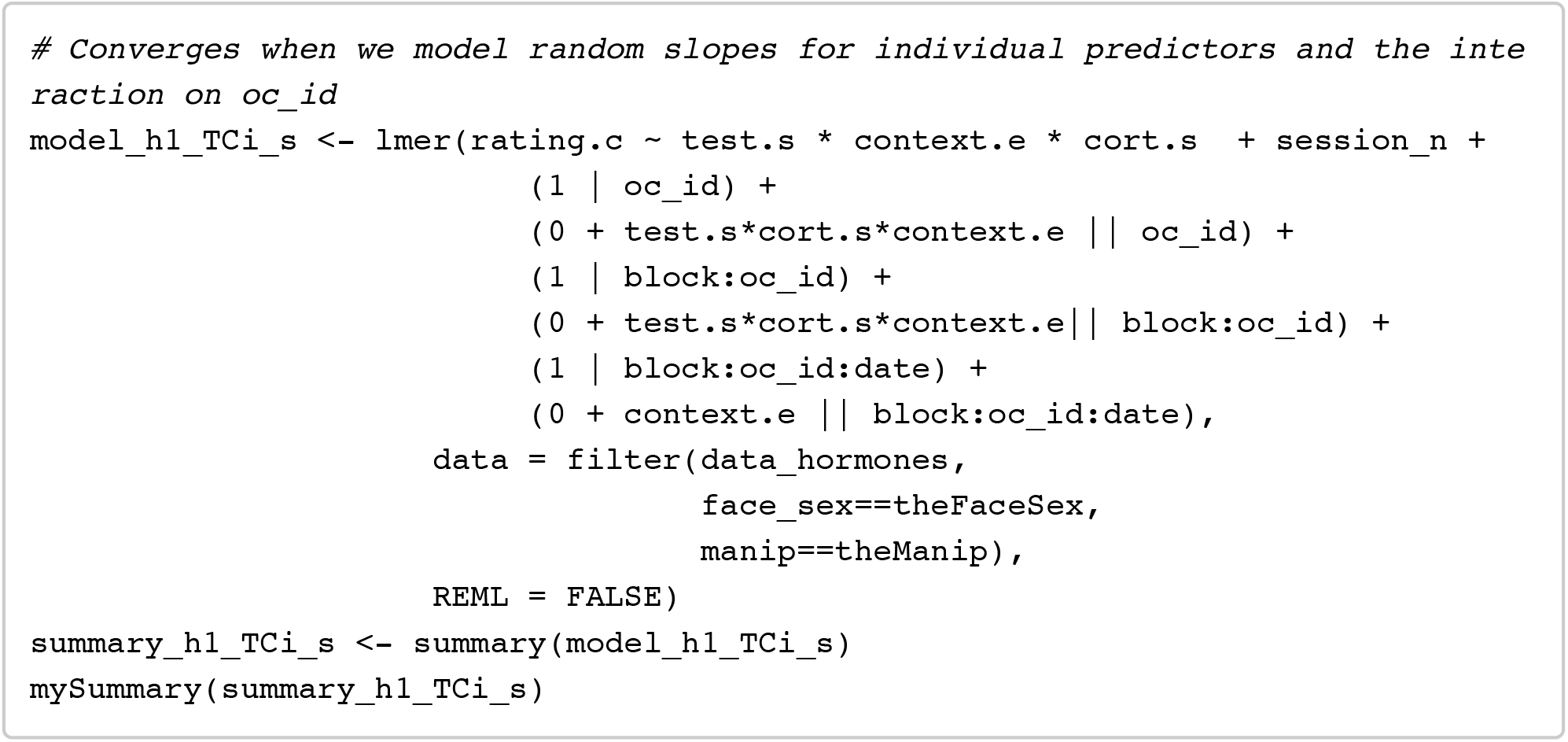

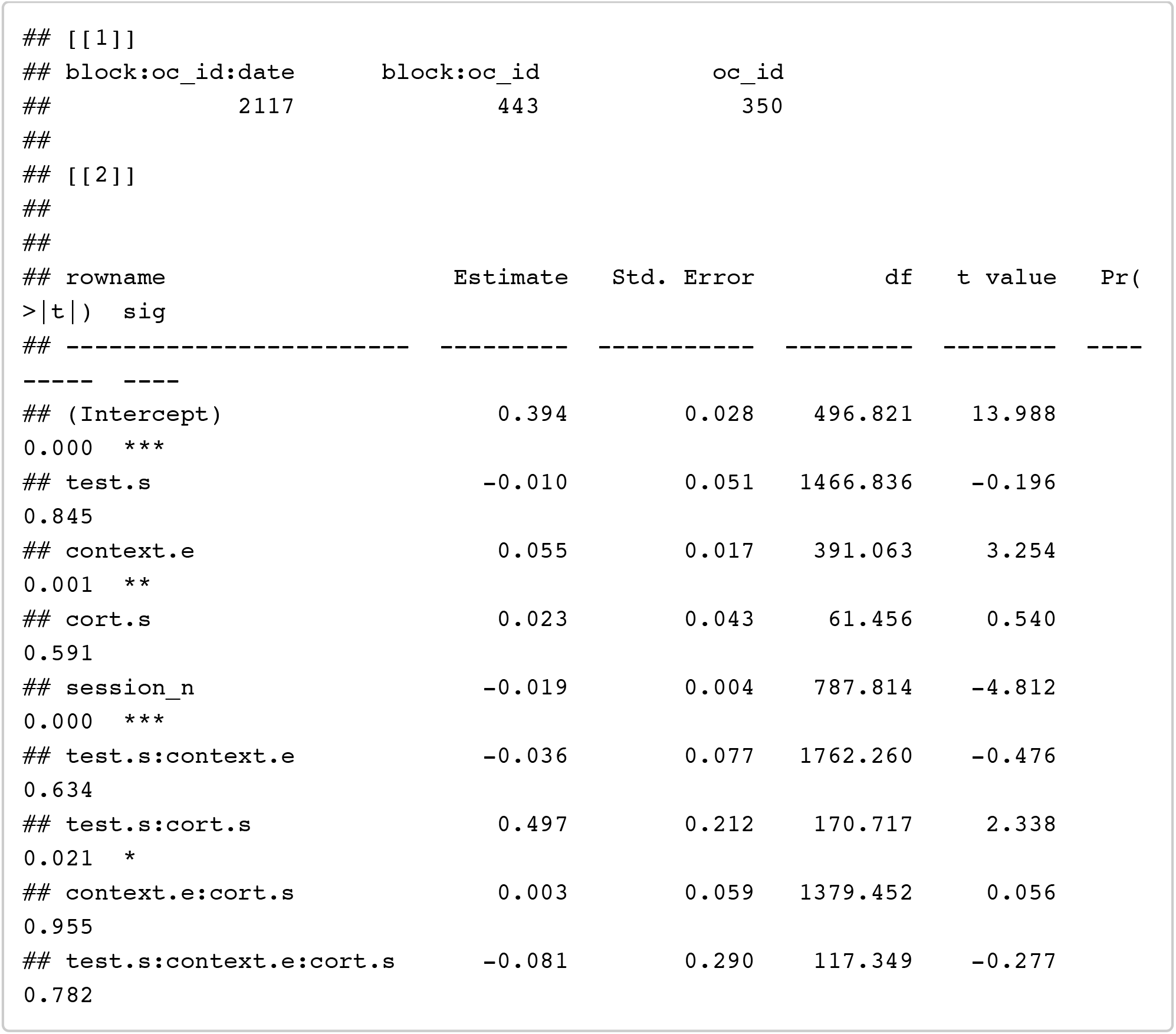

~~~
confint(model_h1_TCi_s, method = confint_method) %>% as.data.frame() %>% rowna mes_to_column() %>% filter(!is.na(‘2.5 %’))
~~~

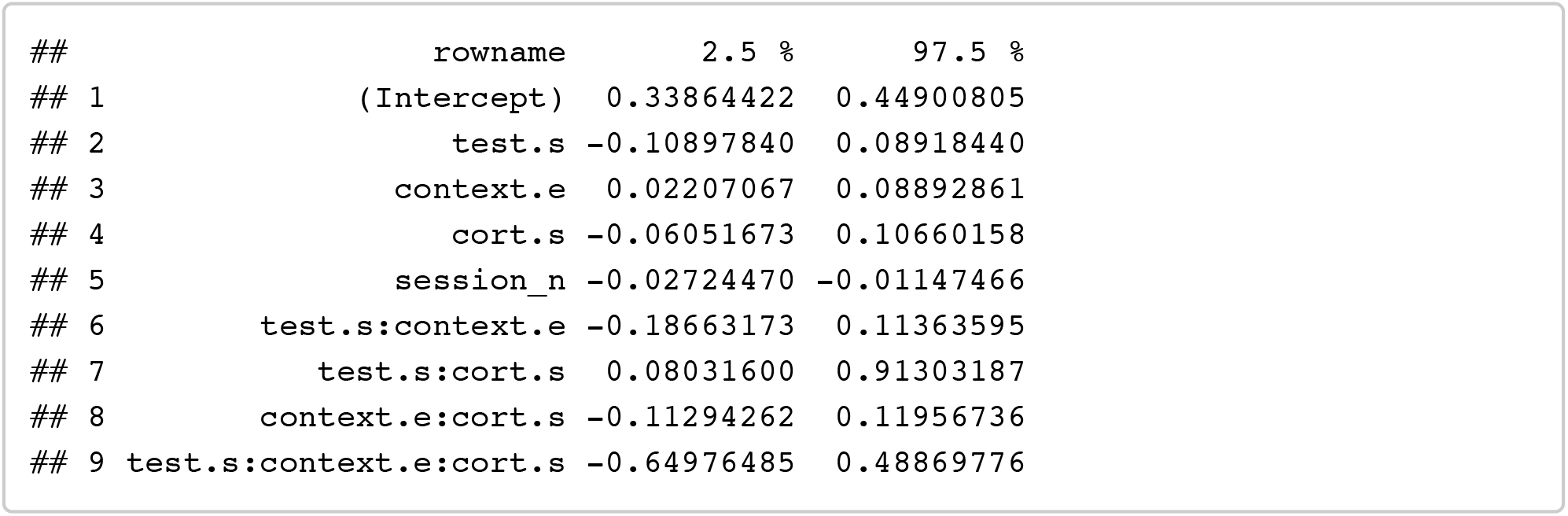

#### Descriptive stats: data_hormones_partner

~~~
data_hormones_partner %>%
  filter(face_sex==theFaceSex, manip==theManip) %>%
  group_by(oc_id, partner.e) %>%
  summarise(overall_rating.c = mean(rating.c)) %>%
  group_by(partner.e) %>%
  summarise(
    n= n_distinct(oc_id),
    mean_dv = mean(overall_rating.c),
    sd_dv = sd(overall_rating.c),
    se_dv = se(overall_rating.c)
)
~~~

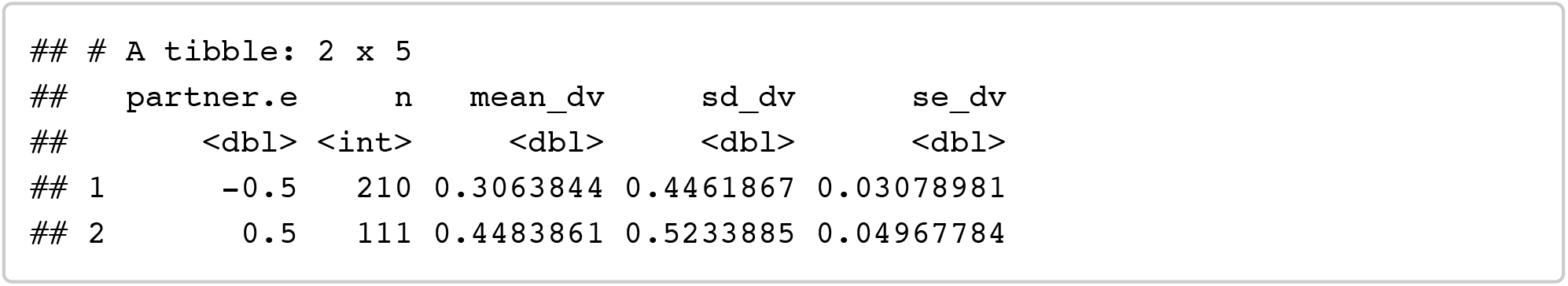

#### Analyses H1 p: Hormones (+ partnership status)

##### E + P + E^*^P: (+ partnership status)

Testing for effects of estradiol, progesterone, and their interaction on preferences

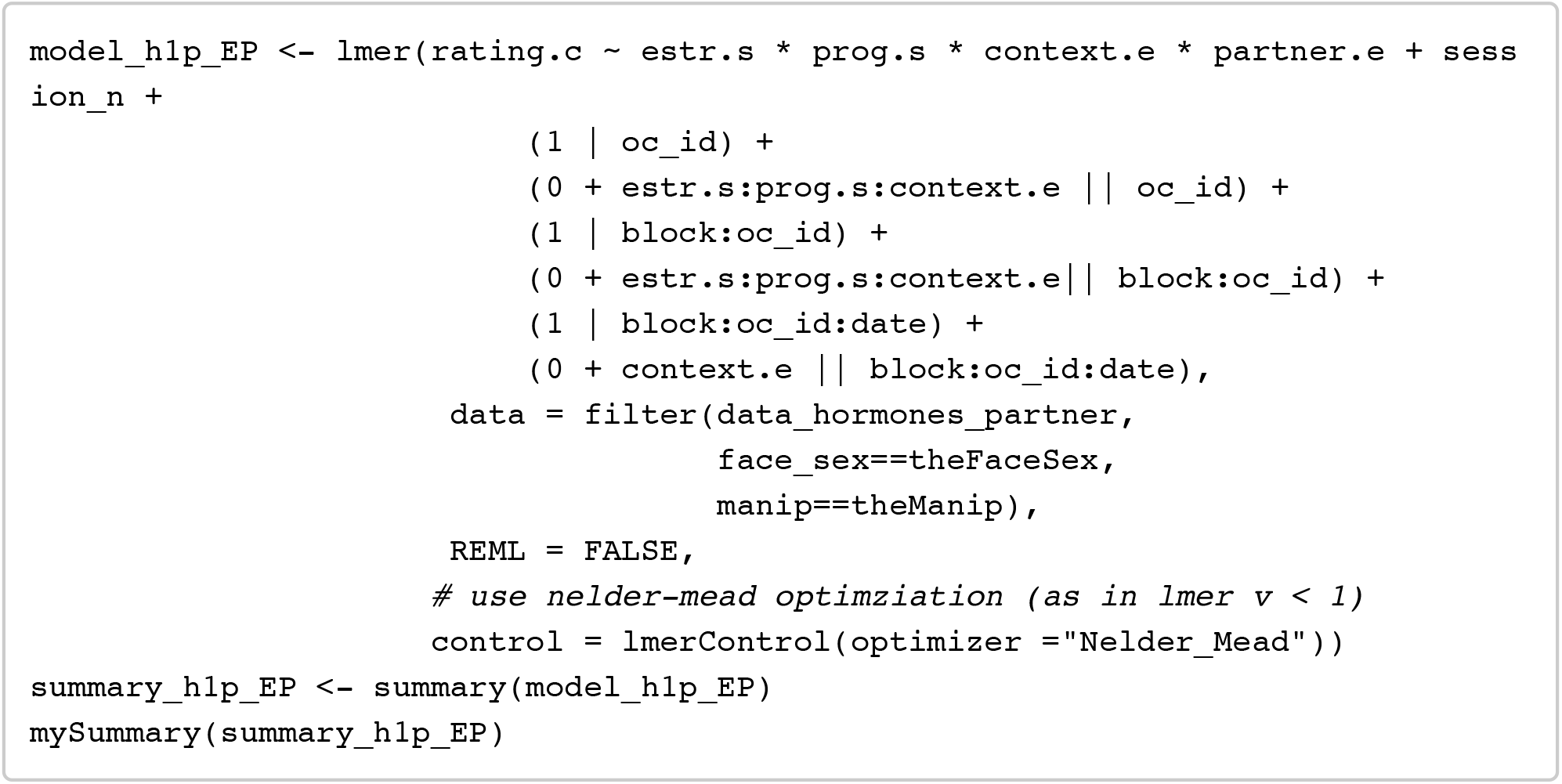

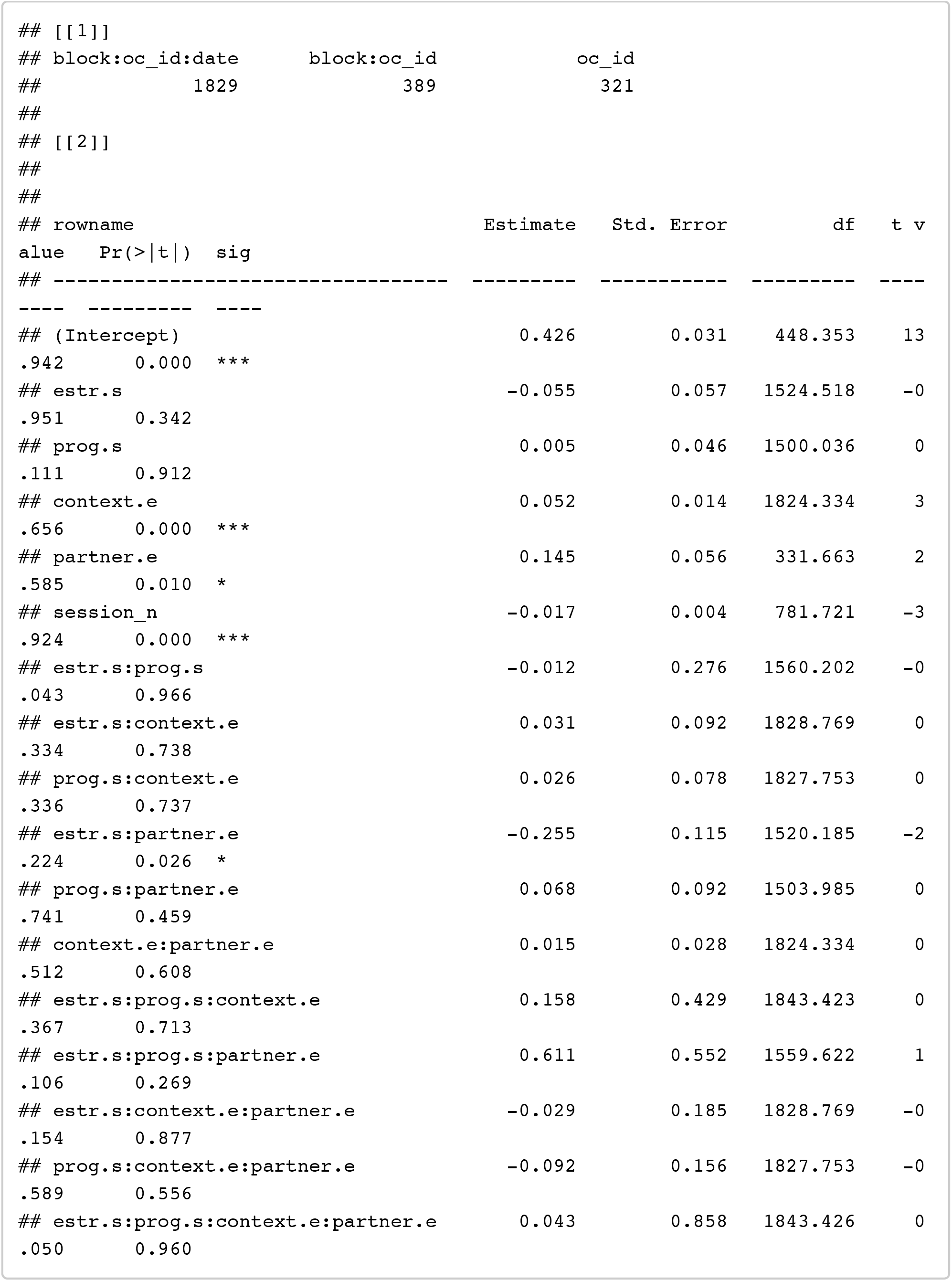

~~~
confint(model_h1p_EP, method = confint_method) %>% as.data.frame() %>% rowname s_to_column() %>% filter(!is.na(‘2.5 %’))
~~~

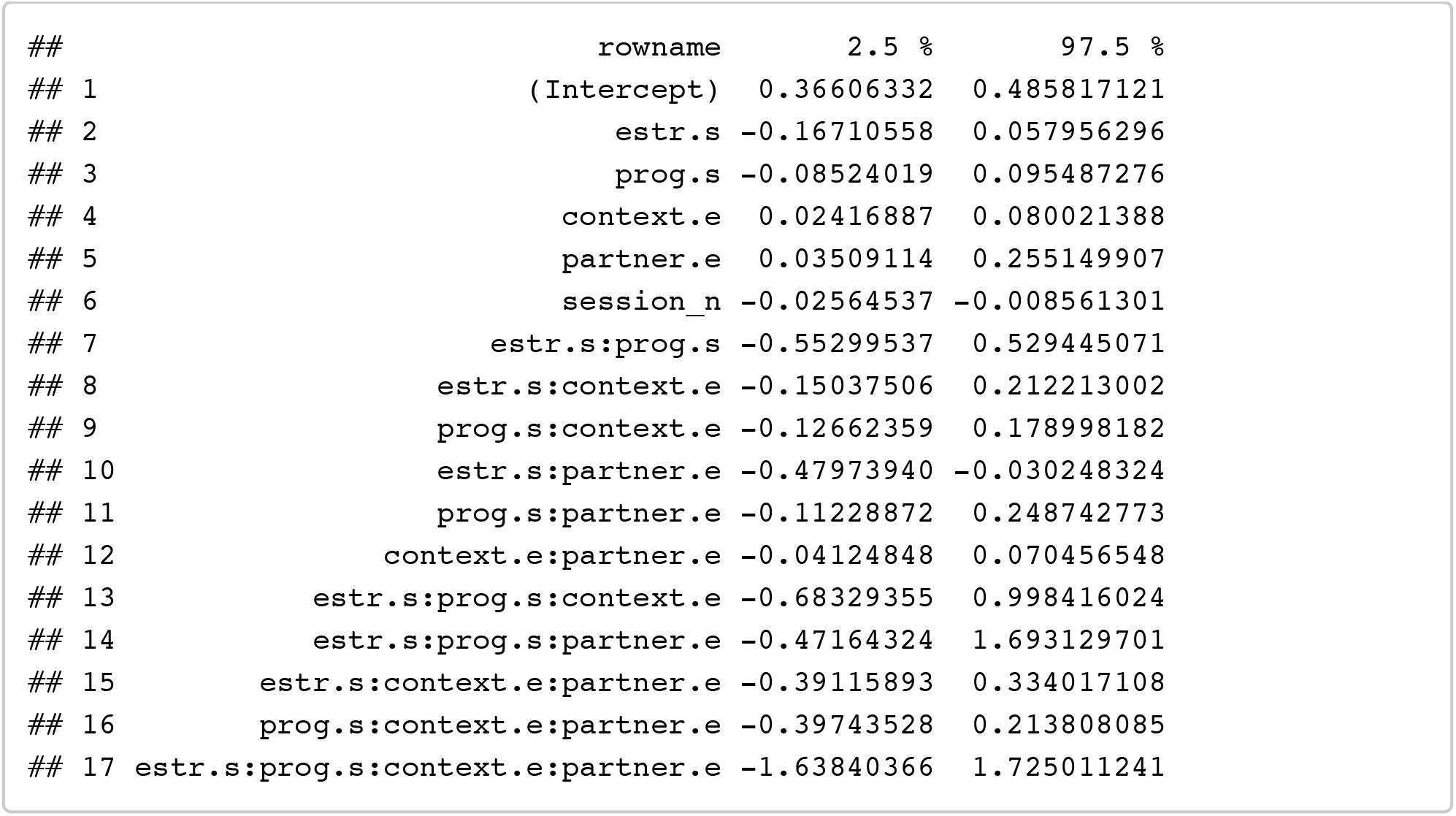

##### E + P + E*P: (single women only to interpret interaction)

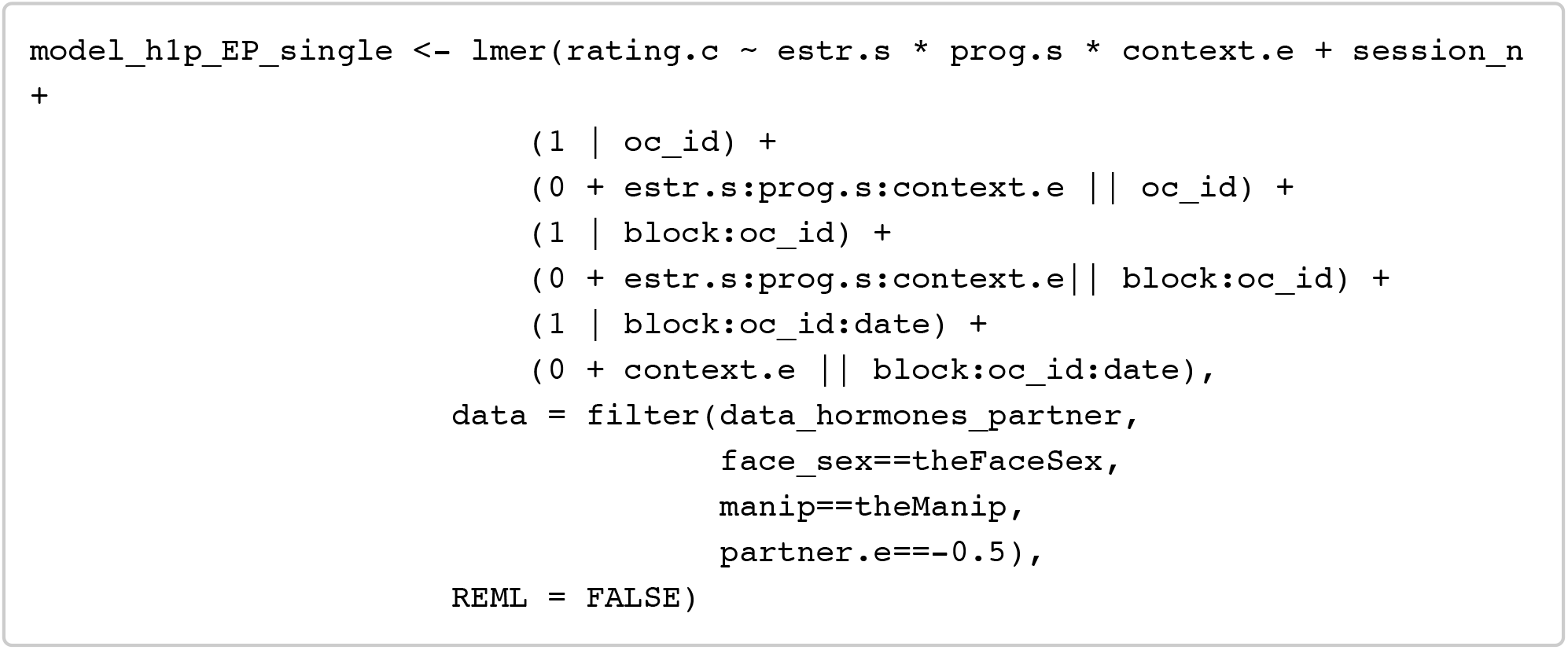

~~~
## Warning in checkConv(attr(opt, “derivs”), opt$par, ctrl = control$checkConv,: Model is nearly unidentifiable: large eigenvalue ratio
## - Rescale variables?
~~~

~~~
summary_h1p_EP_single <- summary(model_h1p_EP_single)
mySummary(summary_h1p_EP_single)
~~~

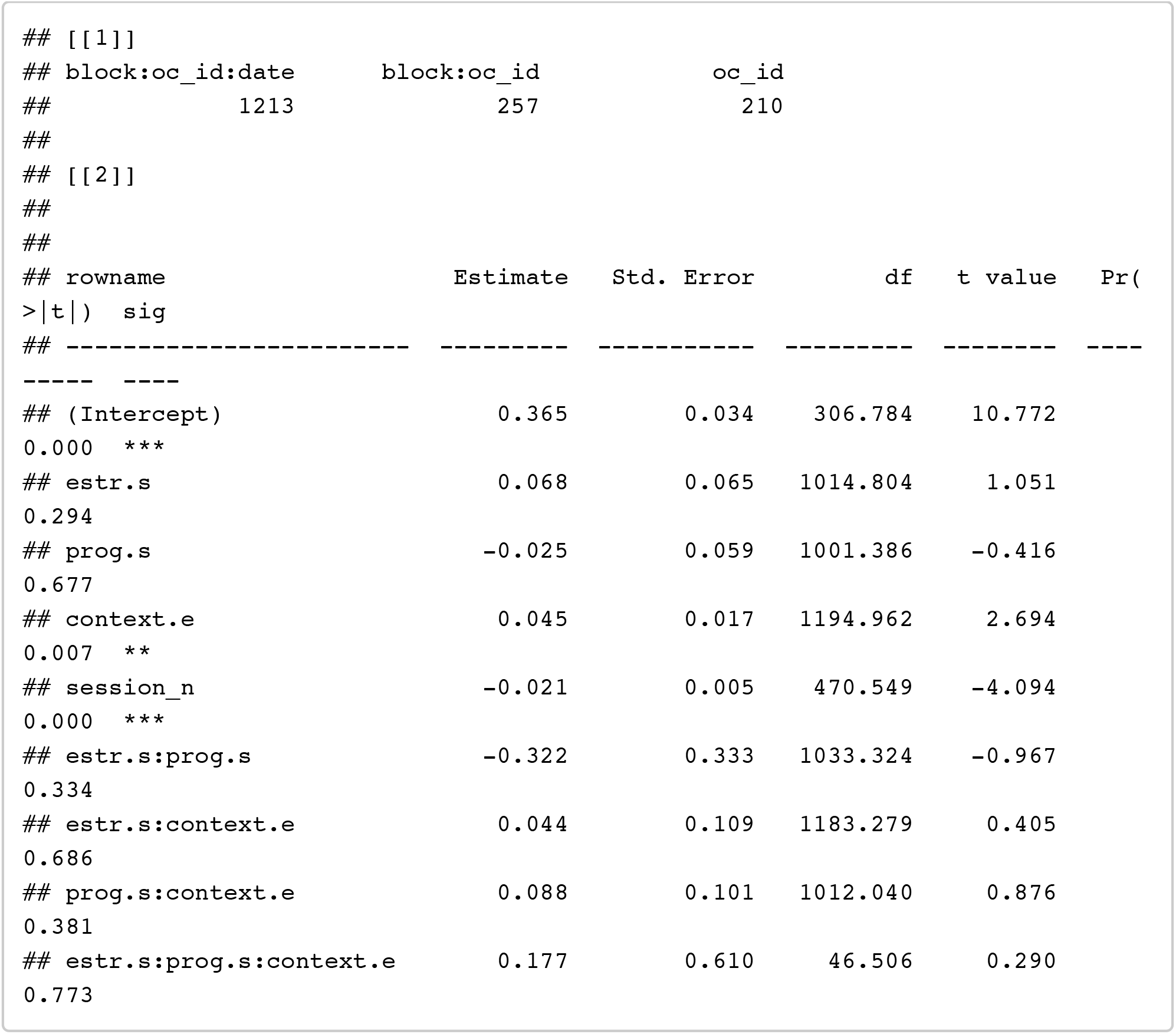

~~~
confint(model_h1p_EP_single, method = confint_method) %>% as.data.frame() %>% rownames_to_column() %>% filter(!is.na(‘2.5 %’))
~~~

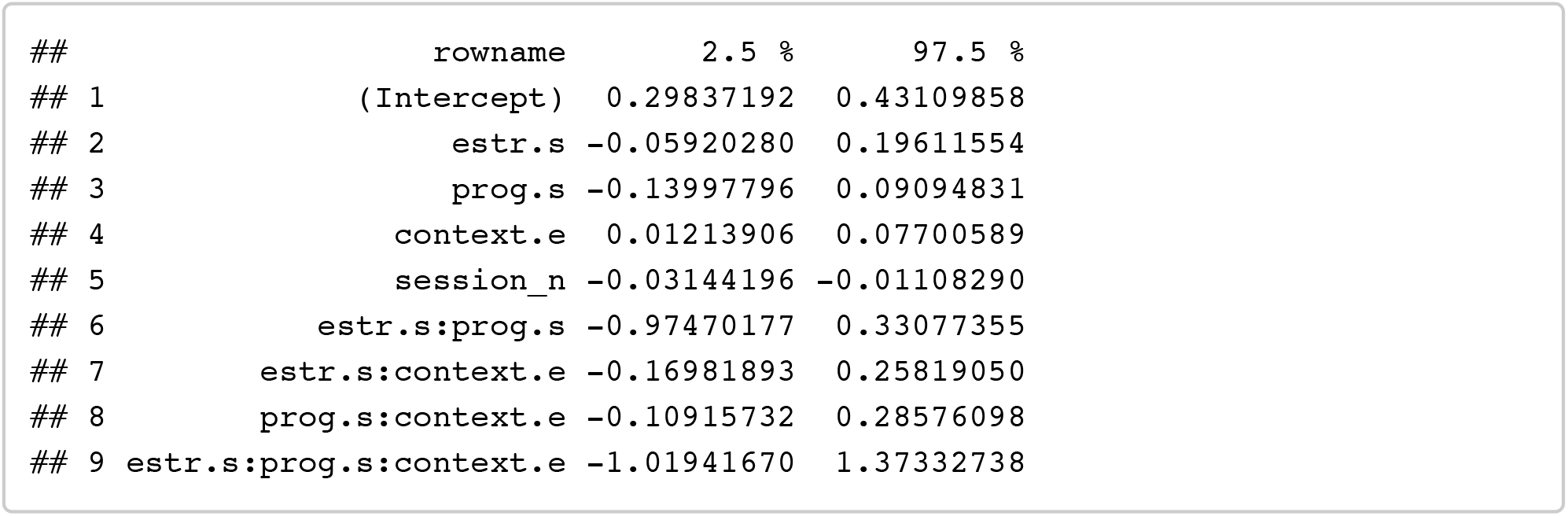

##### E + P + E*P: (partnered women only to interpret interaction)

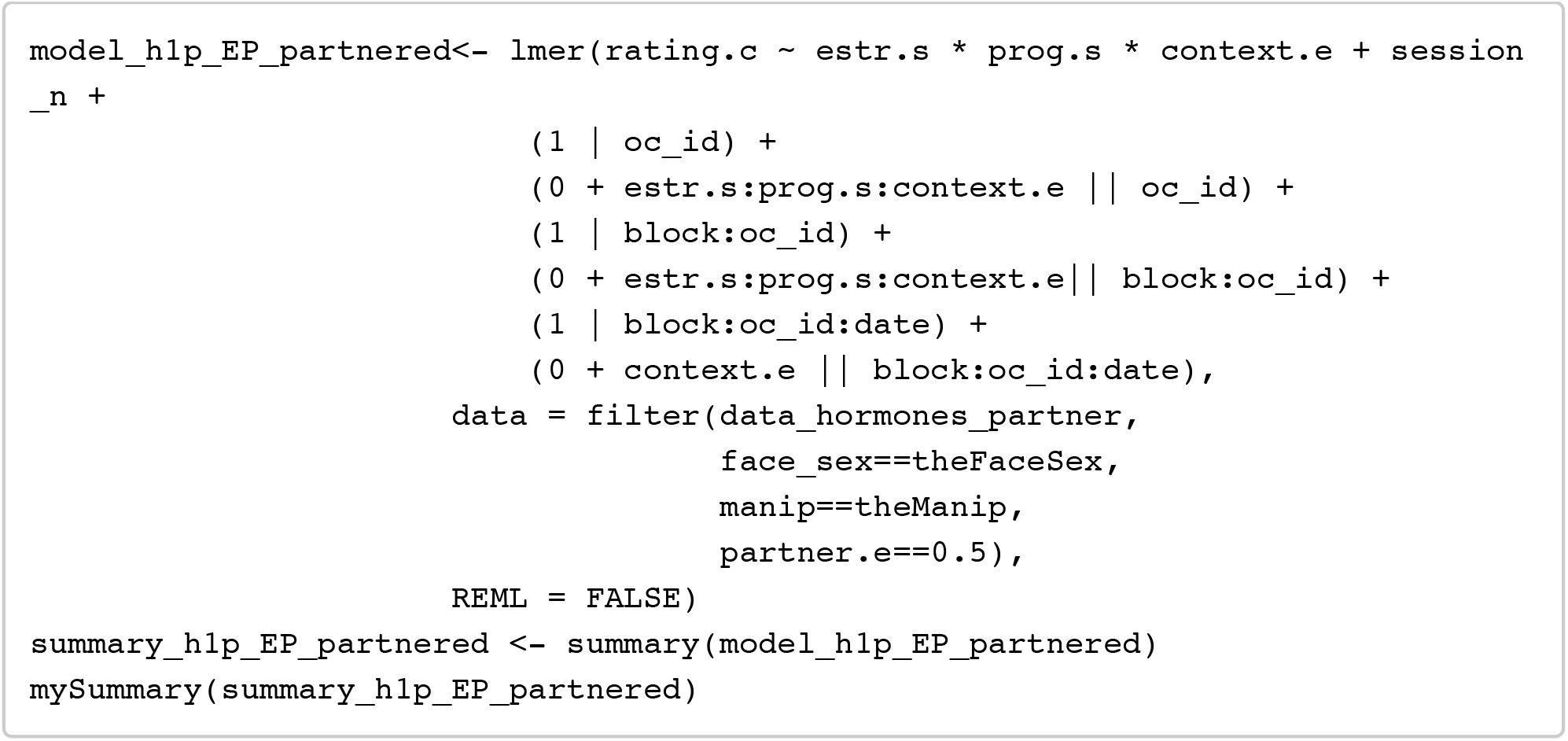

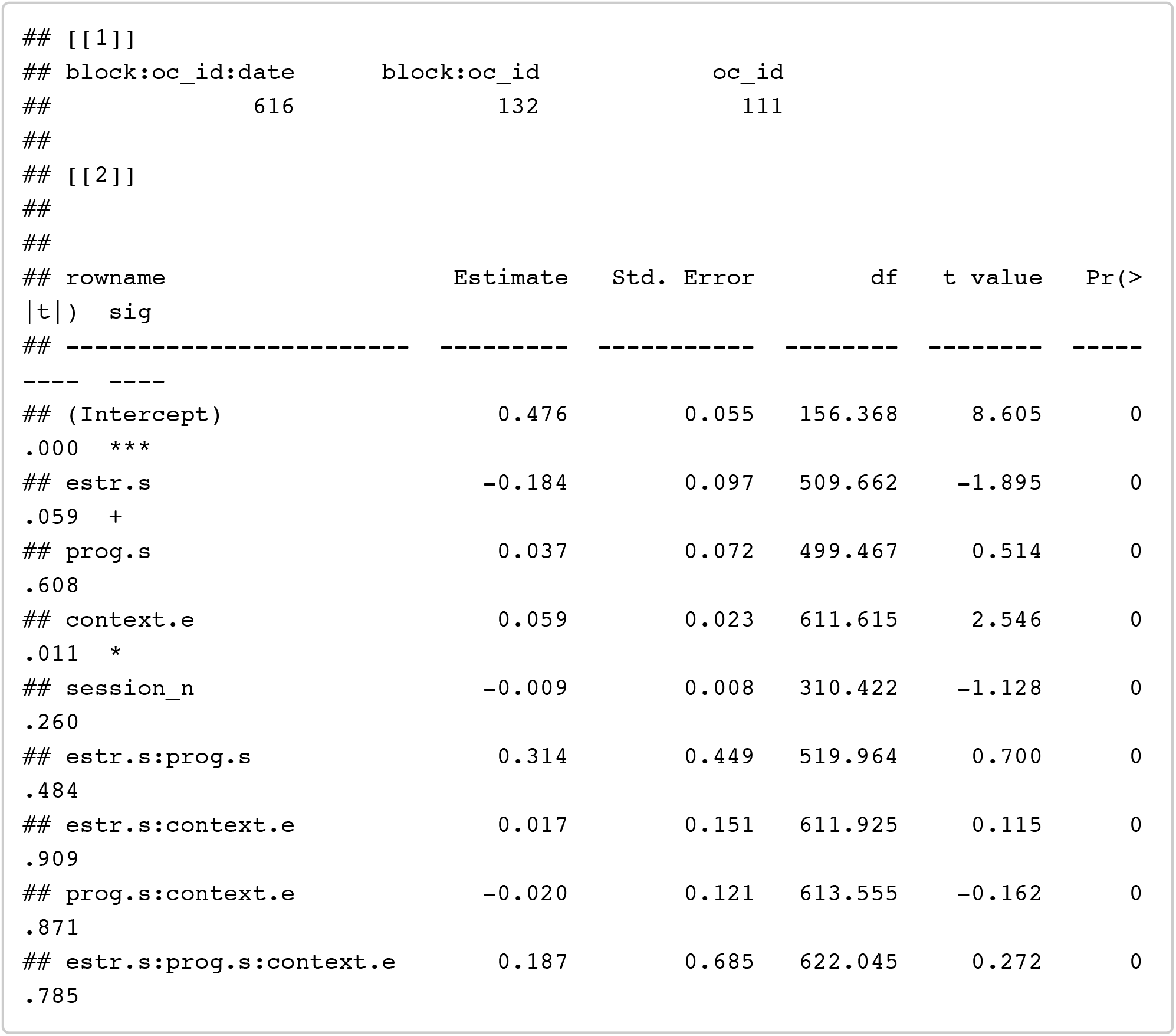

~~~
confint(model_h1p_EP_partnered, method = confint_method) %>% as.data.frame() % >% rownames_to_column() %>% filter(!is.na(‘2.5 %’))
~~~

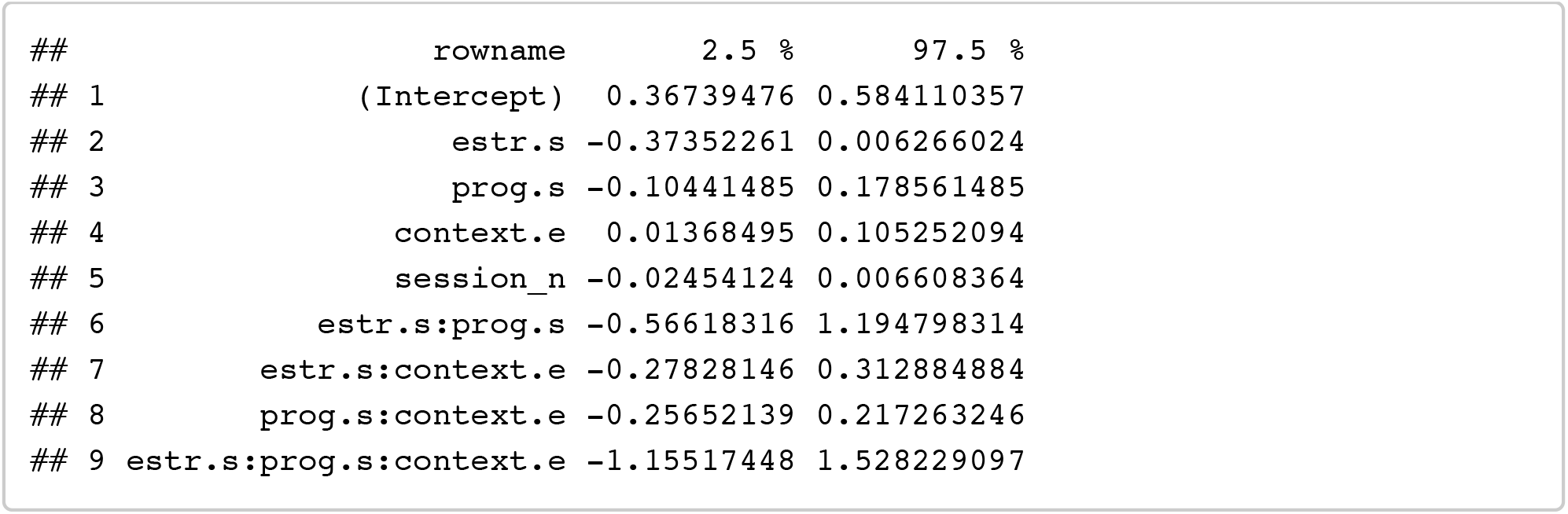

Note that the non-significant effect of estradiol (estimate = -0.18, p = .064) is in the opposite direction to what would be predicted from previous work reporting positive effects of estradiol on preferences.

##### E + P + EPratio: (+ partnership status)

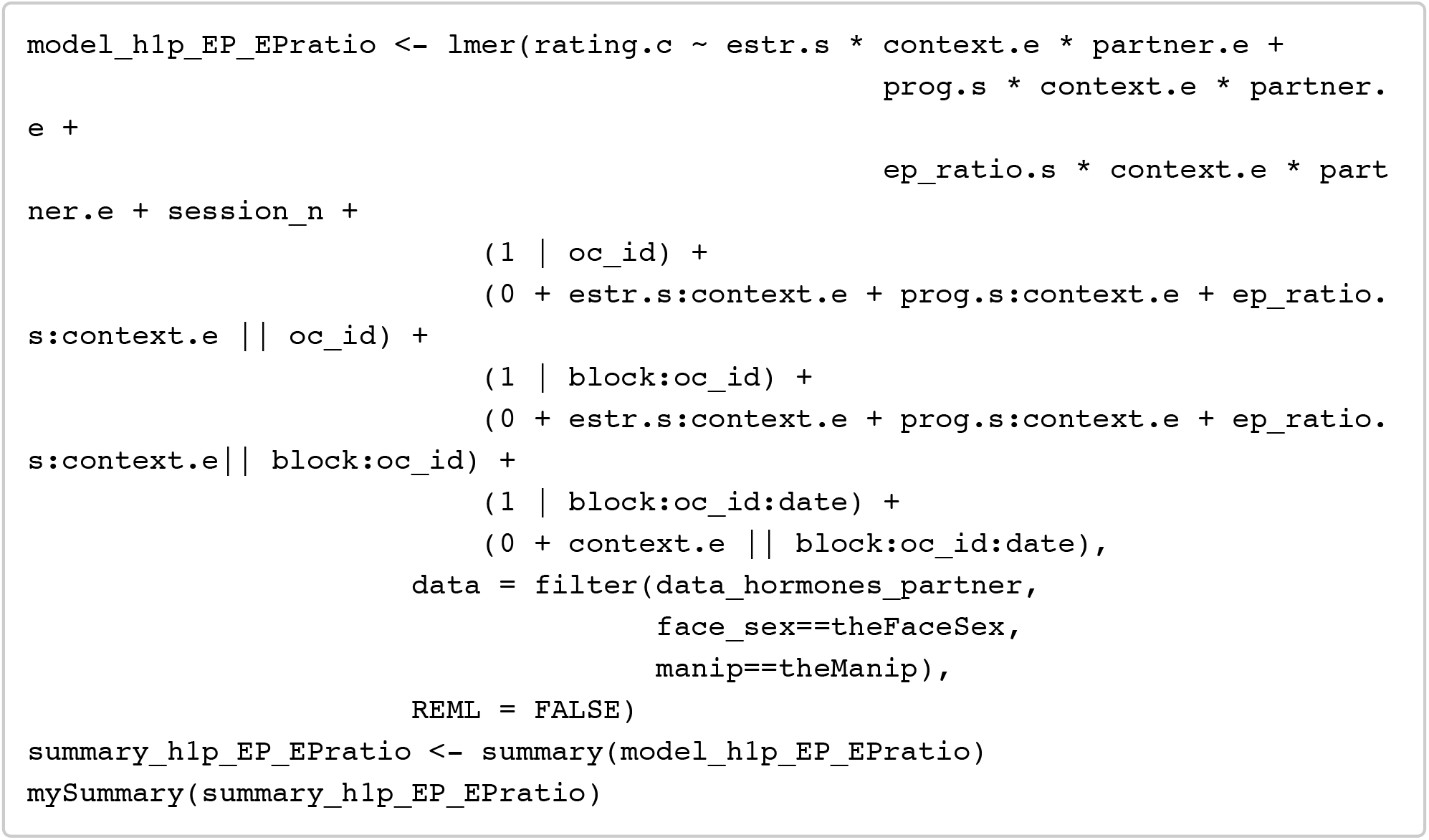

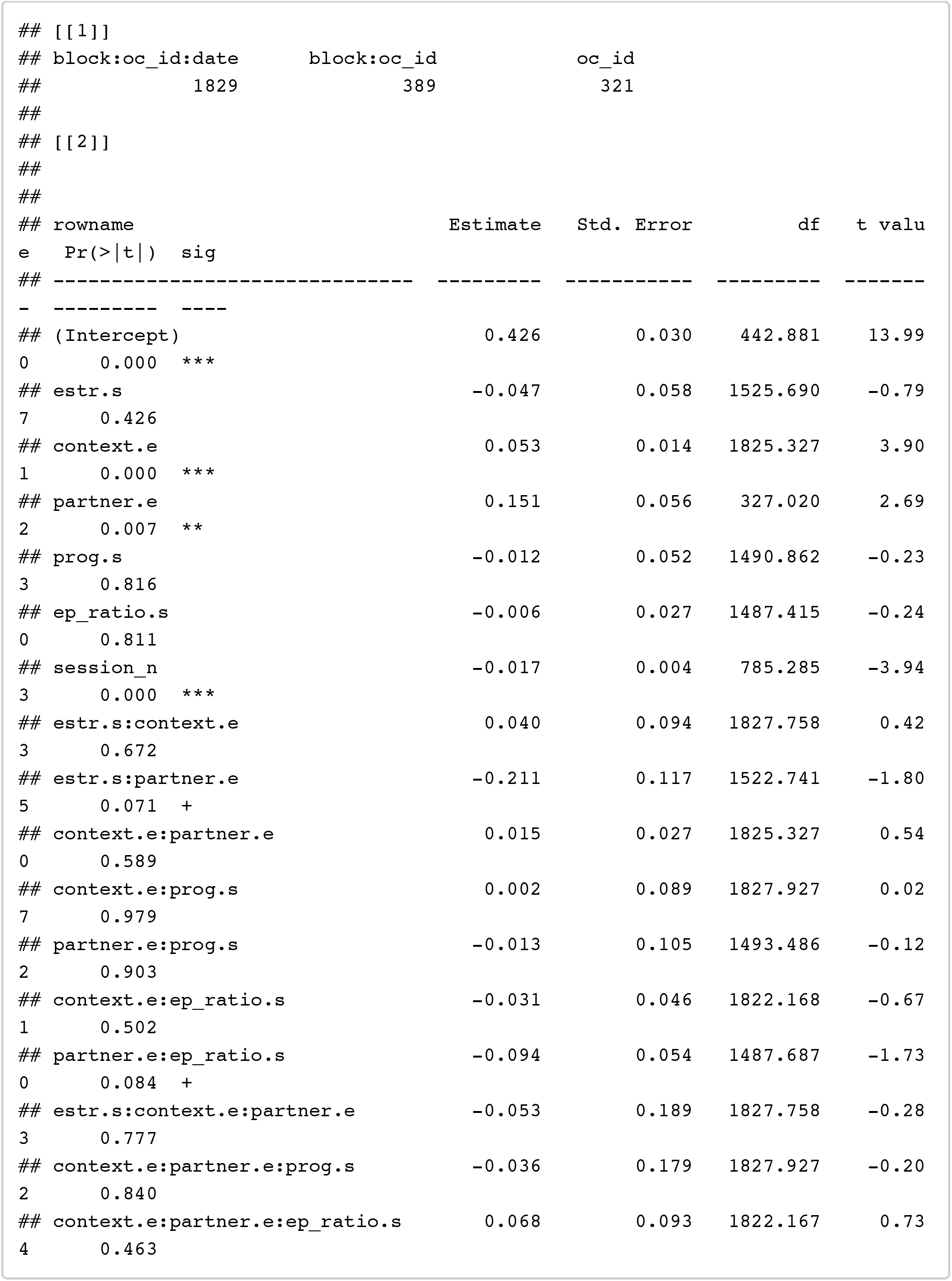

~~~
confint(model_h1p_EP_EPratio, method = confint_method) %>% as.data.frame() %>% rownames_to_column() %>% filter(!is.na(‘2.5 %’))
~~~

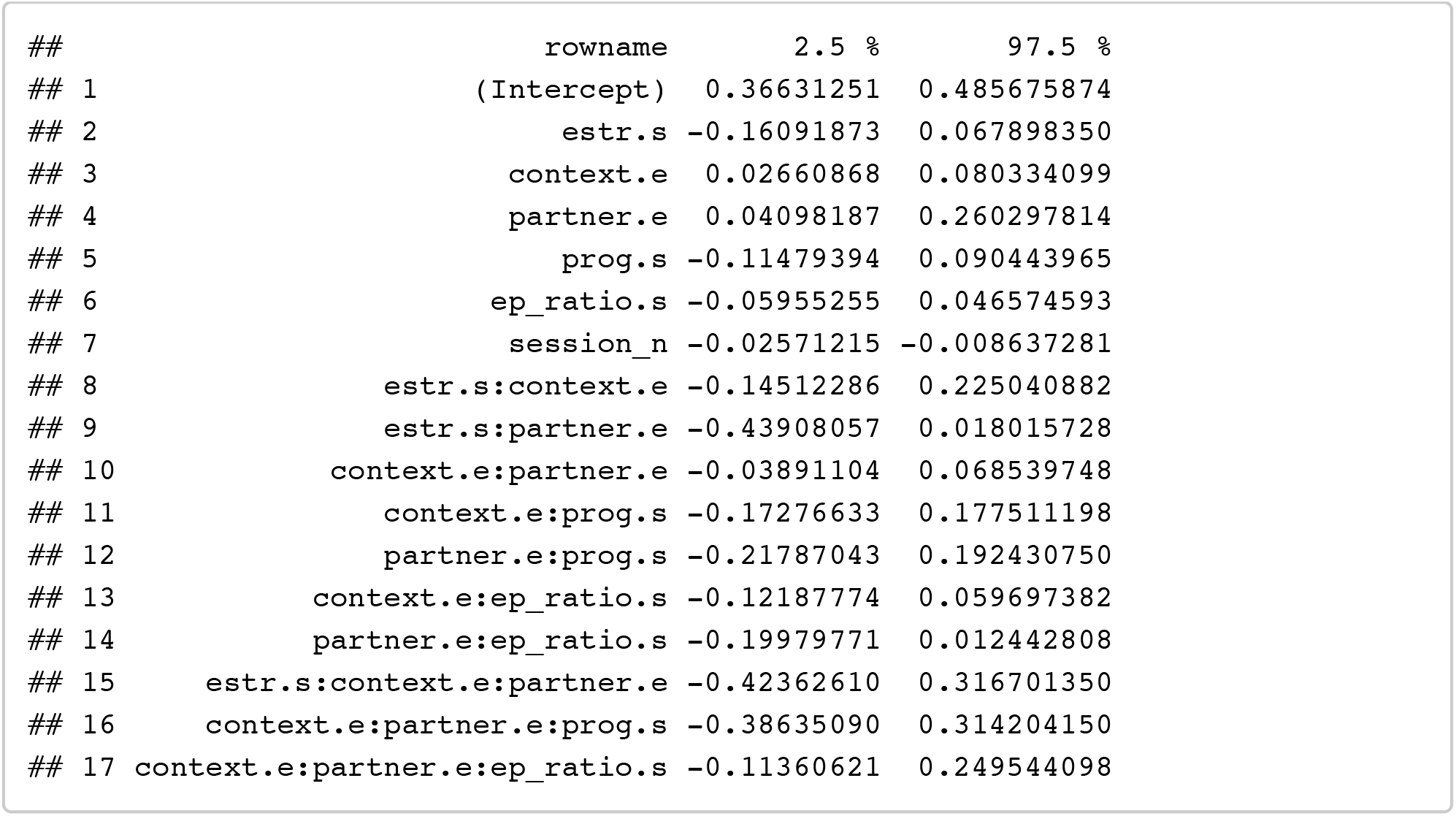

##### T + C: (+ partnership status)

Testing for effects of testosterone and coritsol on preferences

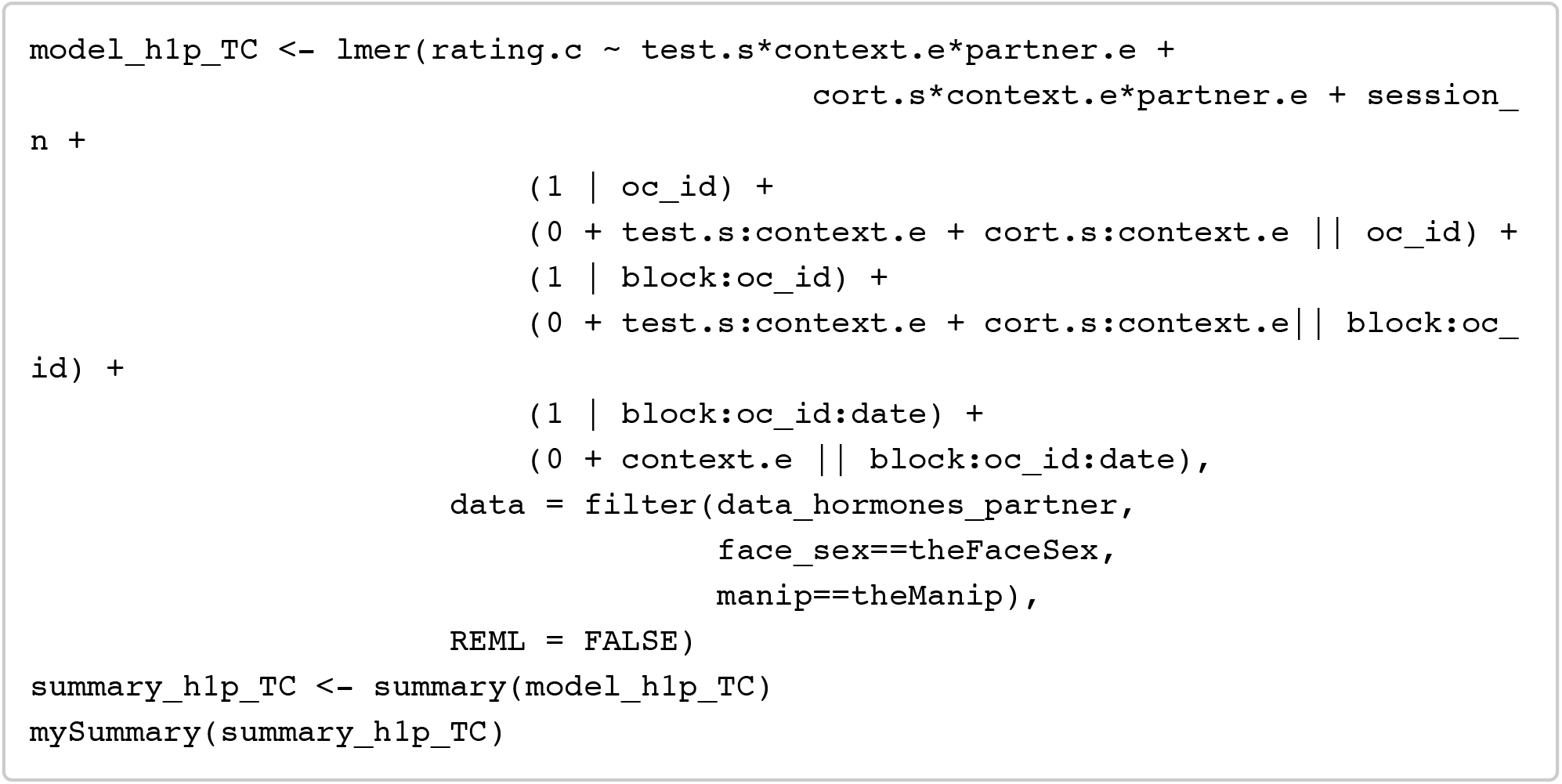

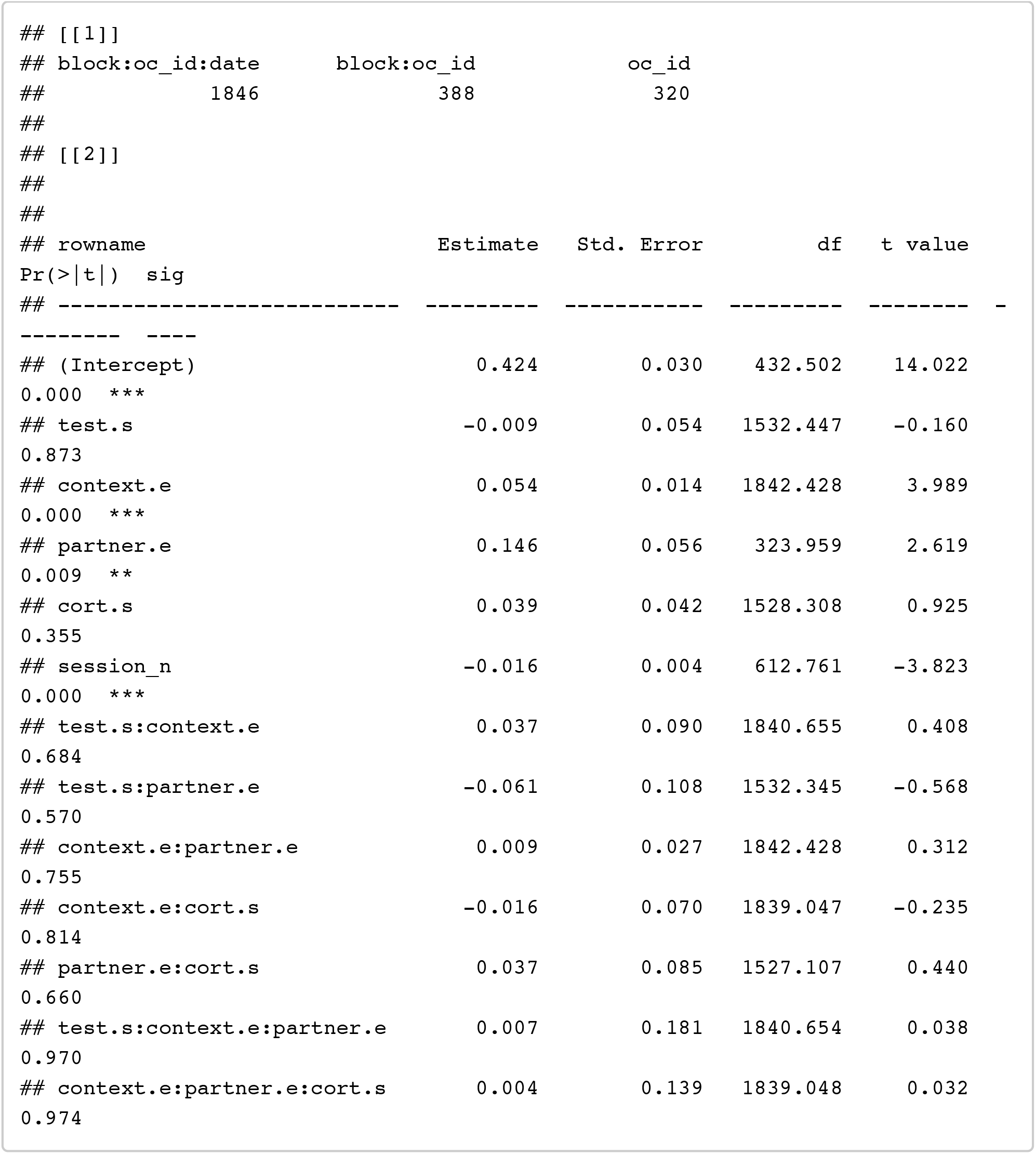

~~~
confint(model_h1p_TC, method = confint_method) %>% as.data.frame() %>% rowname s_to_column() %>% filter(!is.na(‘2.5 %’))
~~~

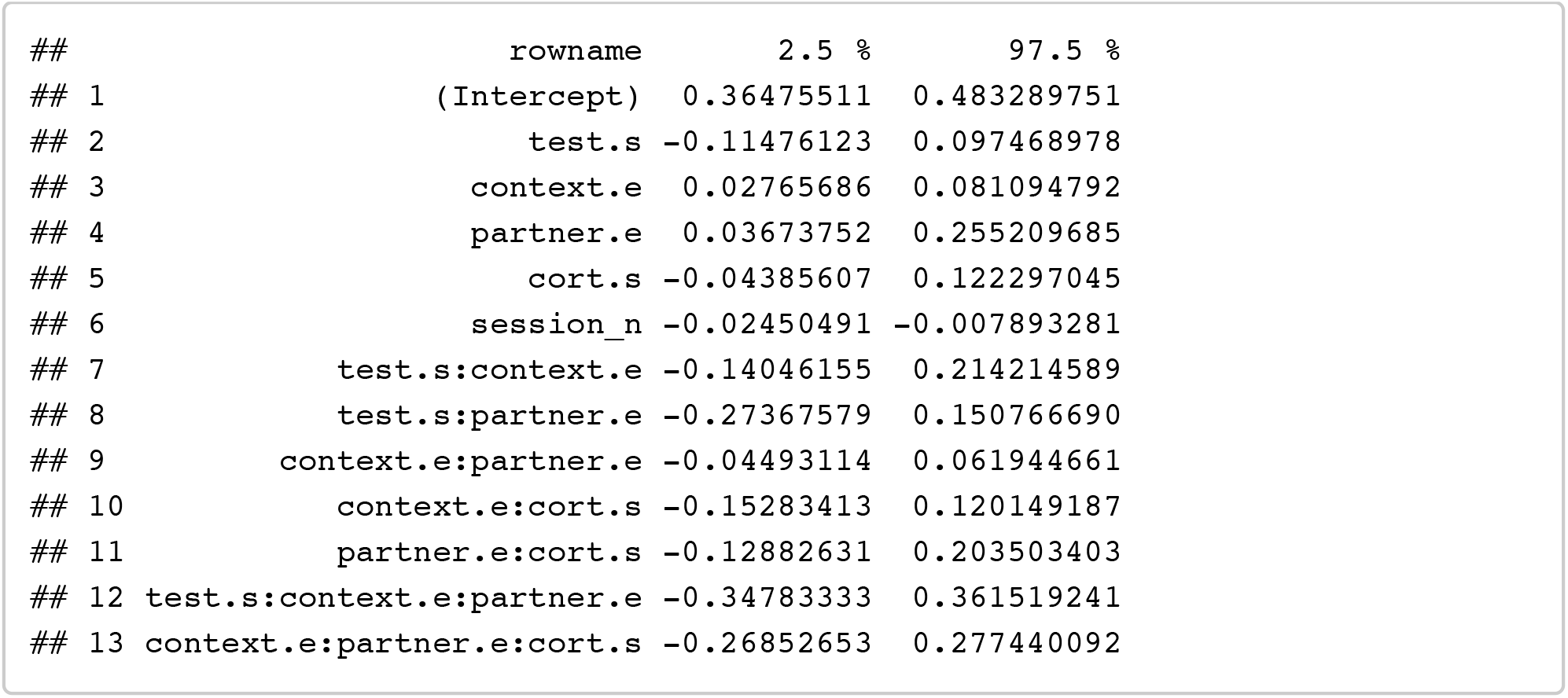

##### T + C + T*C: (+ partnership status)

Testing for effects of testosterone and coritsol plus their interaction on preferences

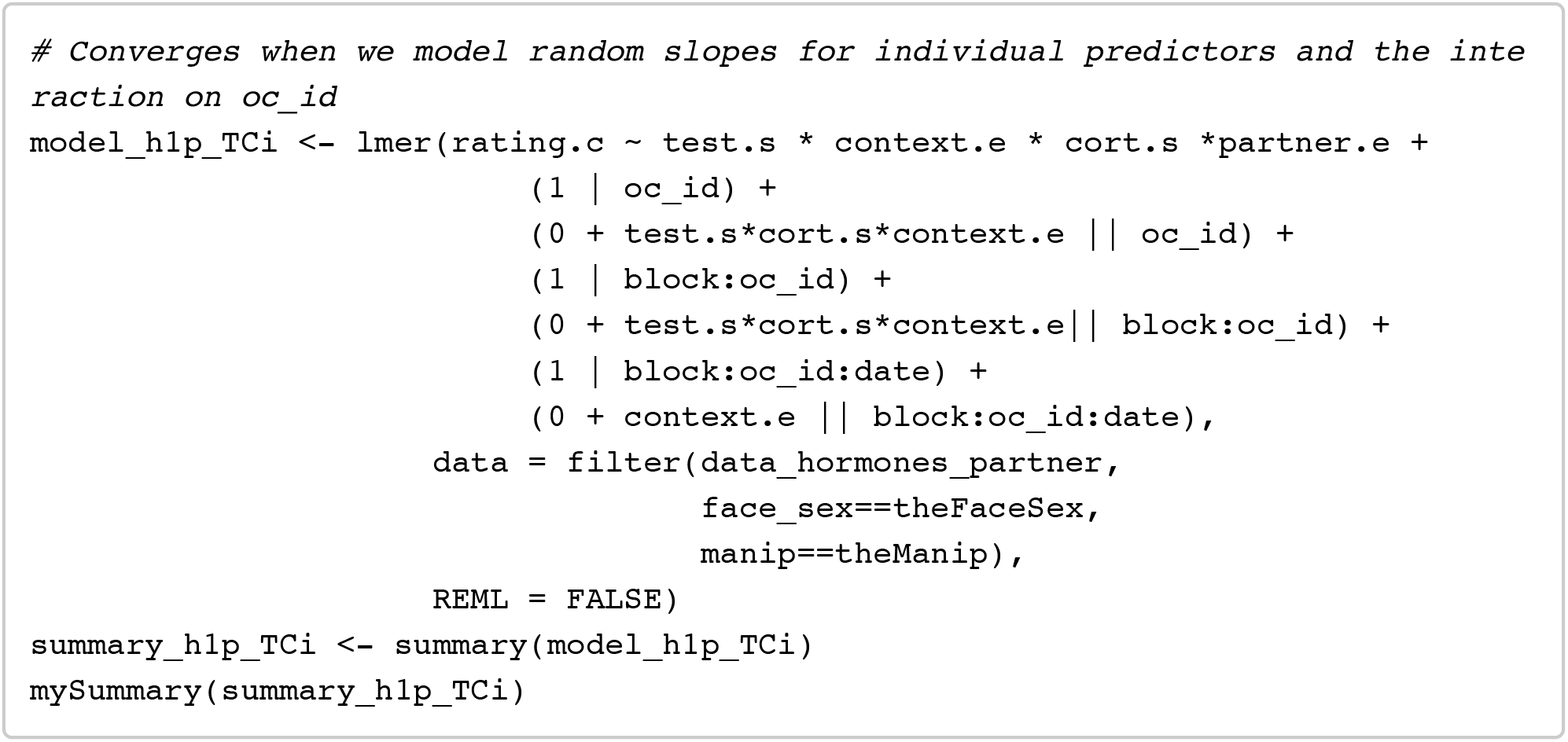

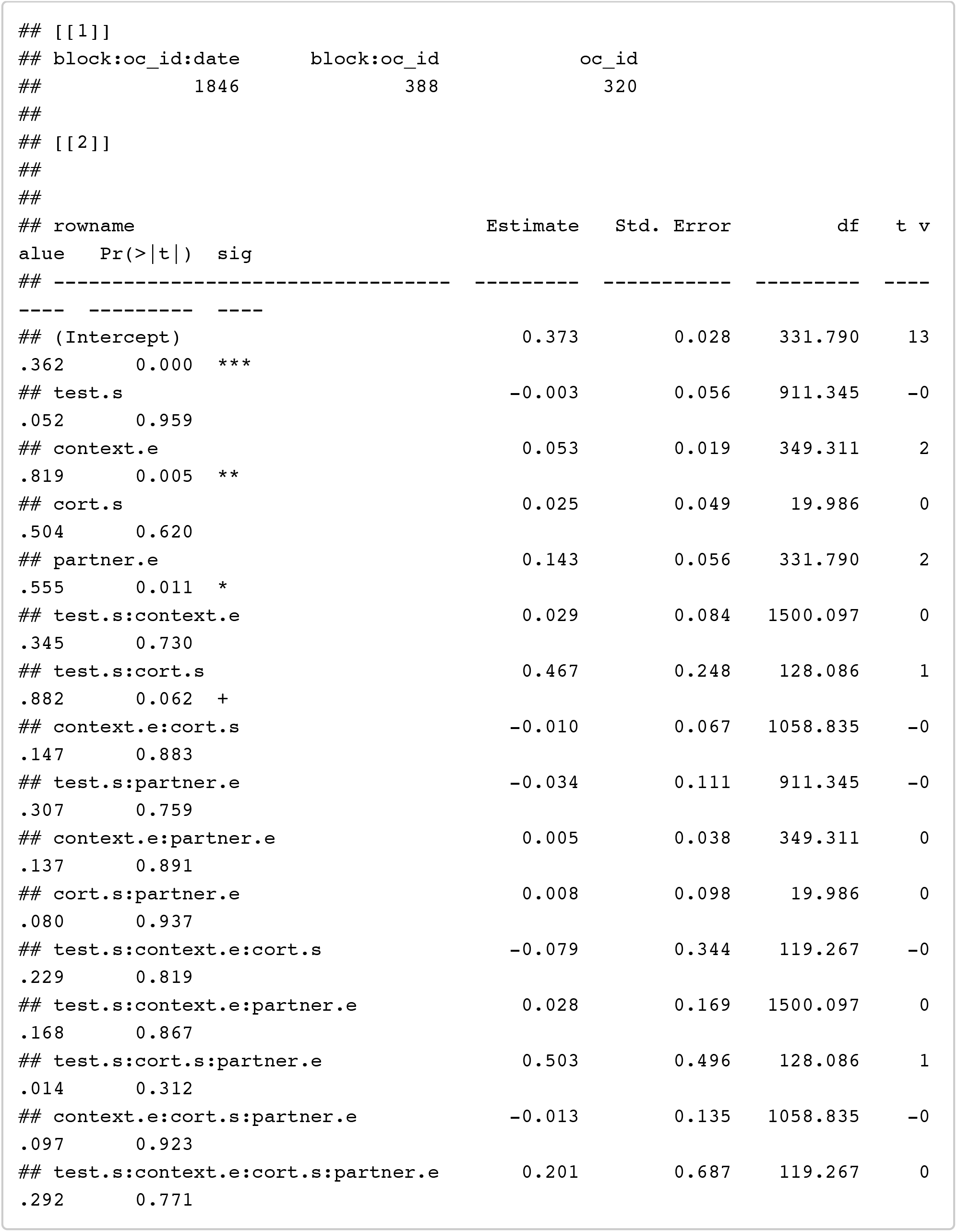

~~~
confint(model_h1p_TCi, method = confint_method) %>% as.data.frame() %>% rownam es_to_column() %>% filter(!is.na(‘2.5 %’))
~~~

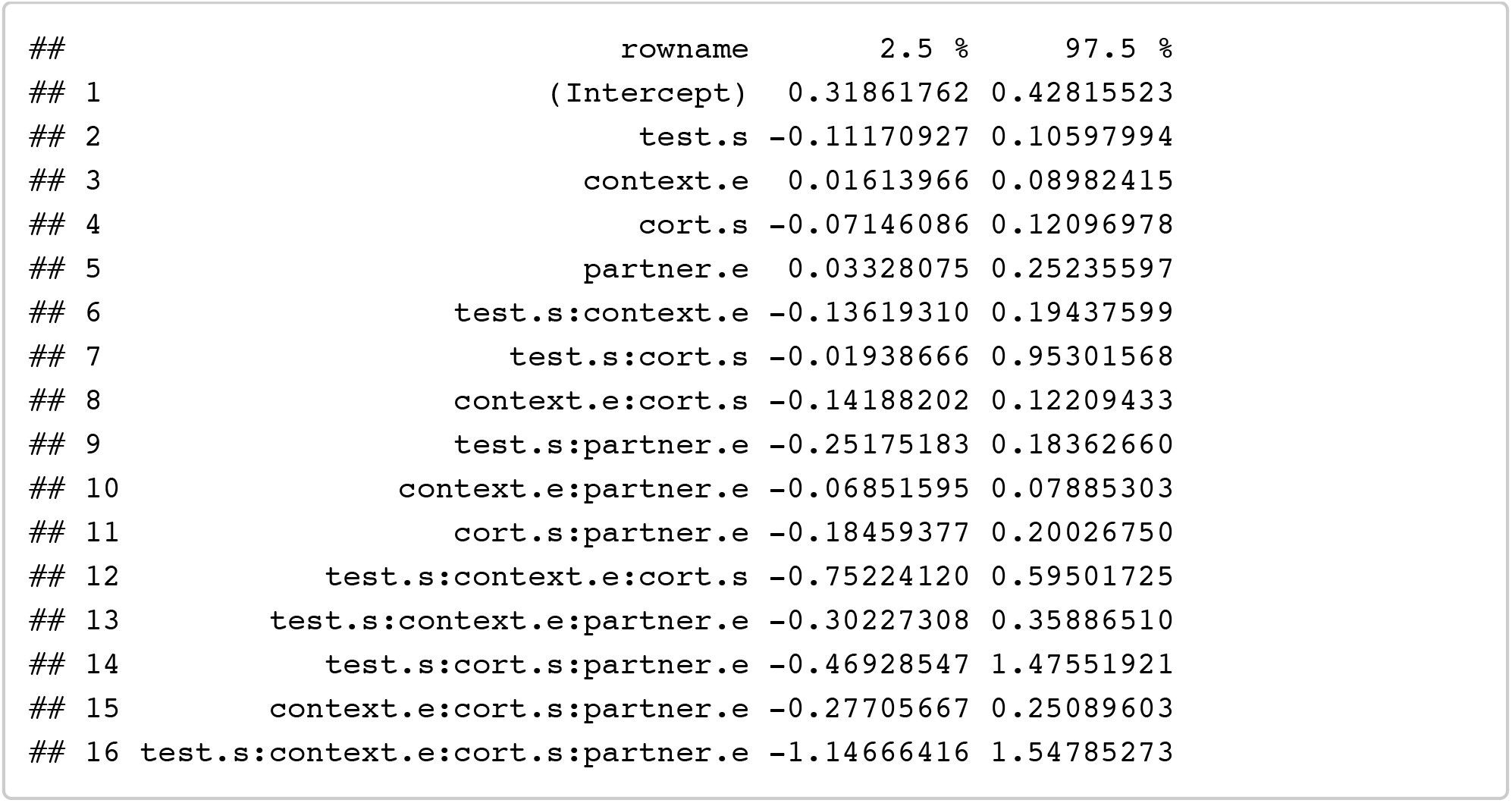

#### Analyses H1 ps: Hormones (+ session order, + partnership status)

##### E + P + E*P: (+ session order, + partnership status)

Testing for effects of estradiol, progesterone, and their interaction on preferences

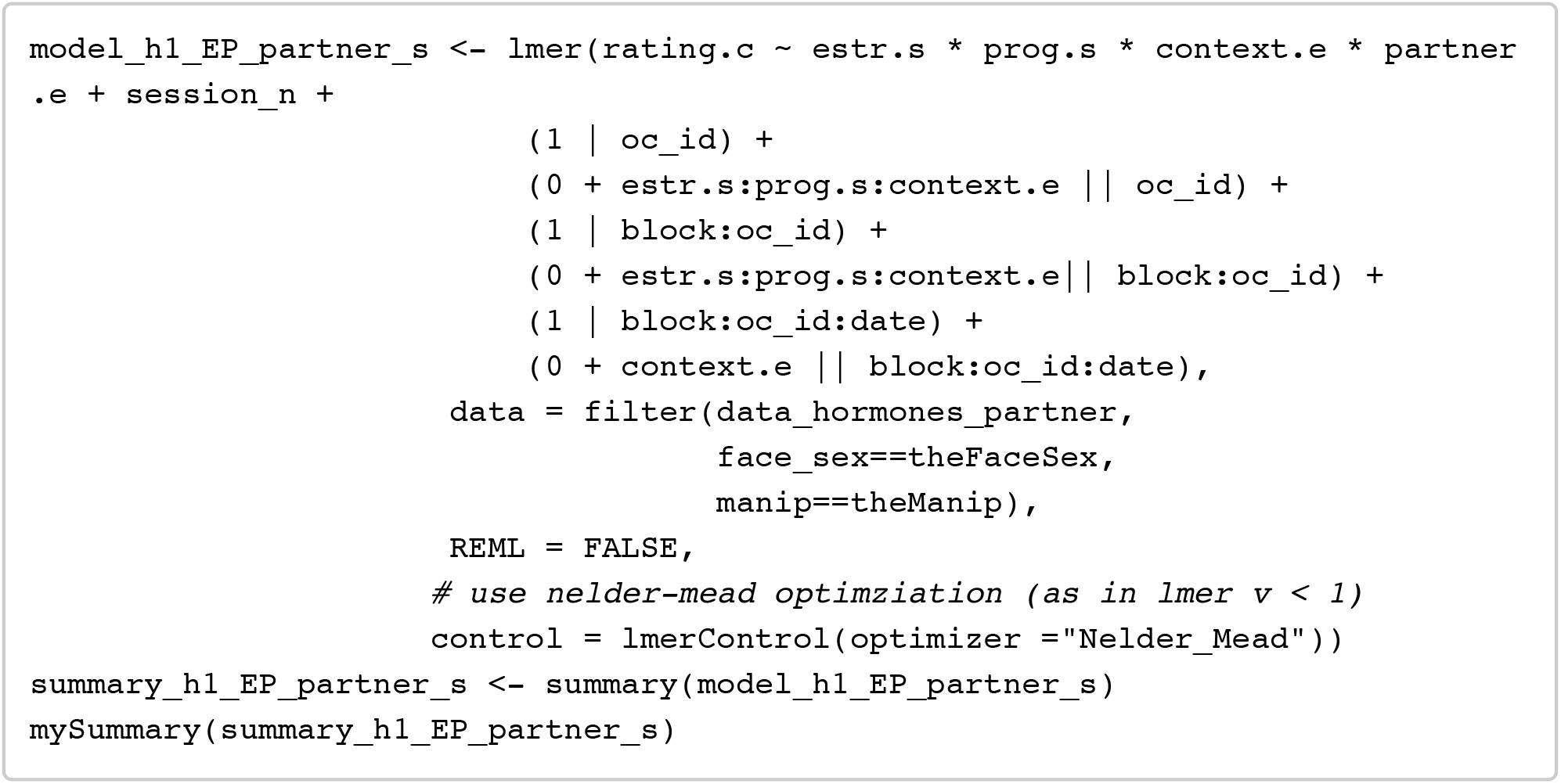

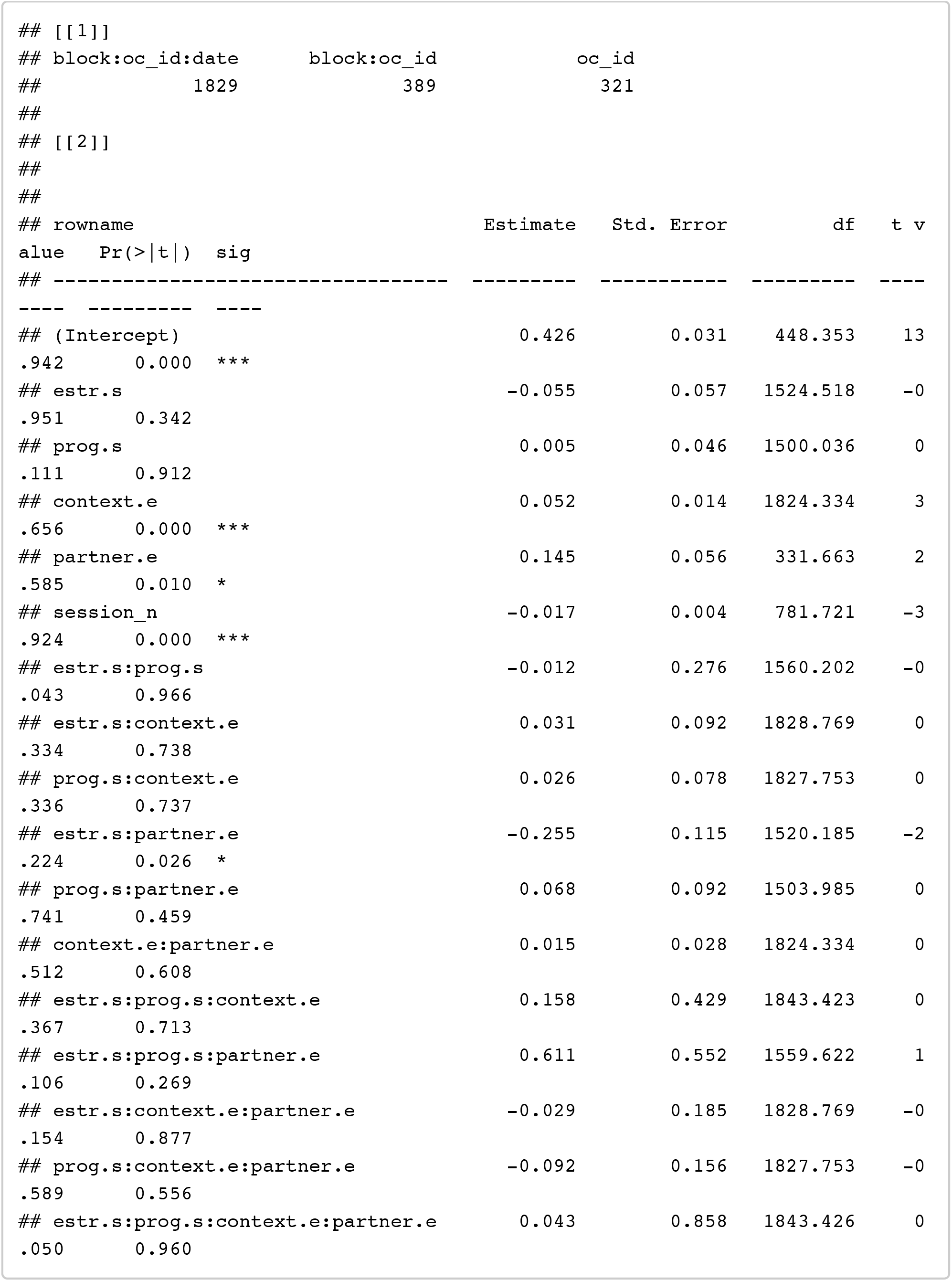

~~~
confint(model_h1_EP_partner_s, method = confint_method) %>% as.data.frame() %> % rownames_to_column() %>% filter(!is.na(‘2.5 %’))
~~~

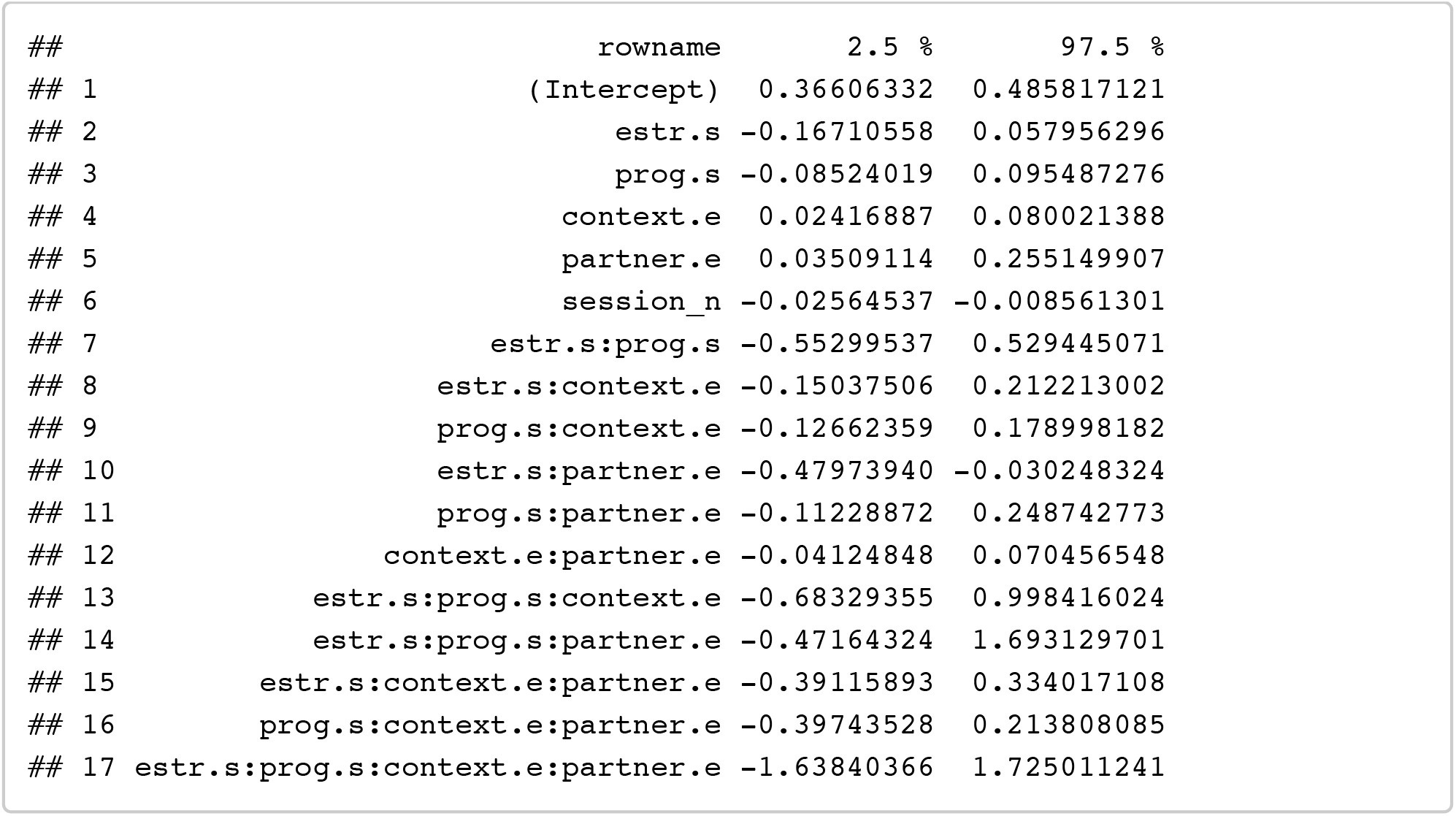

##### E + P + E*P: (single women only to interpret interaction)

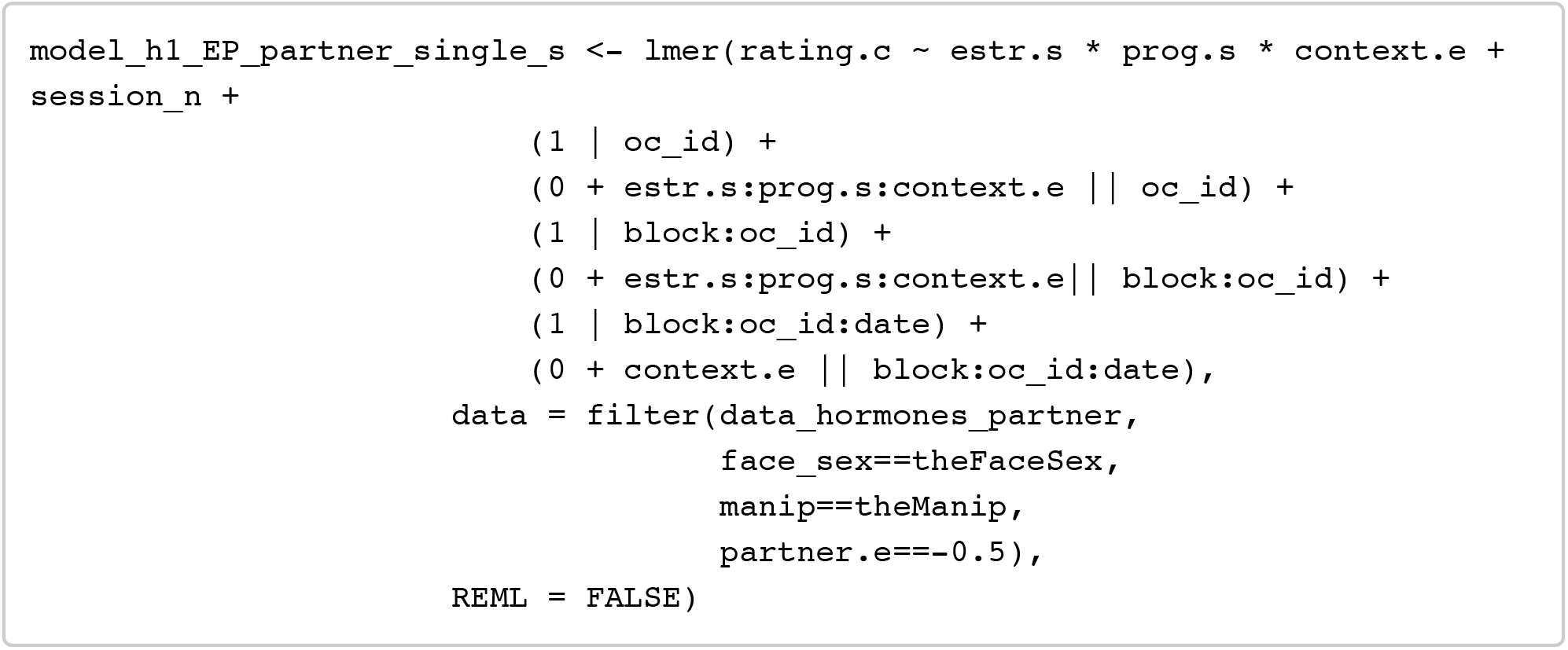

~~~
## Warning in checkConv(attr(opt, “derivs”), opt$par, ctrl = control$checkConv ,: Model is nearly unidentifiable: large eigenvalue ratio
## - Rescale variables?
~~~

~~~
summary_h1_EP_partner_single_s <- summary(model_h1_EP_partner_single_s) mySummary(summary_h1_EP_partner_single_s)
~~~

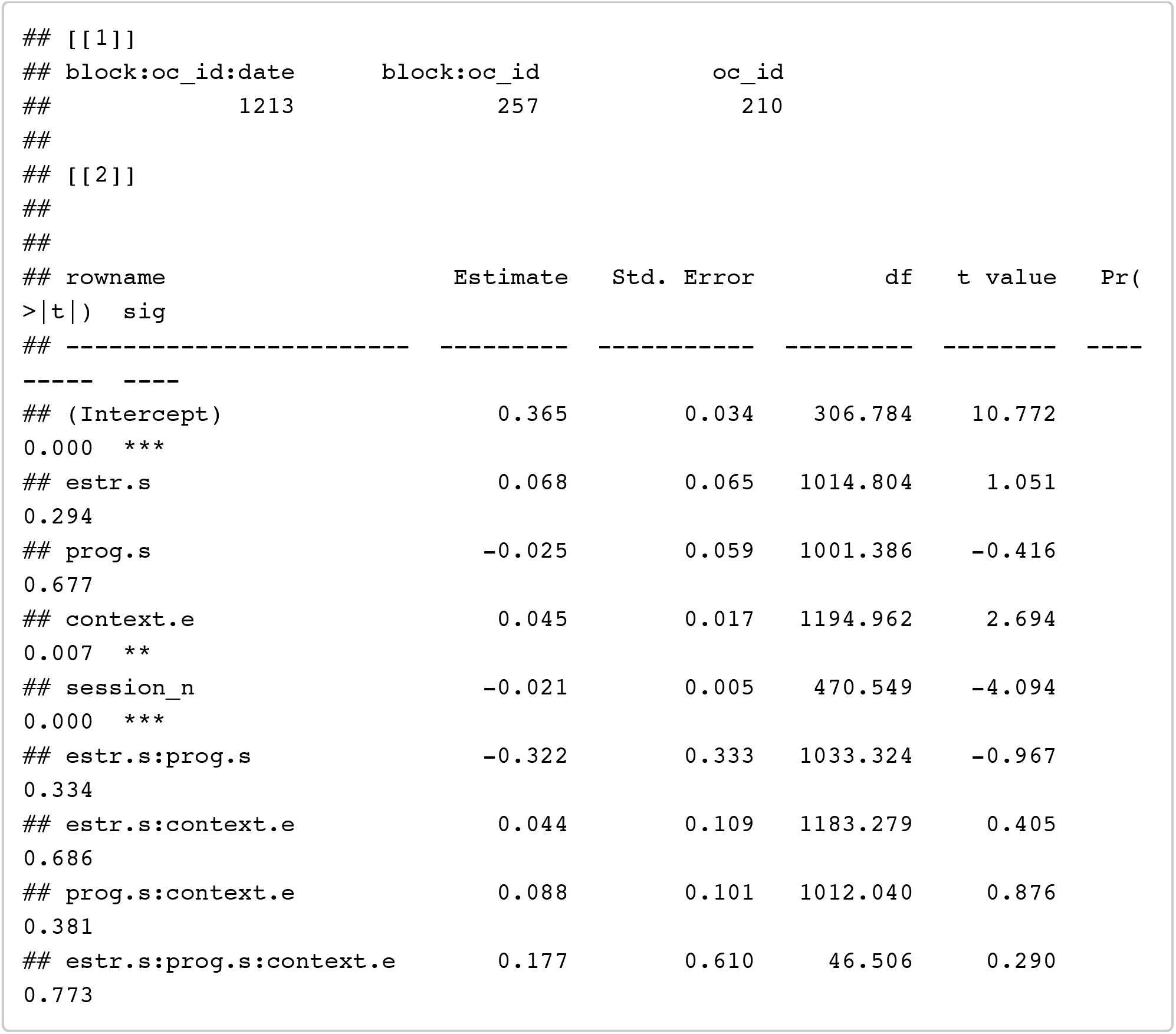

~~~
confint(model_h1_EP_partner_single_s, method = confint_method) %>% as.data.frame() %>% rownames_to_column() %>% filter(!is.na(‘2.5 %’))
~~~

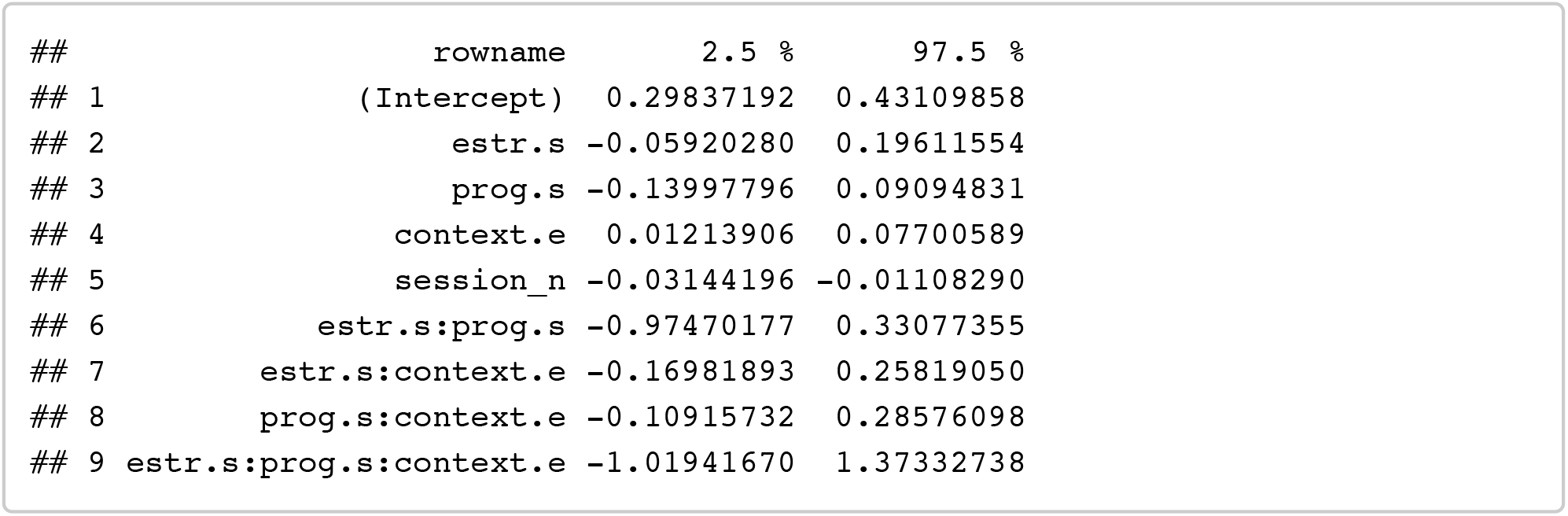

##### E + P + E*P: (partnered women only to interpret interaction)

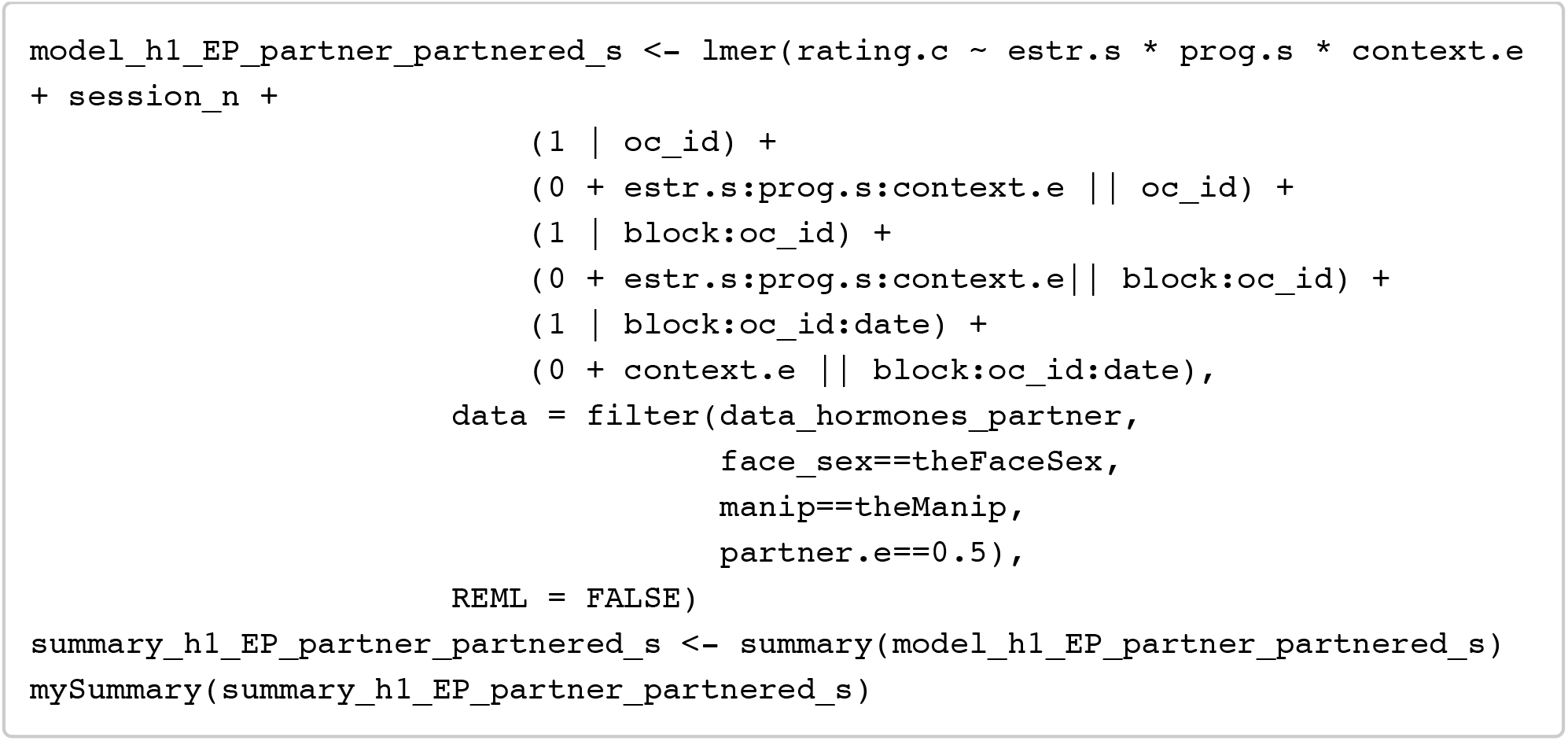

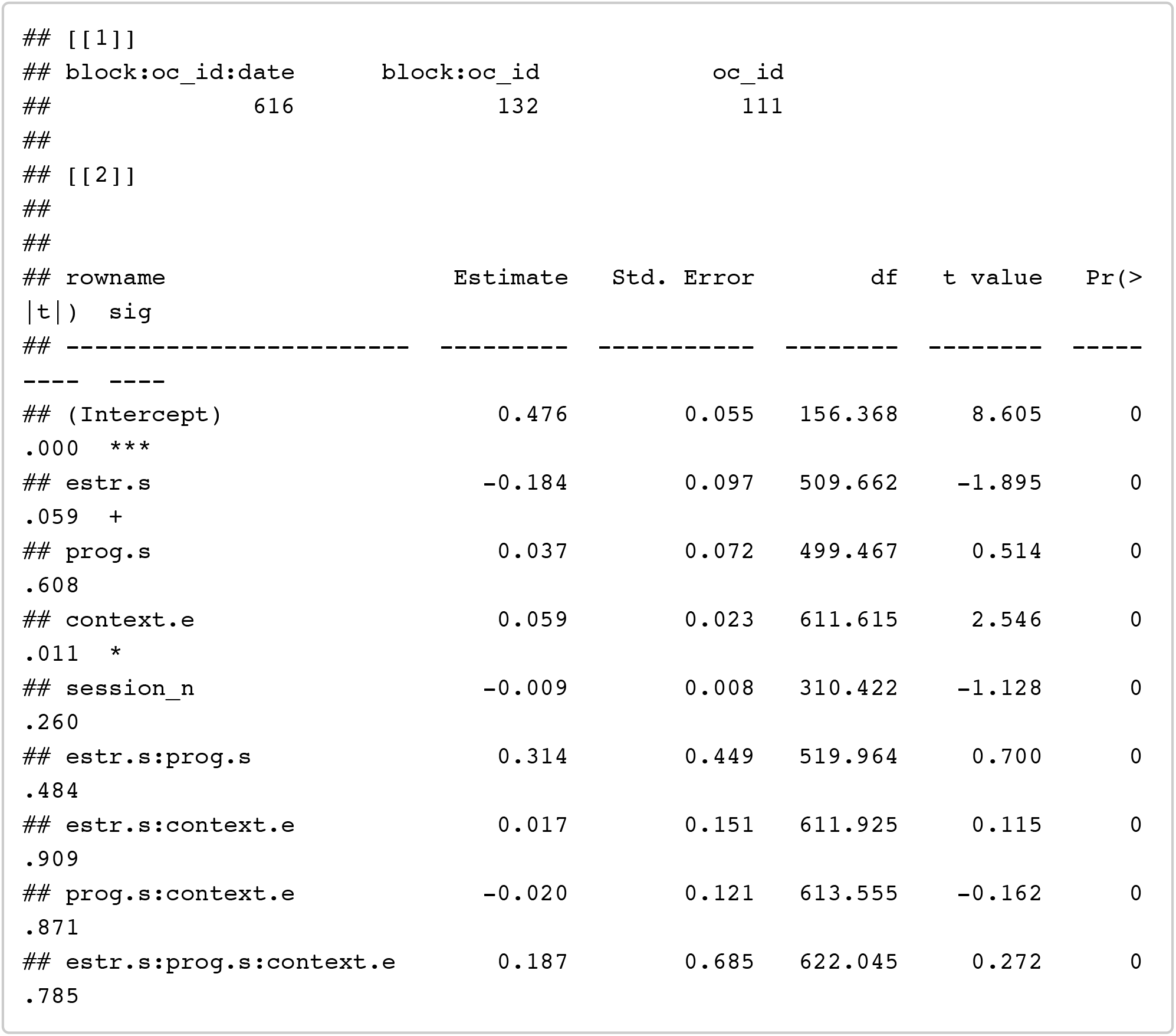

~~~
confint(model_h1_EP_partner_partnered_s, method = confint_method) %>% as.data. frame() %>% rownames_to_column() %>% filter(!is.na(‘2.5 %’))
~~~

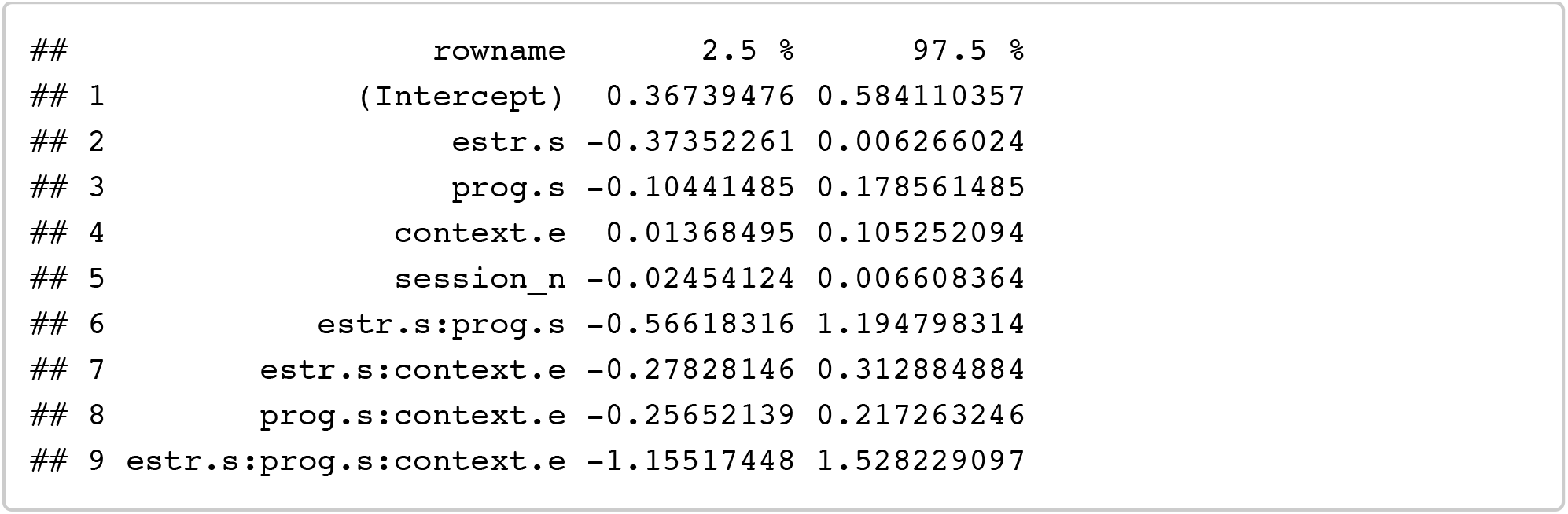

Note that the non-significant effect of estradiol (estimate = -0.18, p = .064) is in the opposite direction to what would be predicted from previous work reporting positive effects of estradiol.

##### E + P + EPratio: (+ session order, + partnership status)

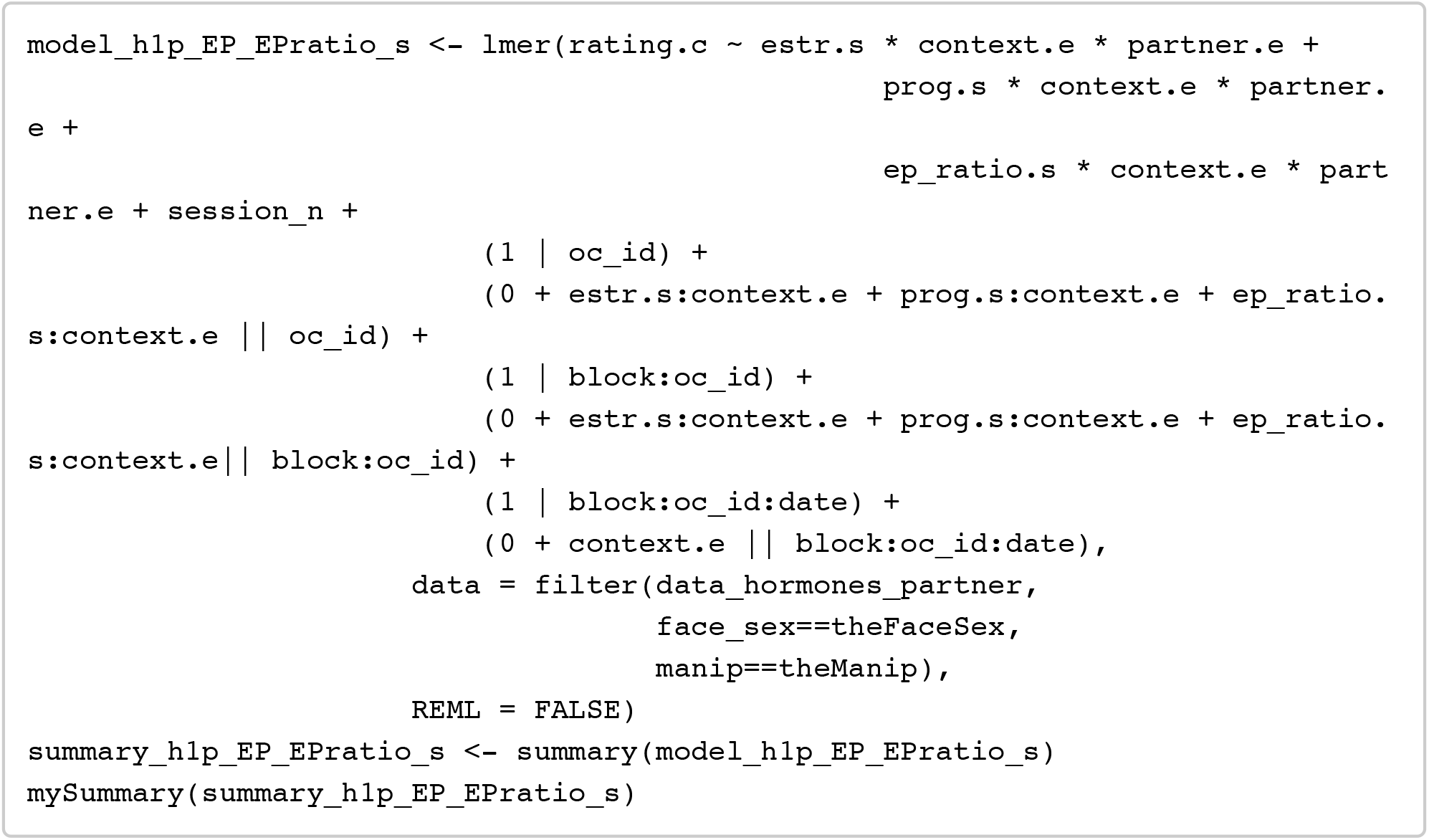

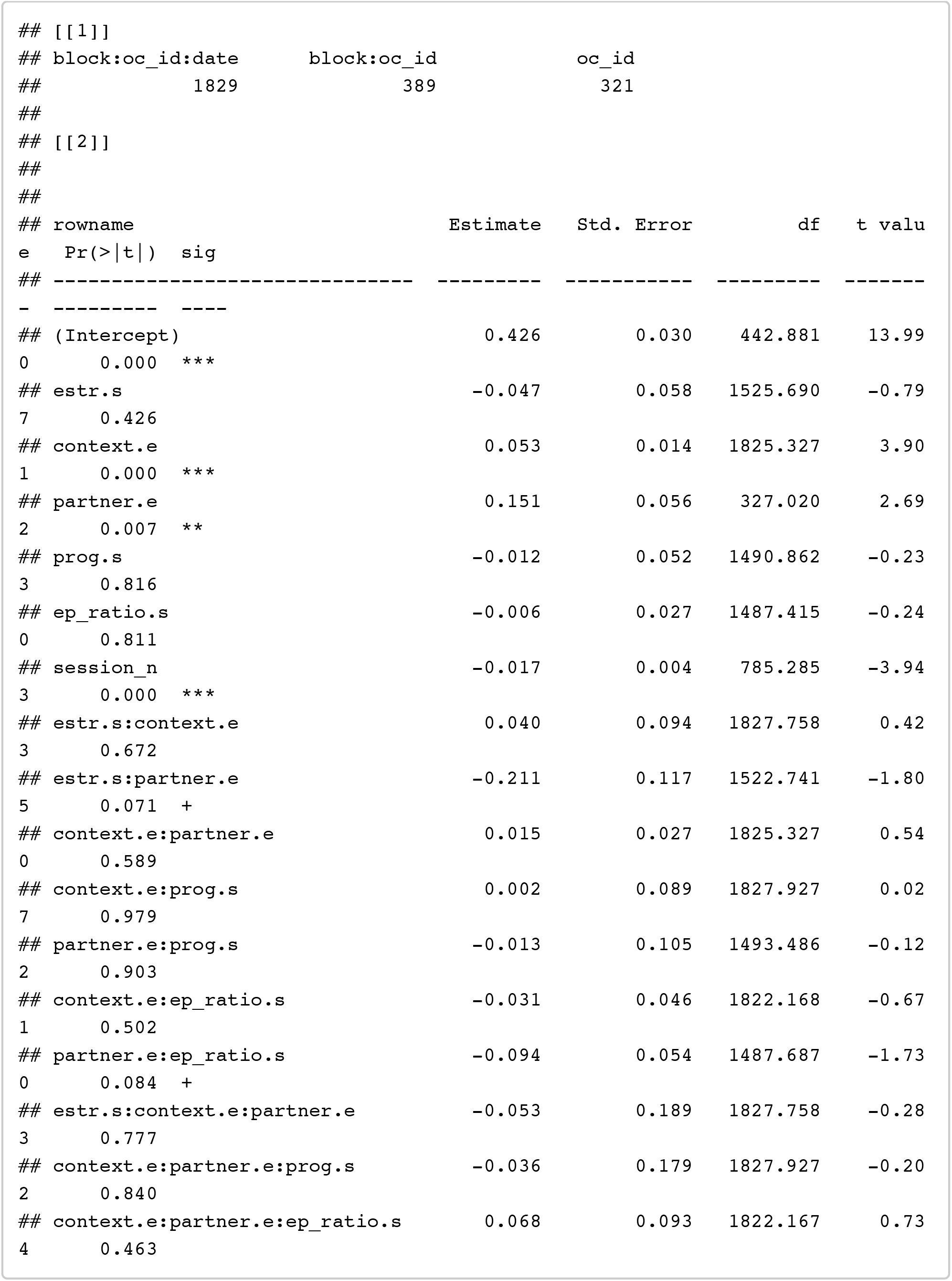

~~~
confint(model_h1p_EP_EPratio_s, method = confint_method) %>% as.data.frame() % >% rownames_to_column() %>% filter(!is.na(‘2.5 %’))
~~~

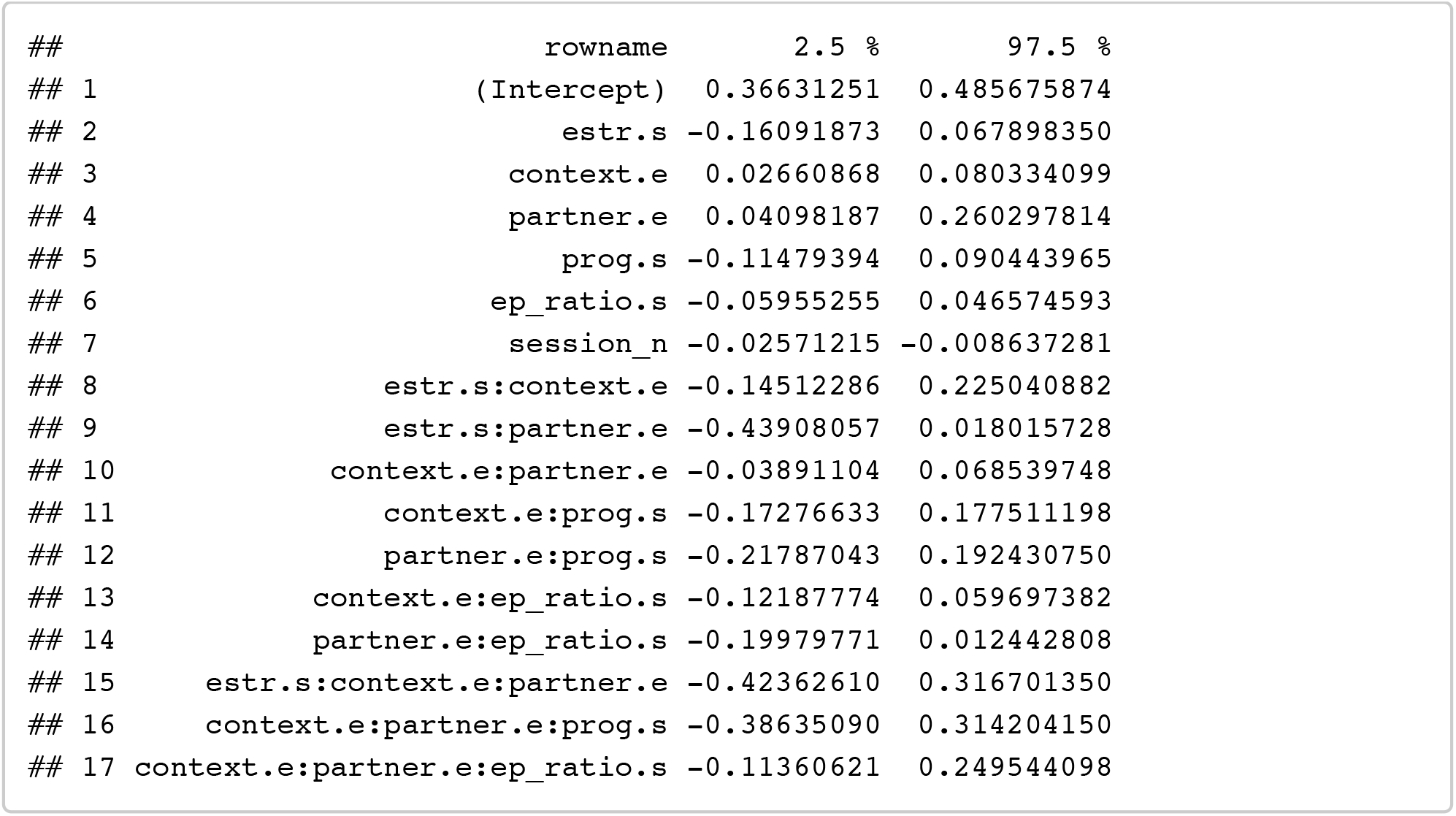

##### T + C: (+ session order, + partnership status)

Testing for effects of testosterone and coritsol on preferences

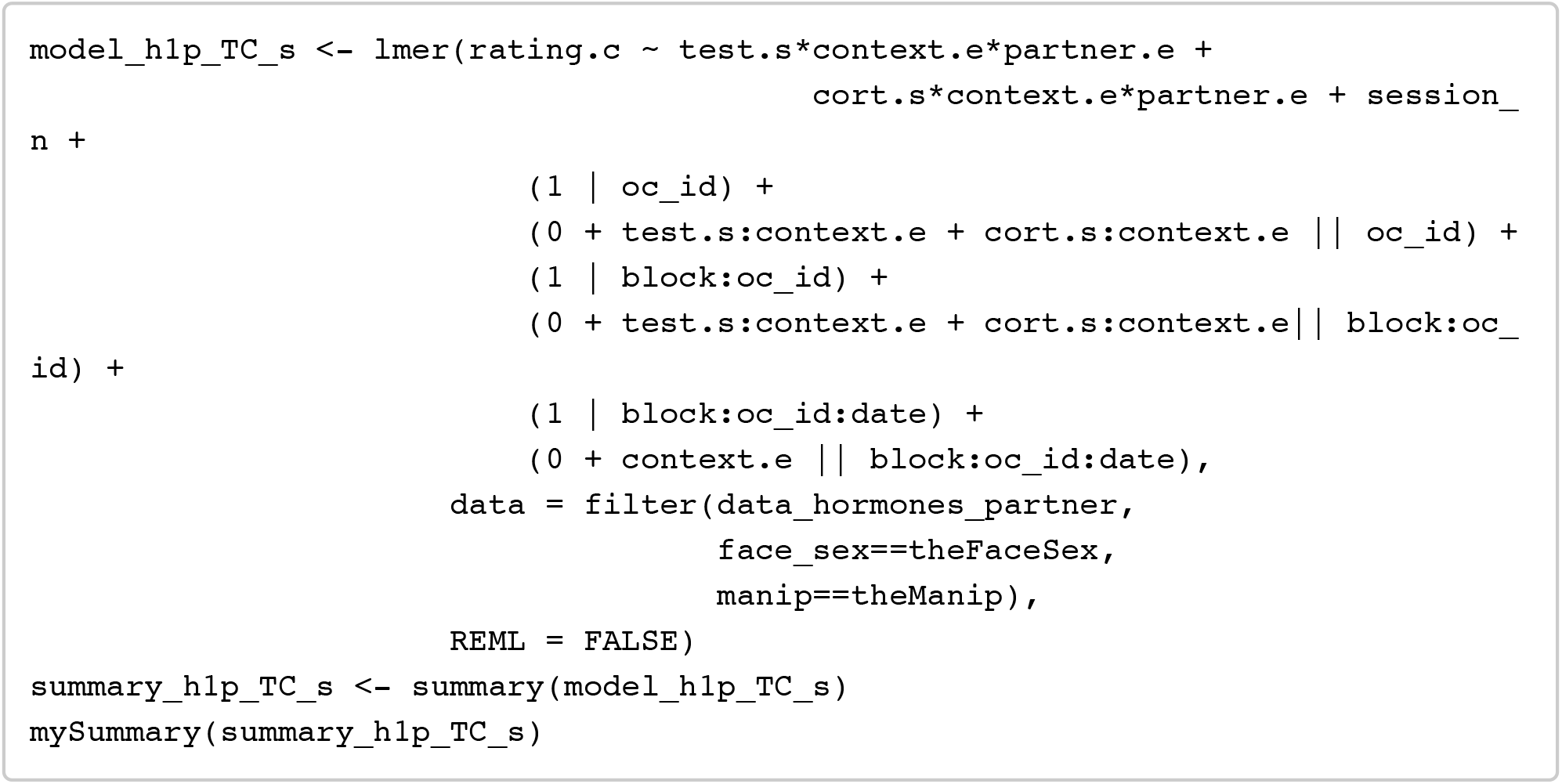

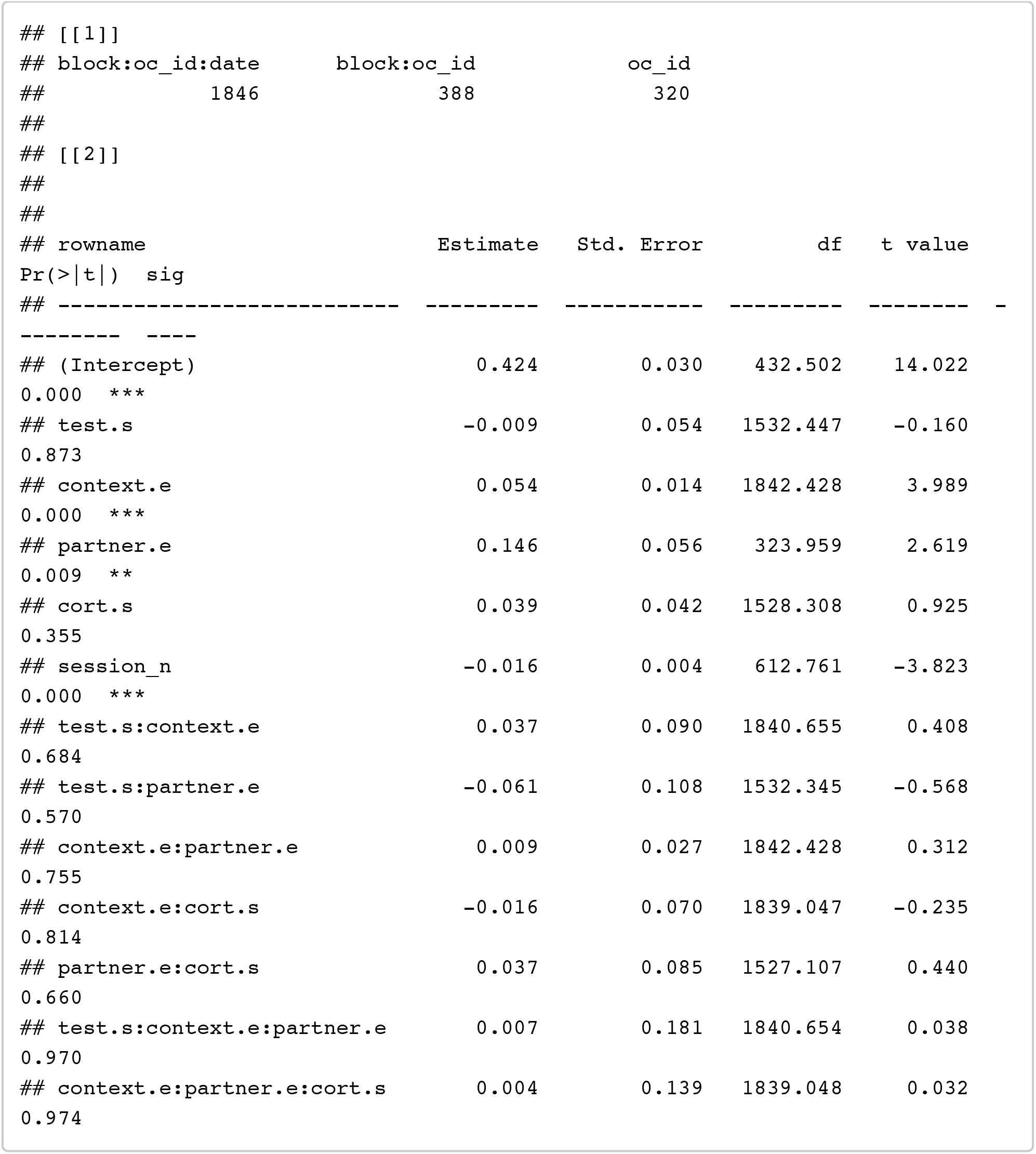

~~~
confint(model_h1p_TC_s, method = confint_method) %>% as.data.frame() %>% rowna mes_to_column() %>% filter(!is.na(‘2.5 %’))
~~~

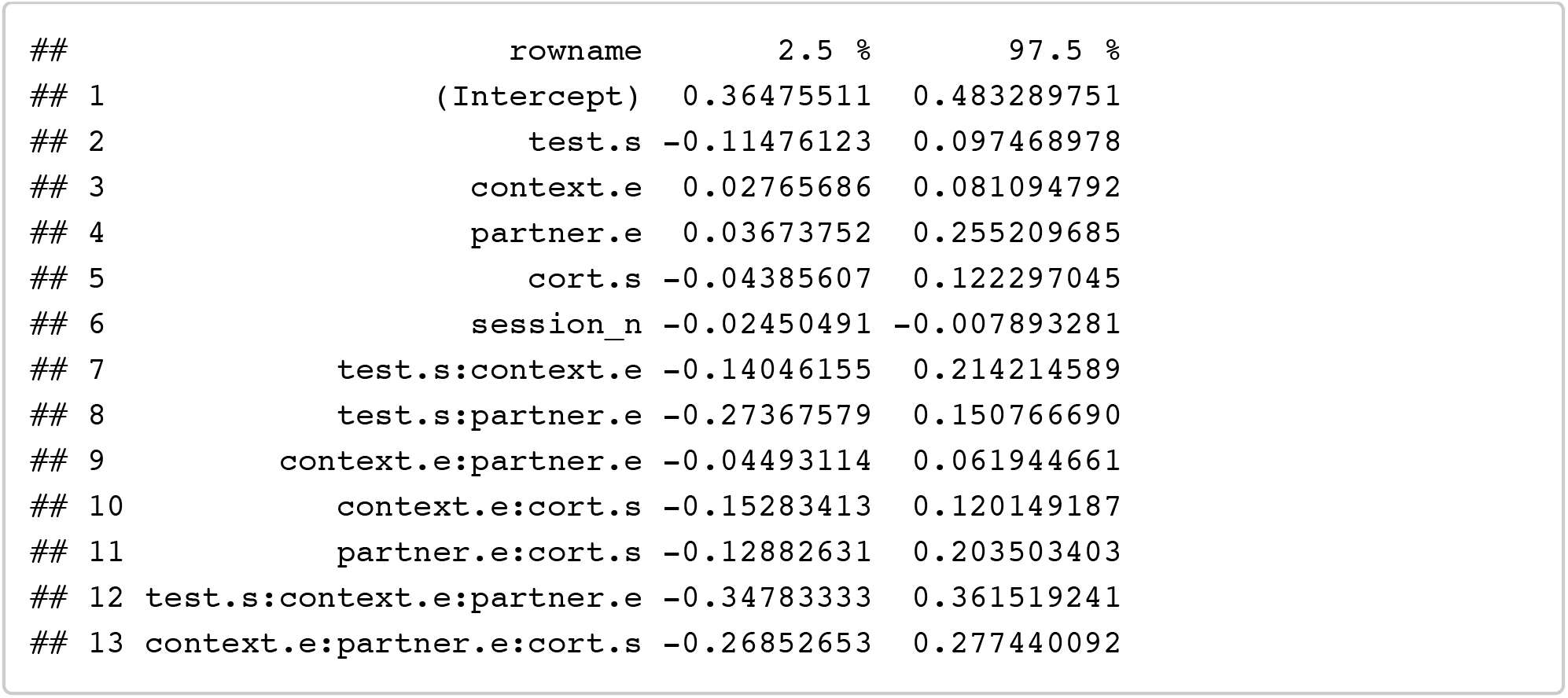

##### T + C + T*C: (+ session order, + partnership status)

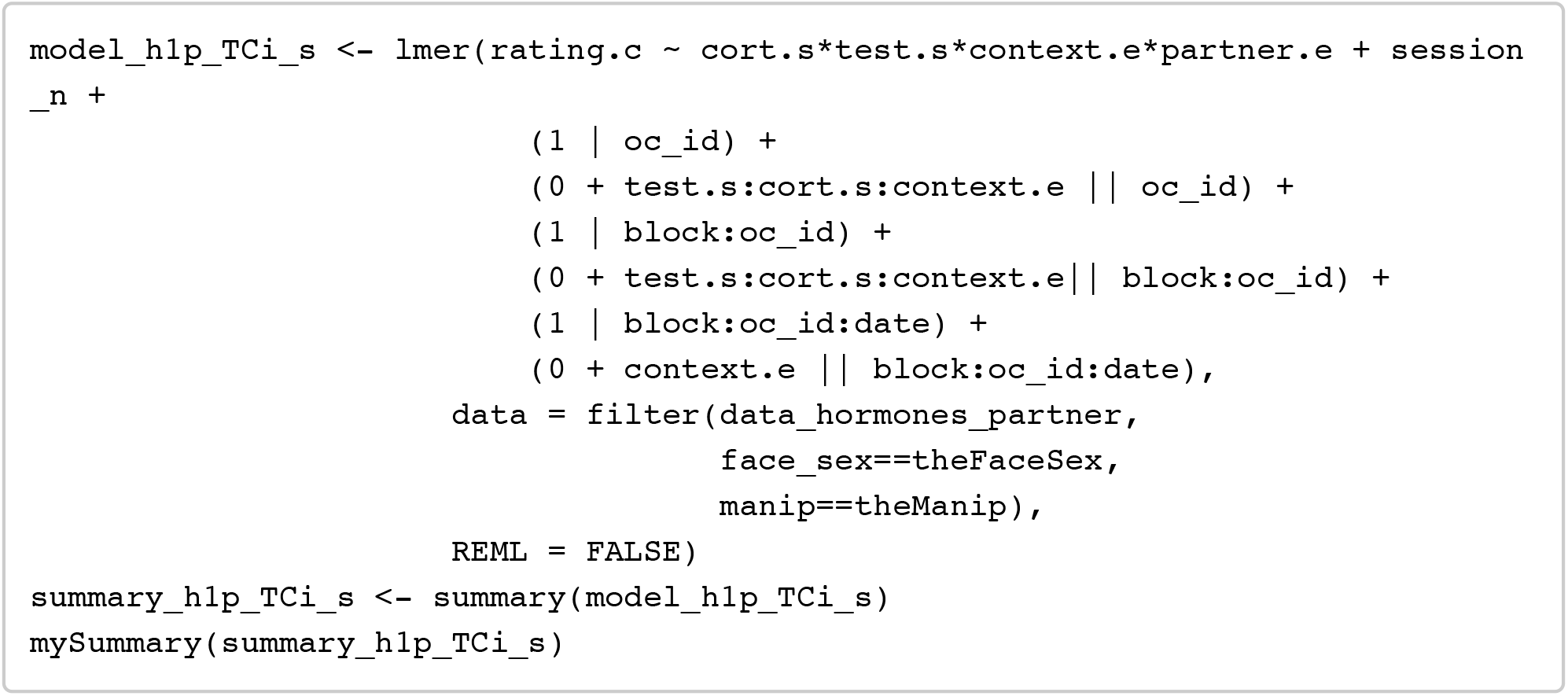

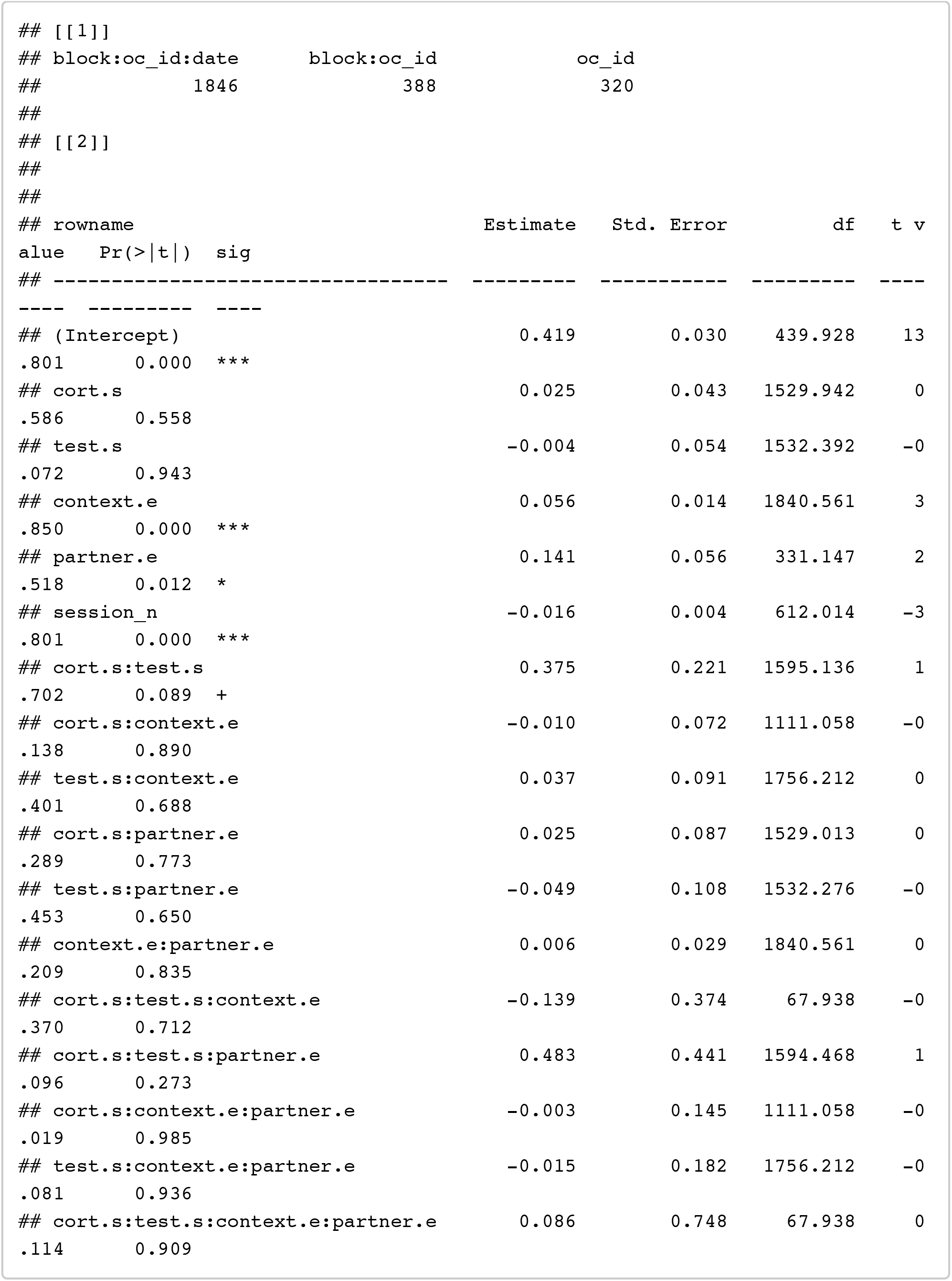

~~~
confint(model_h1p_TCi_s, method = confint_method) %>% as.data.frame() %>% rown ames_to_column() %>% filter(!is.na(‘2.5 %’))
~~~

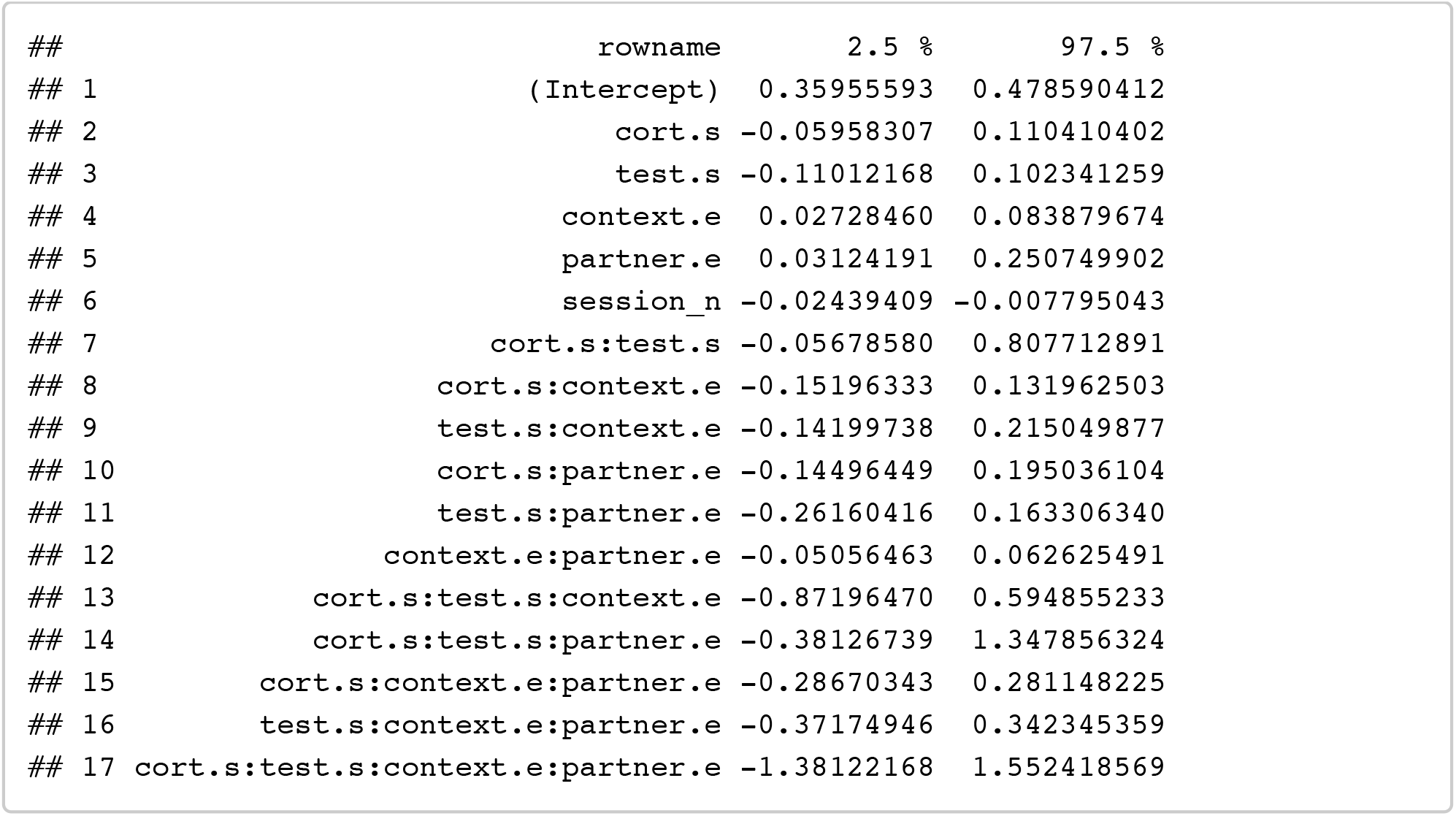

### Hypothesis 2

Do women not using hormonal contraceptives show stronger preferences than women using the combined oral contraceptive pill?

One of these women was excluded from analyses because she did not complete any male face preference tests.

#### Descriptive stats: data_between

~~~
*# create mean DV for all ratings by oc_id*
stats_overall <- filter(data_between, face_sex==theFaceSex, manip==theManip) %>%
 group_by(oc_id) %>%
 summarise(
   overall_rating.c = mean(rating.c)
 ) %>%
 ungroup() %>%
 group_by() %>%
 summarise(
   context = “overall”,
   pill.e = “overall”,
   n= n_distinct(oc_id),
   mean_dv = mean(overall_rating.c),
   sd_dv = sd(overall_rating.c),
   se_dv = se(overall_rating.c)
 )
*# create mean DV splitting by context*
stats_context <- filter(data_between, face_sex==theFaceSex, manip==theManip) %>%
 group_by(oc_id, context) %>%
 summarise(
   context_rating.c = mean(rating.c)
 ) %>%
 group_by(context) %>%
 summarise(
   pill.e = “overall”,
   n= n_distinct(oc_id),
   mean_dv = mean(context_rating.c),
   sd_dv = sd(context_rating.c),
   se_dv = se(context_rating.c)
 )
*# stats by pill.e*
stats_pill <- filter(data_between, face_sex==theFaceSex, manip==theManip) %>%
 group_by(oc_id, pill.e) %>%
 summarise(
  overall_rating.c = mean(rating.c)
 ) %>%
 ungroup() %>%
 group_by(pill.e) %>%
 summarise(
   context = “overall”,
   n= n_distinct(oc_id),
   mean_dv = mean(overall_rating.c),
   sd_dv = sd(overall_rating.c),
   se_dv = se(overall_rating.c)
 ) %>% select(context, pill.e, n, mean_dv,sd_dv, se_dv)
rbind(stats_overall, rbind(stats_context, stats_pill))
~~~

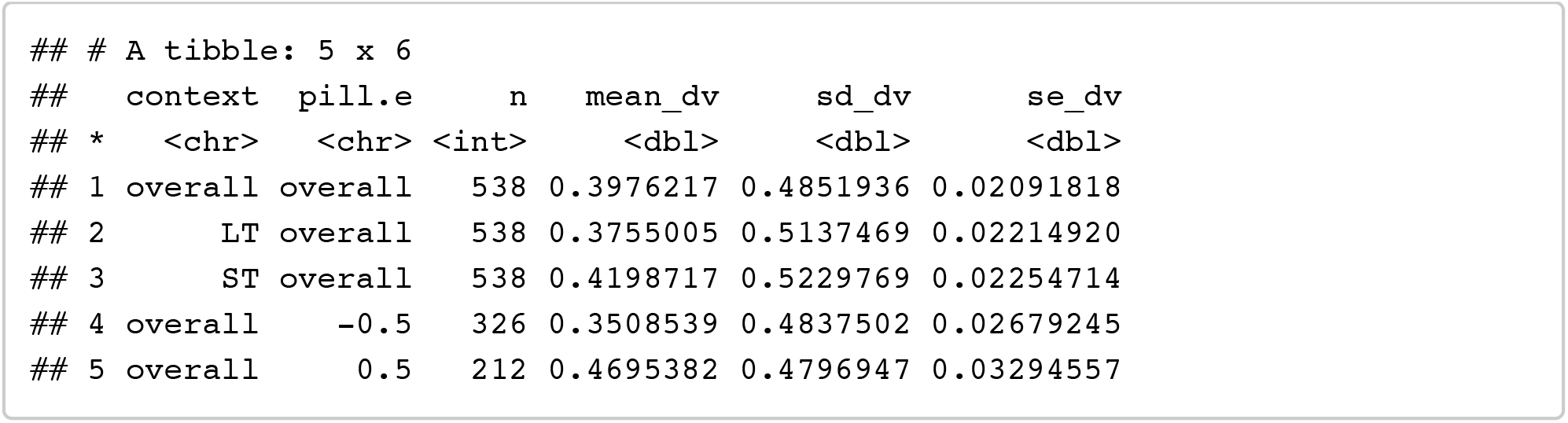

#### Analyses H2: Pill

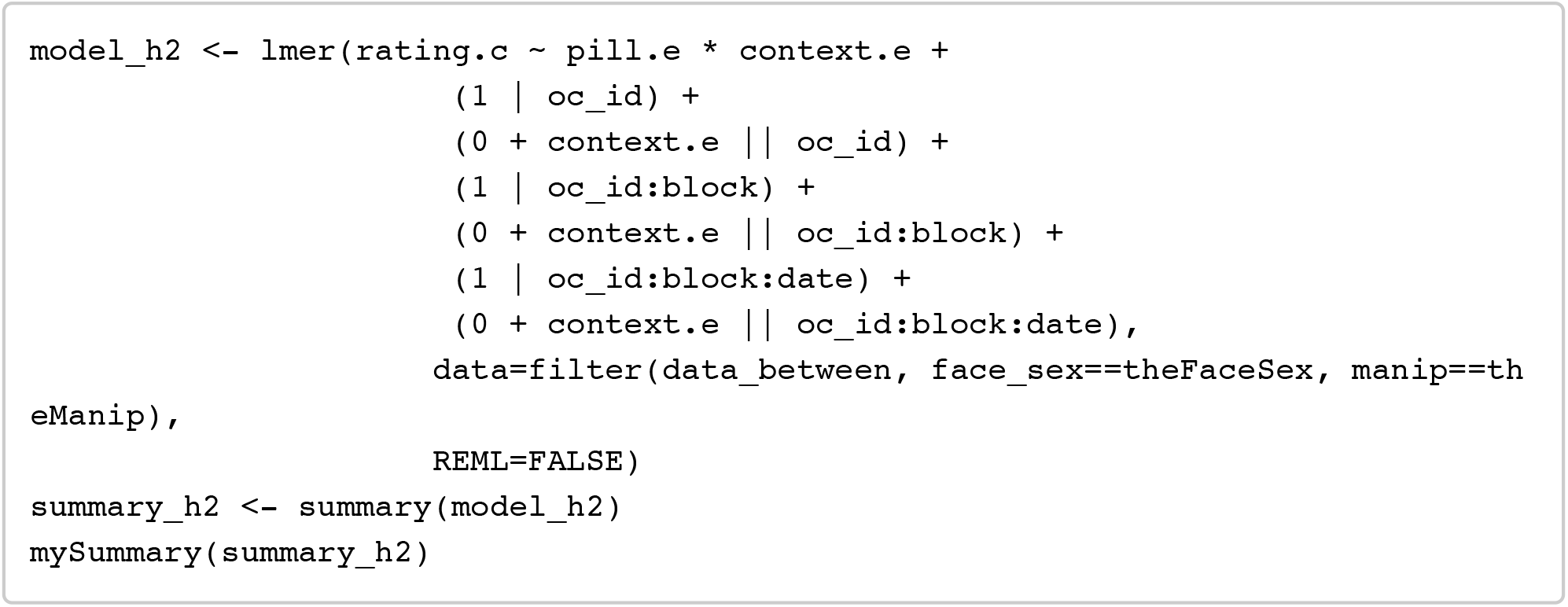

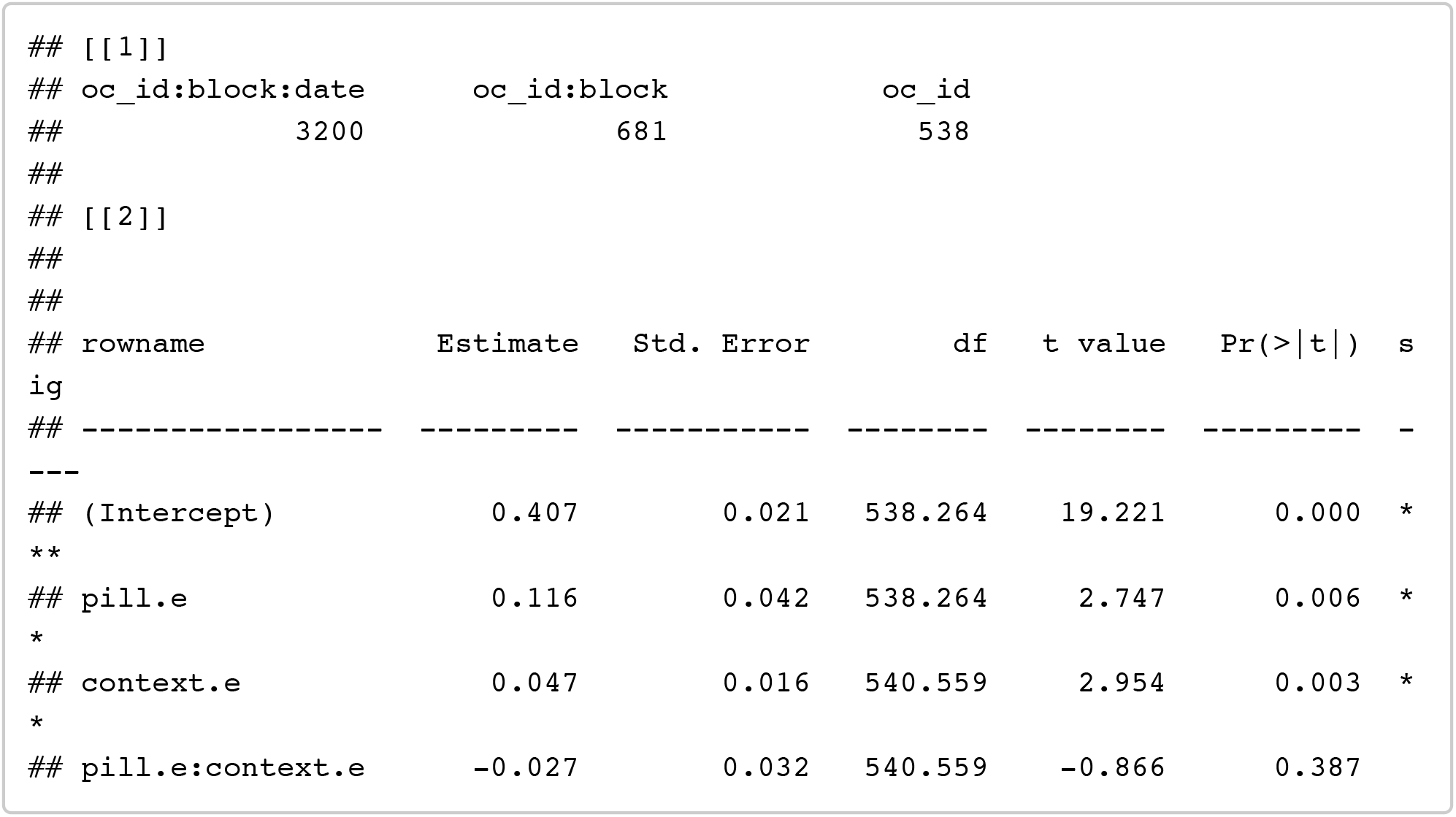

~~~
confint(model_h2, method = confint_method) %>% as.data.frame() %>% rownames_to _column() %>% filter(!is.na(‘2.5 %’))
~~~

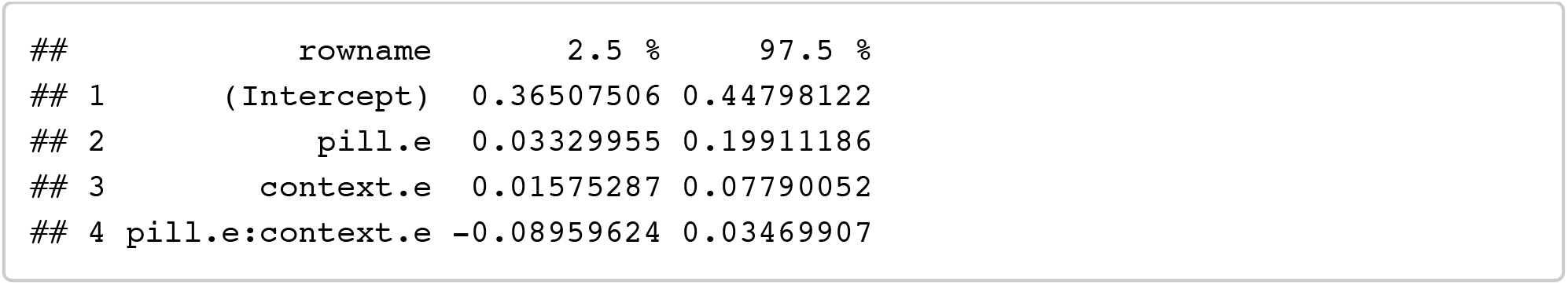

Note that the effect of OCP use (estimate = +0.116, p = .006) is in the opposite direction to what would be expected if fertility is positively associated with preference.

#### Analyses H2p: Pill (+ partnership status)

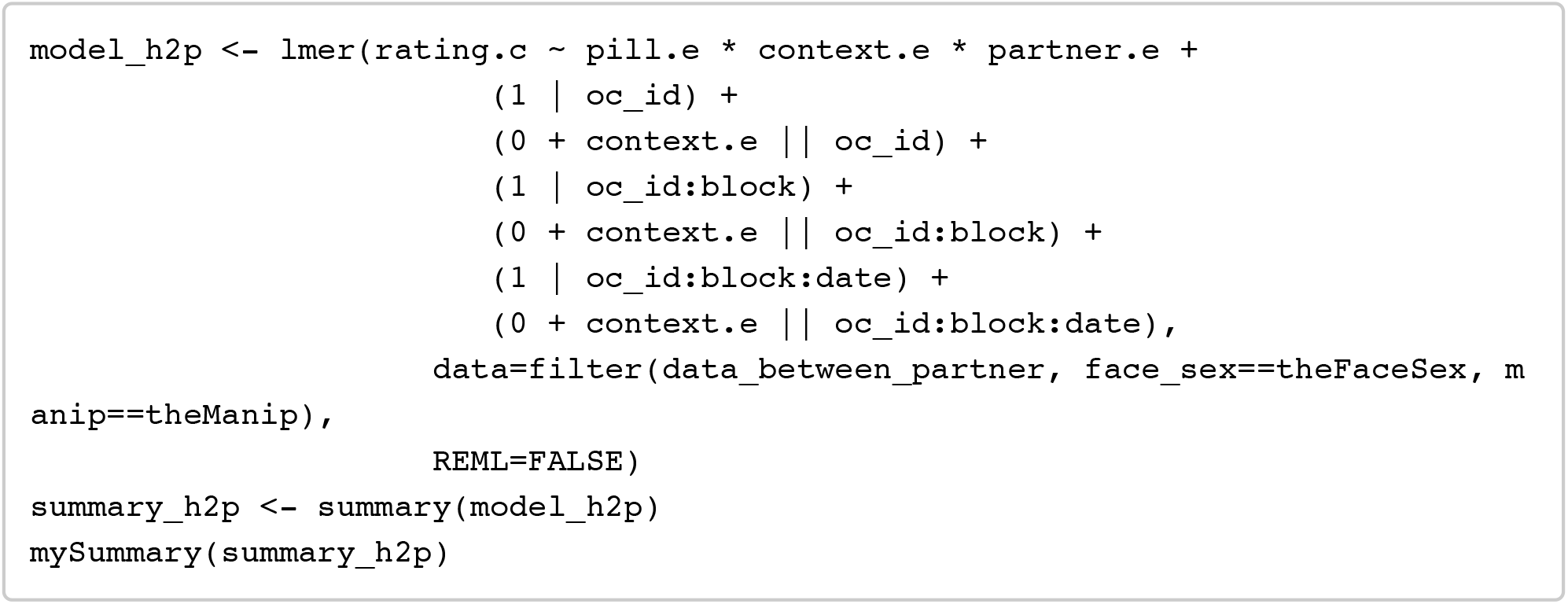

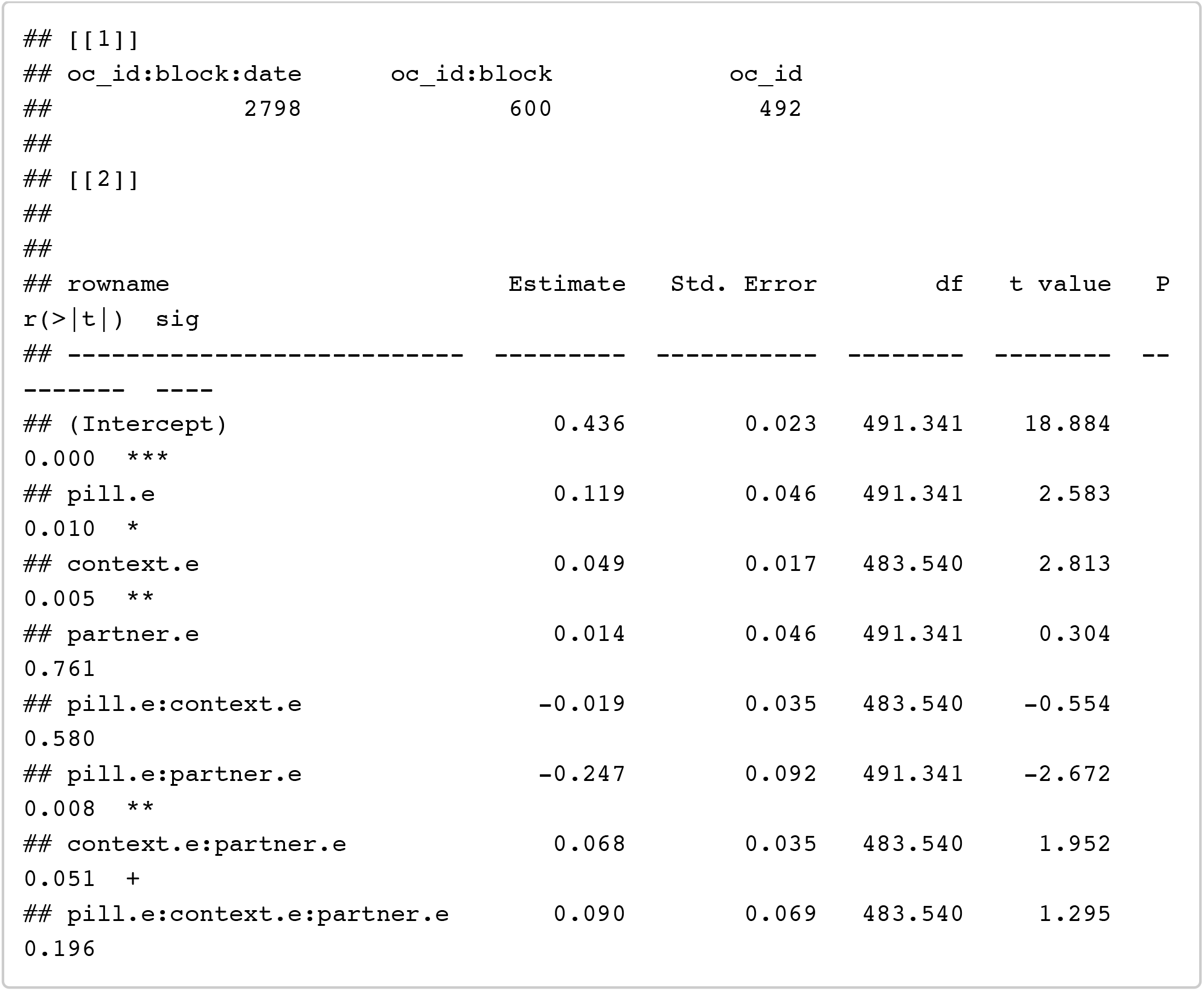

~~~
confint(model_h2p, method = confint_method) %>% as.data.frame() %>% rownames_t o_column() %>% filter(!is.na(‘2.5 %’))
~~~

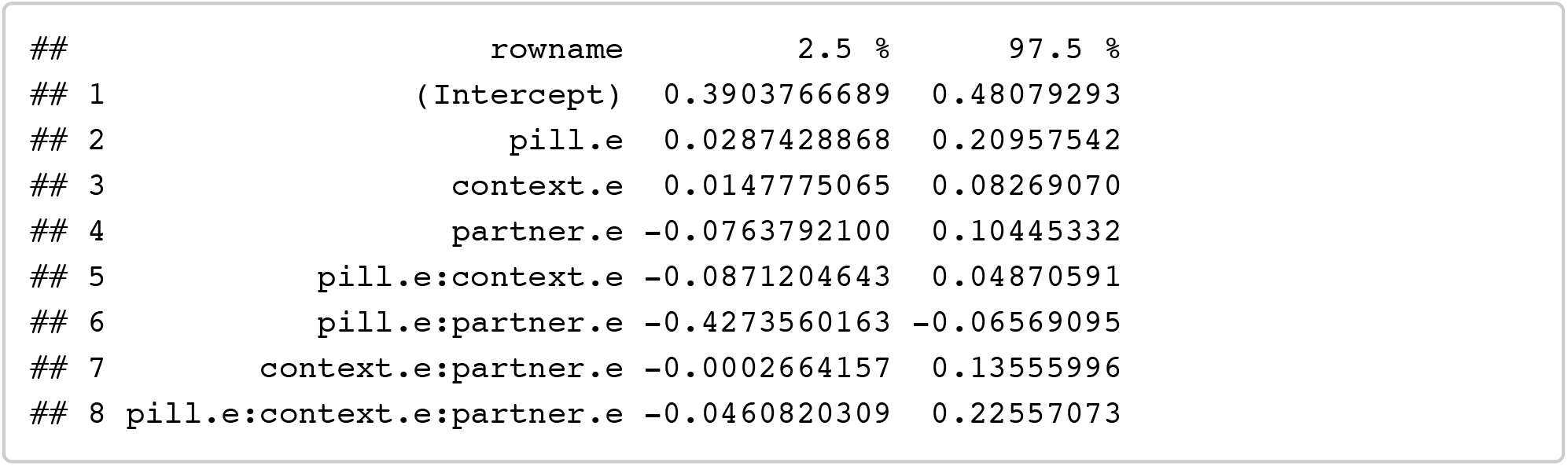

##### Pill (single women only to interpret interaction)

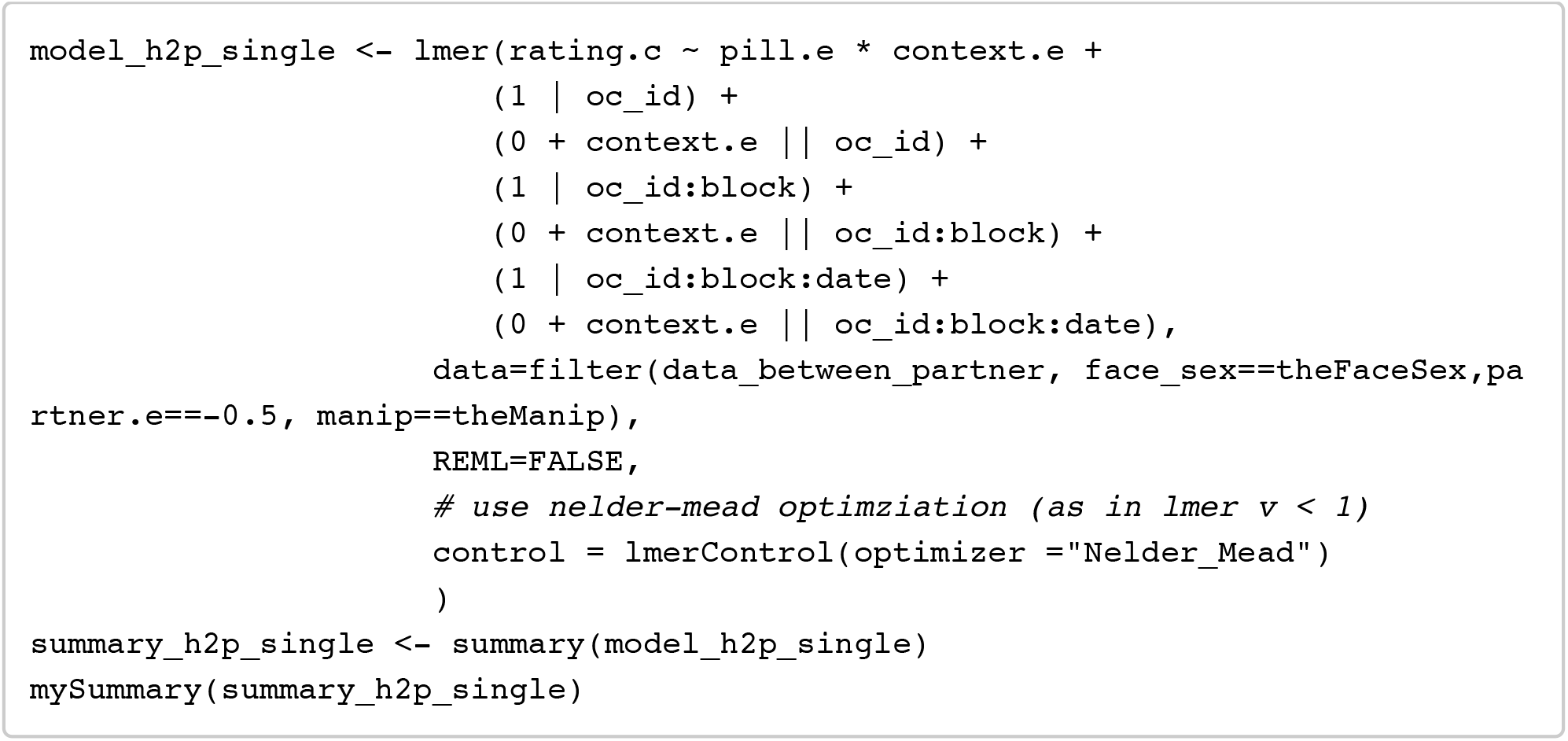

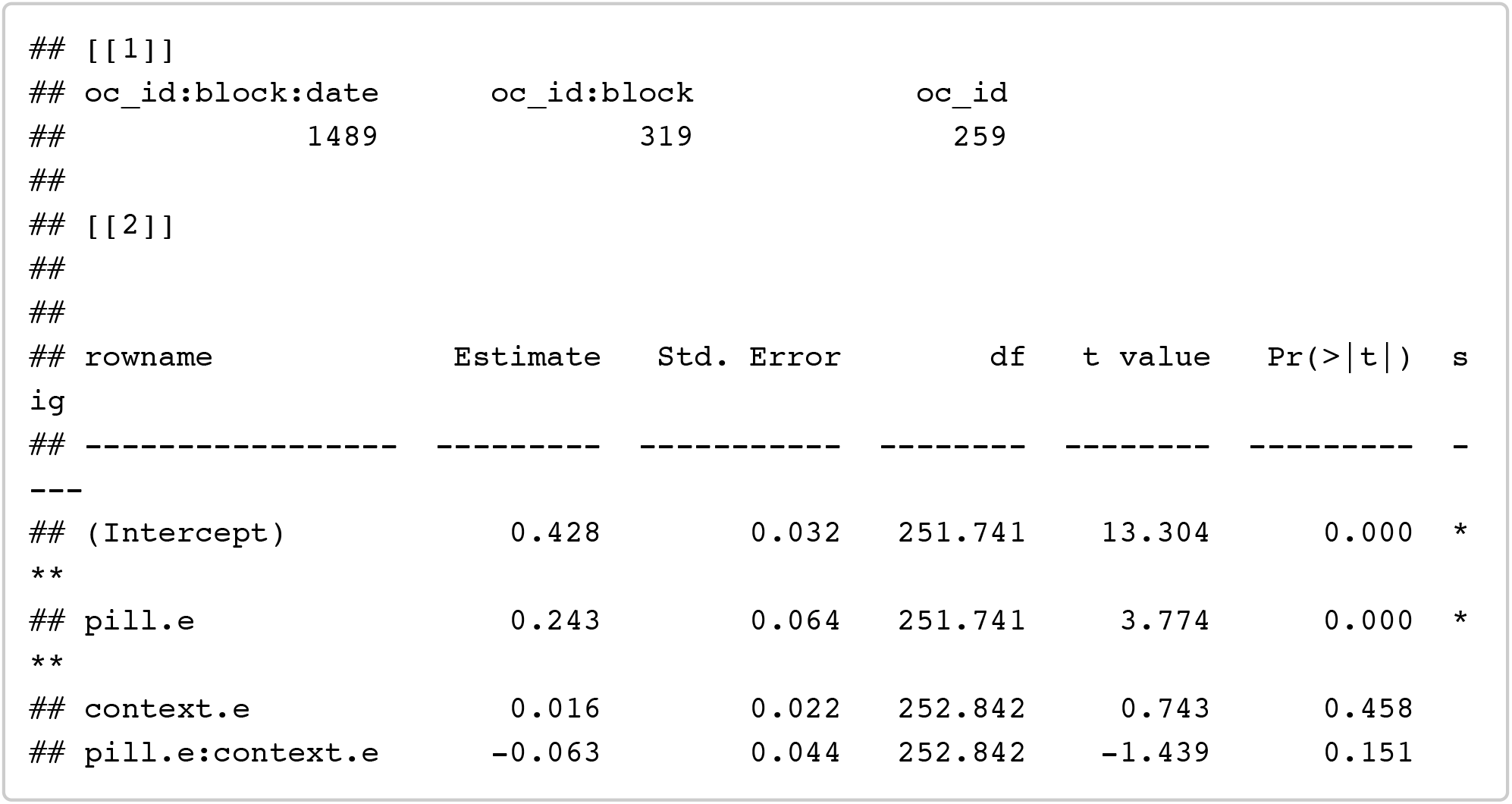

~~~
confint(model_h2p_single, method = confint_method) %>% as.data.frame() %>% row names_to_column() %>% filter(!is.na(‘2.5 %’))
~~~

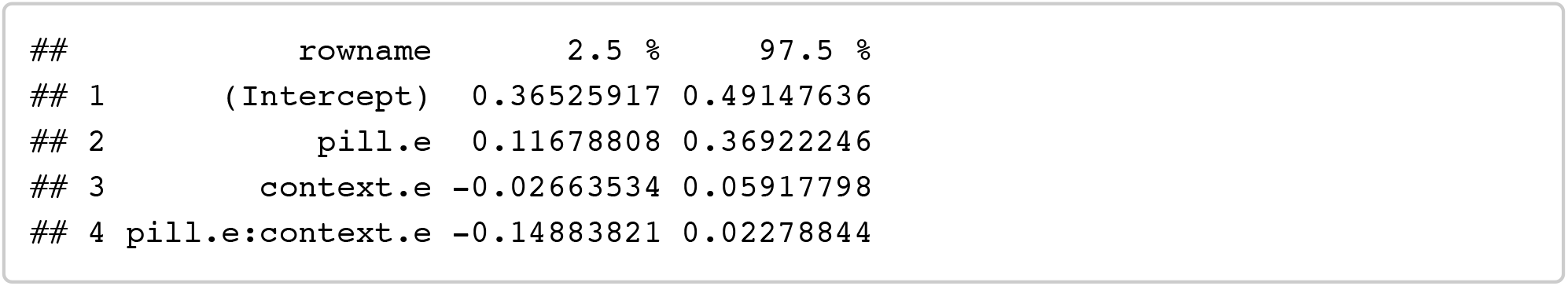

##### Pill (partnered women only to interpret interaction)

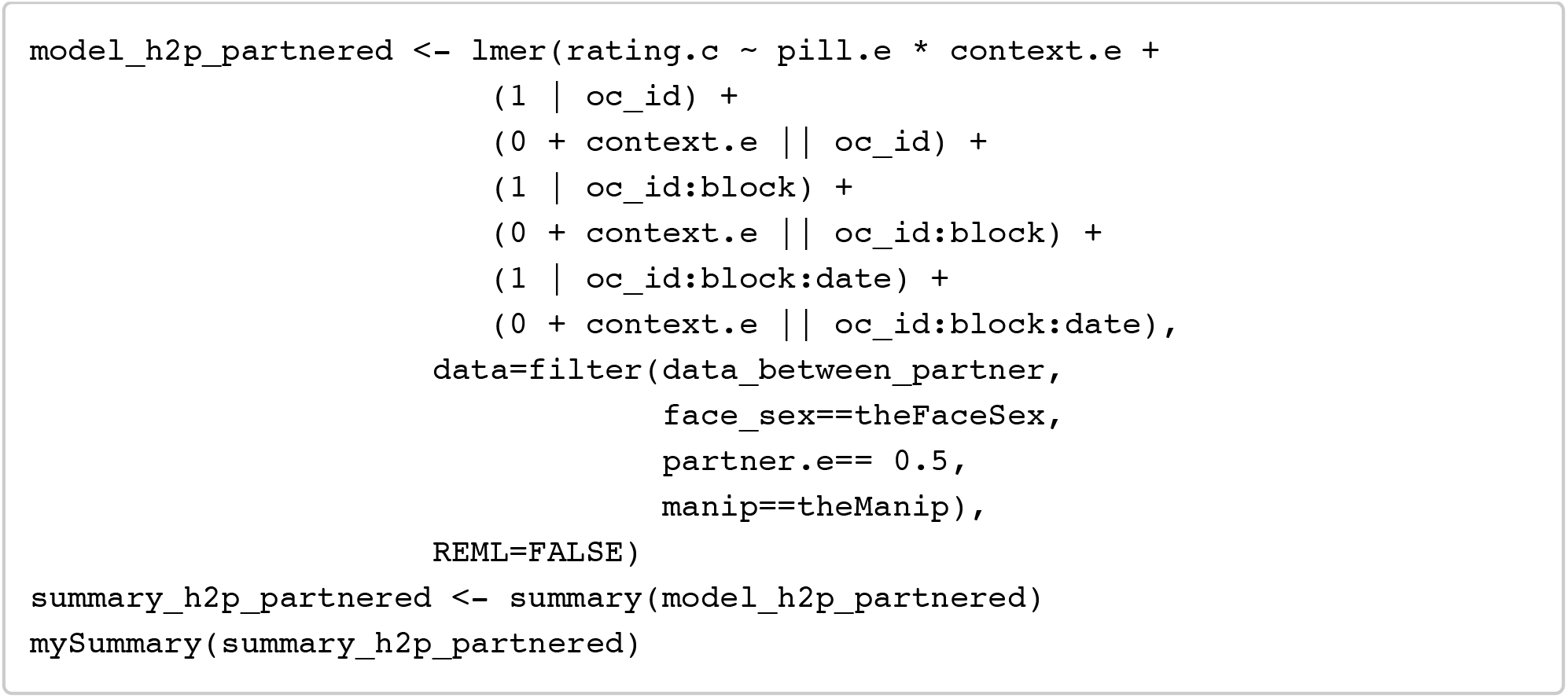

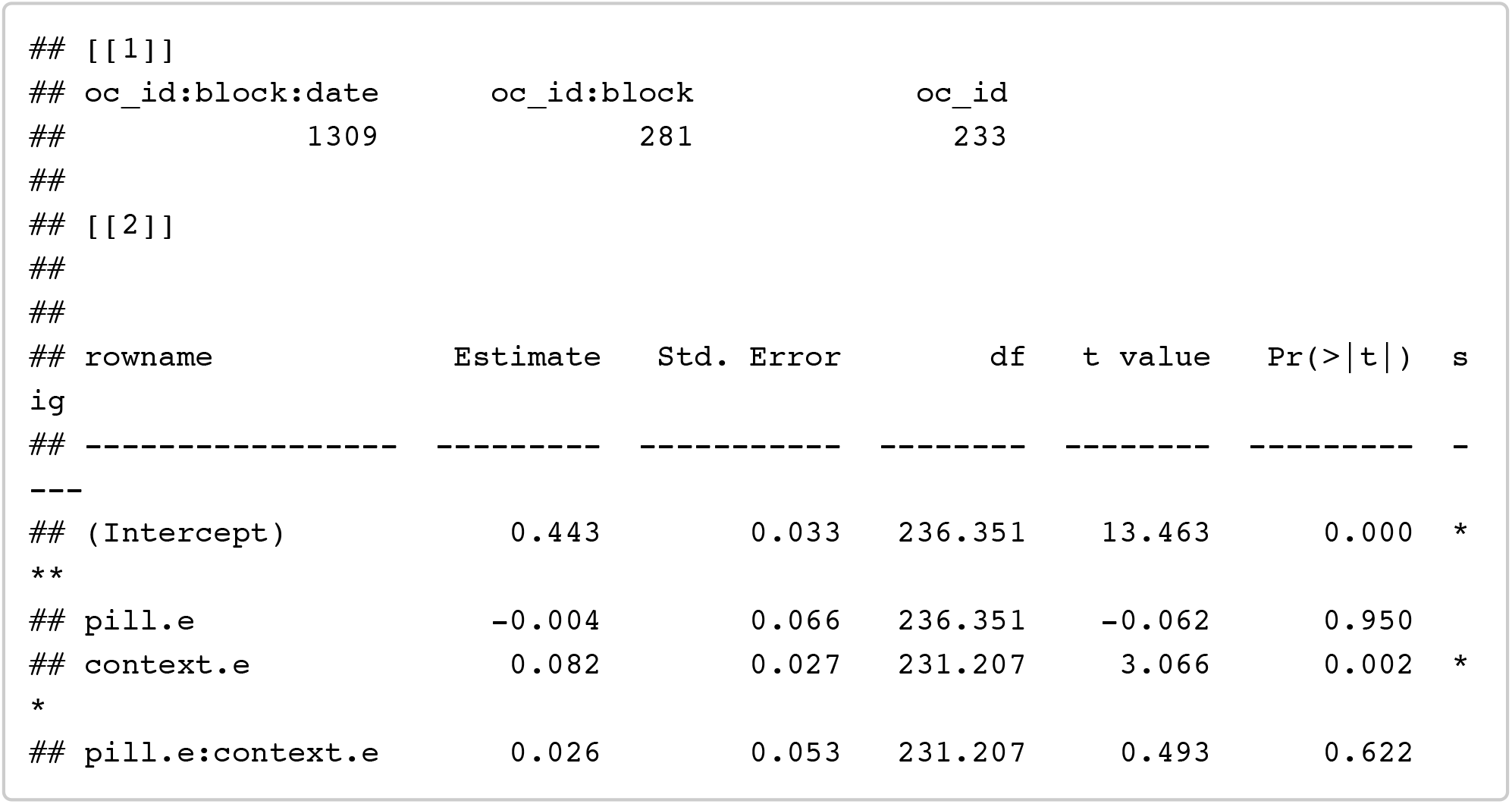

~~~
confint(model_h2p_partnered, method = confint_method) %>% as.data.frame() %>% rownames_to_column() %>% filter(!is.na(‘2.5 %’))
~~~

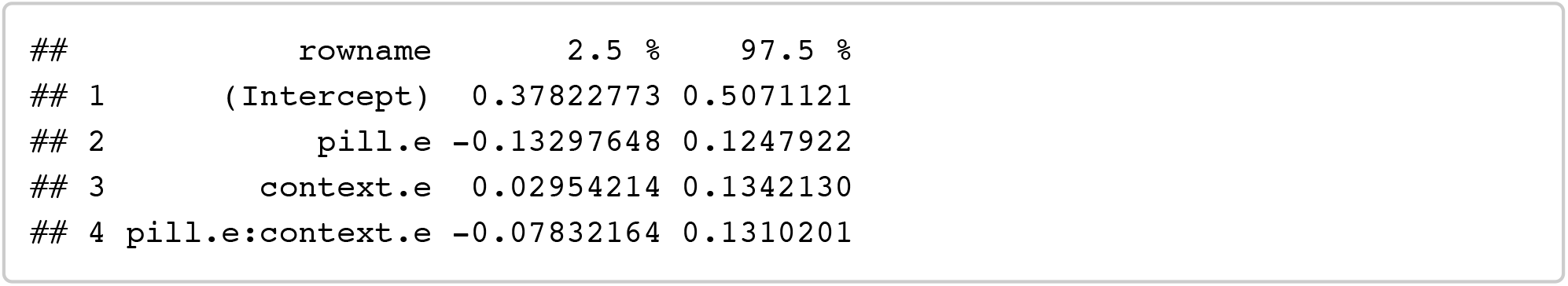

### Hypothesis 3

Do preferences of women using the combined oral contraceptive pill change when they are taking inactive pills?

pill_break.e coding: using active pill= 0.5, using inactive pill/being on on a pill break = -0.5

#### Descriptive stats: data_pillbreak

~~~
*# create mean DV for all ratings by oc_id*
stats_overall <- filter(data_pillbreak, face_sex==theFaceSex, manip==theManip) %>%
 group_by(oc_id) %>%
 summarise(
   overall_rating.c = mean(rating.c)
 ) %>%
   ungroup() %>%
   group_by() %>%
   summarise(
      context = “overall”,
      n= n_distinct(oc_id),
      mean_dv = mean(overall_rating.c),
      sd_dv = sd(overall_rating.c),
      se_dv = se(overall_rating.c)
 )
*# create mean DV splitting by context*
stats_context <- filter(data_pillbreak, face_sex==theFaceSex, manip==theManip) %>%
 group_by(oc_id, context) %>%
 summarise(
    context_rating.c = mean(rating.c)
 ) %>%
 group_by(context) %>%
 summarise(
    n= n_distinct(oc_id),
    mean_dv = mean(context_rating.c),
    sd_dv = sd(context_rating.c),
    se_dv = se(context_rating.c)
 )
rbind(stats_overall, stats_context)
~~~

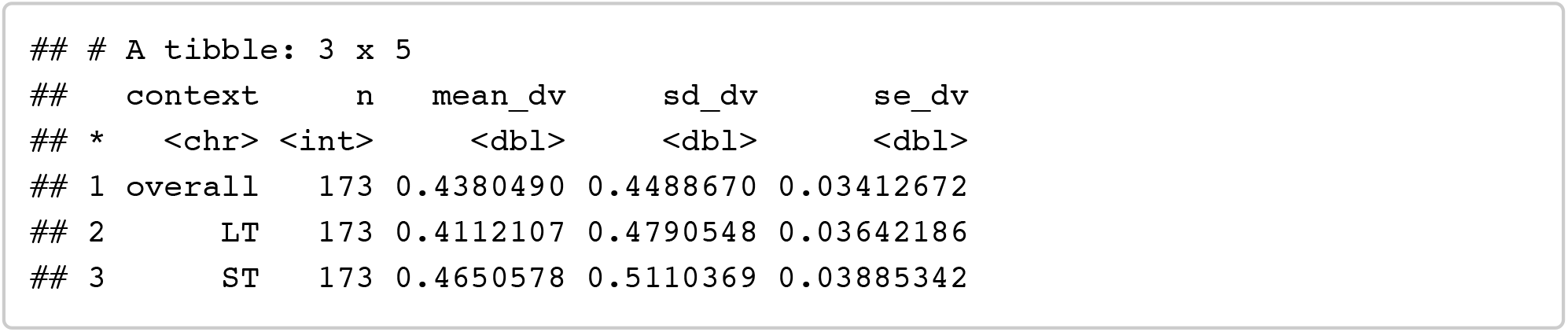

#### Analyses H3: Pill-break

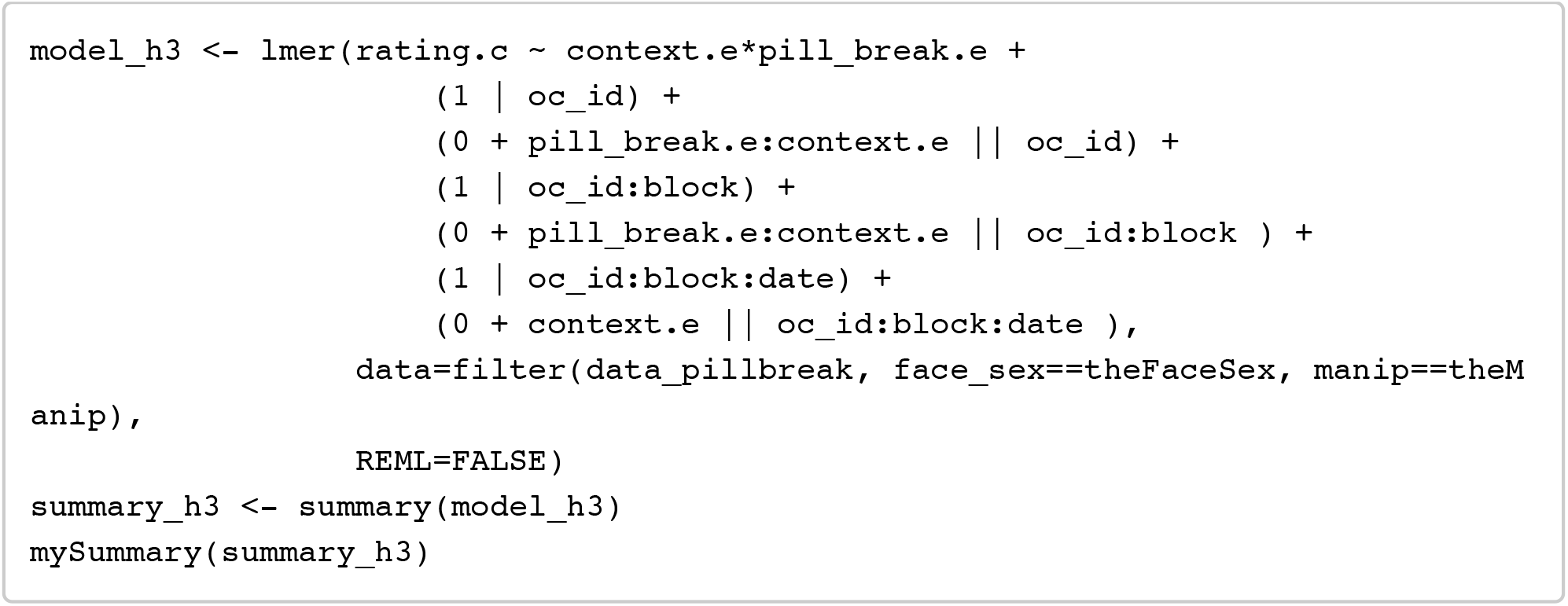

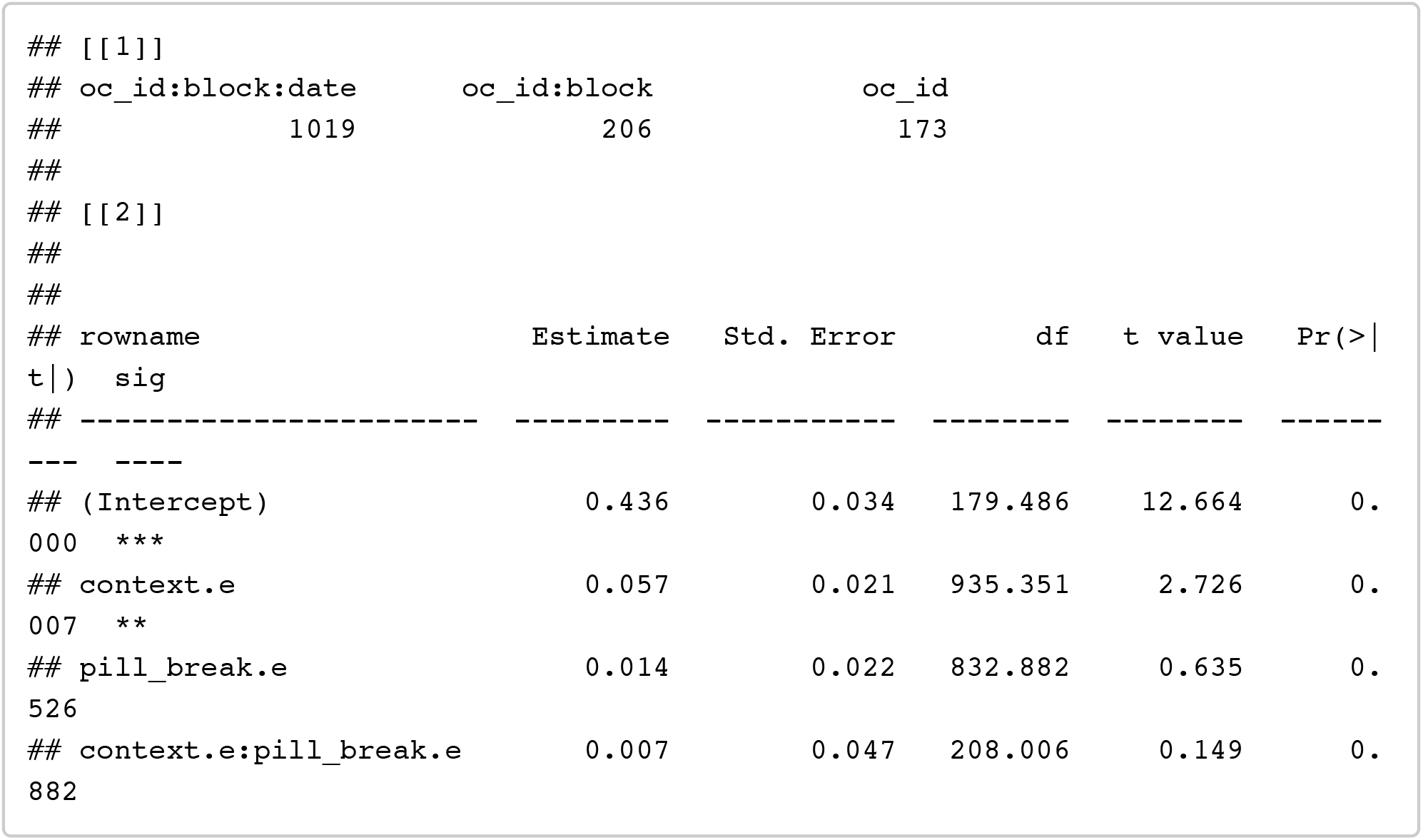

~~~
confint(model_h3, method = confint_method) %>% as.data.frame() %>% rownames_to _column() %>% filter(!is.na(‘2.5 %’))
~~~

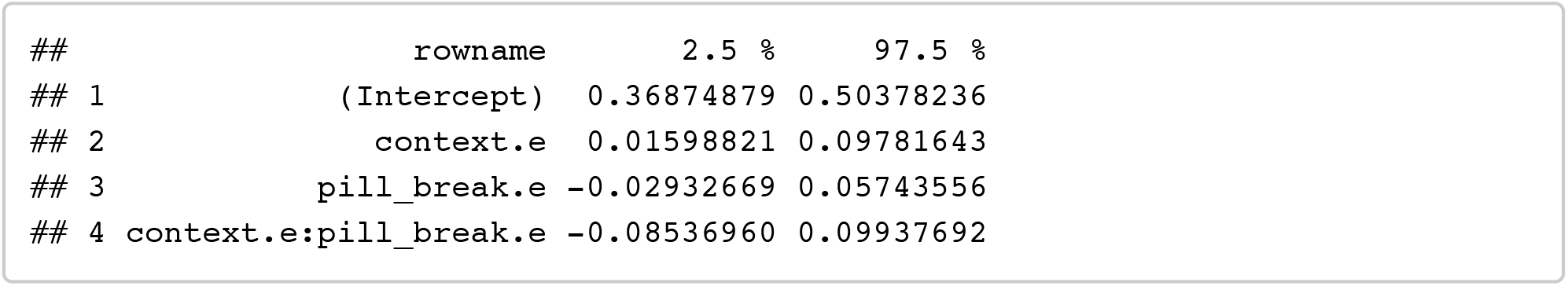

#### Analyses H3p: Pill-break (+ partnership status)

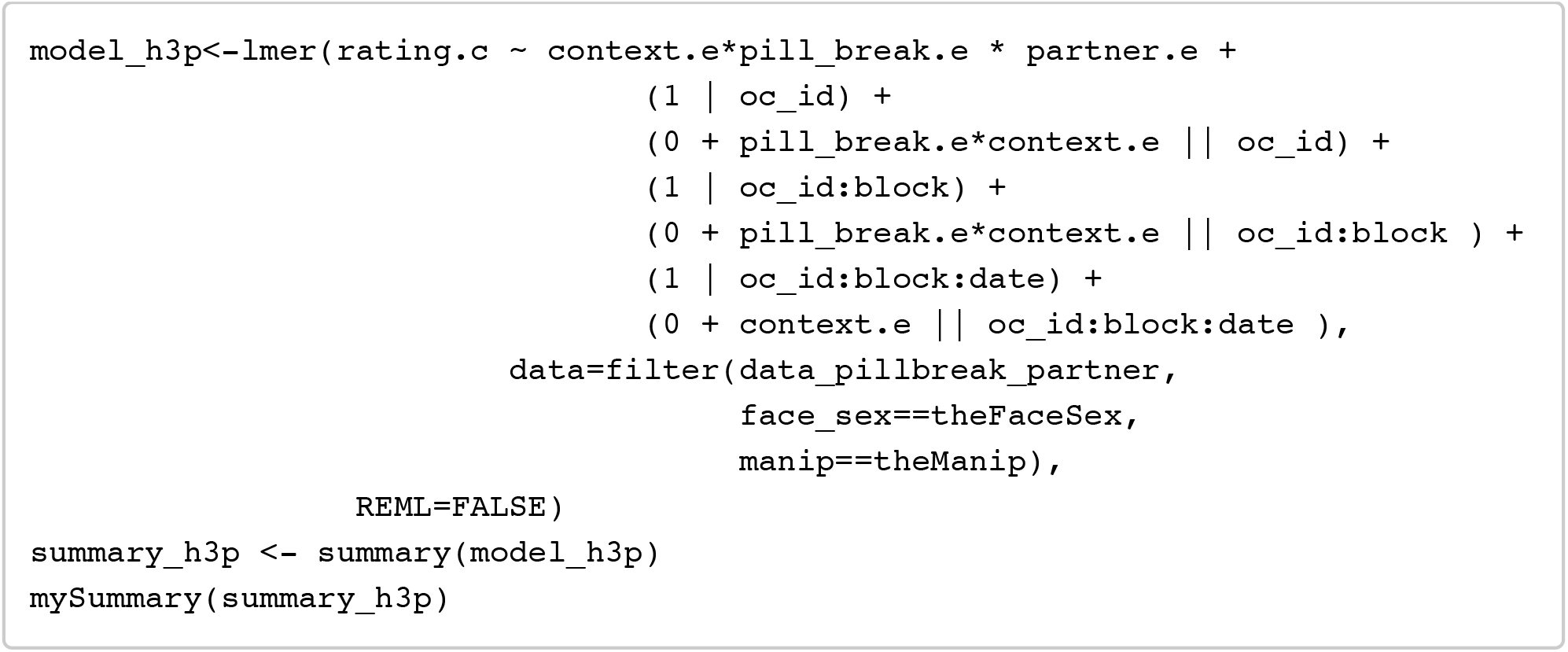

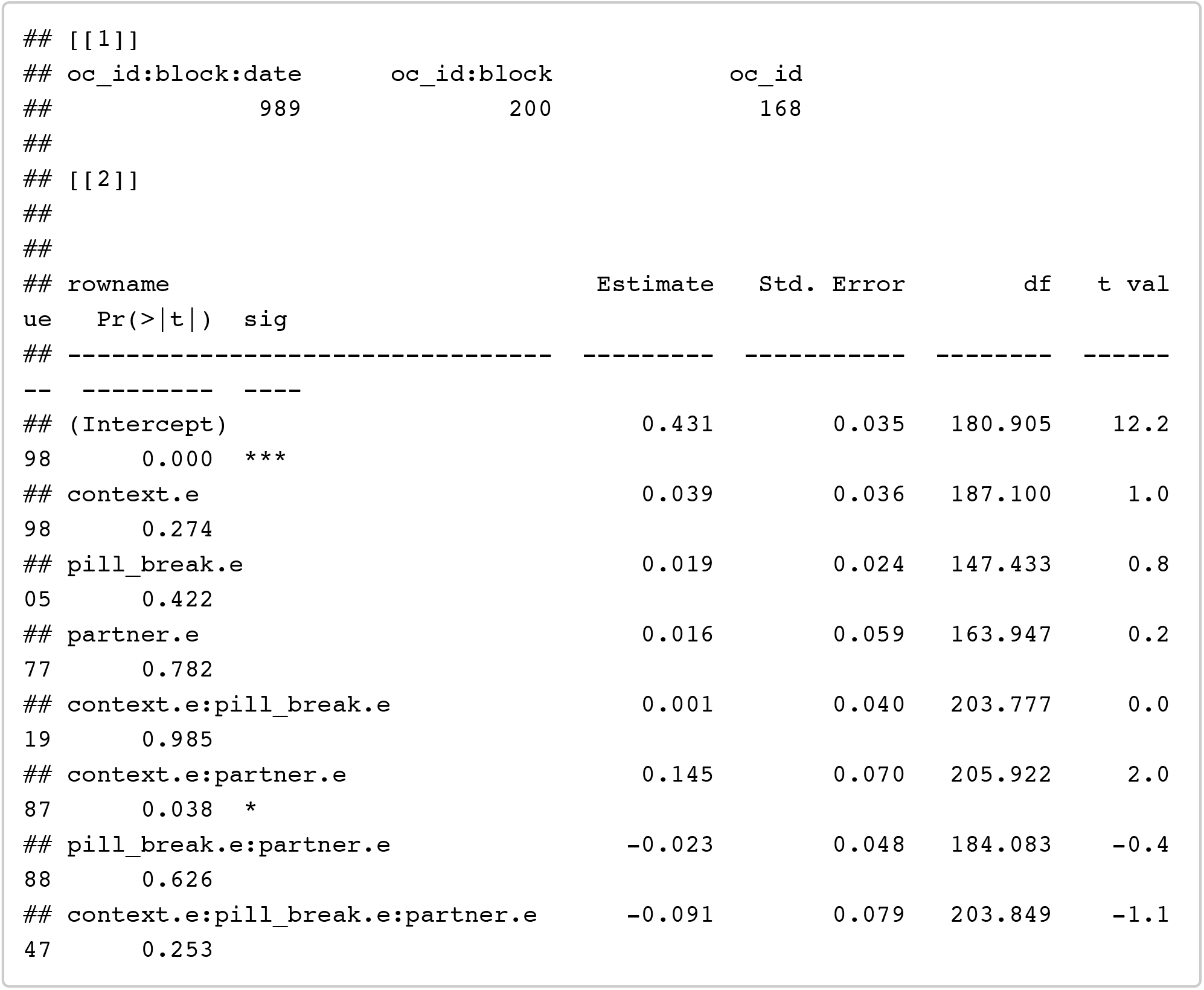

~~~
confint(model_h3p, method = confint_method) %>% as.data.frame() %>% rownames_t o_column() %>% filter(!is.na(‘2.5 %’))
~~~

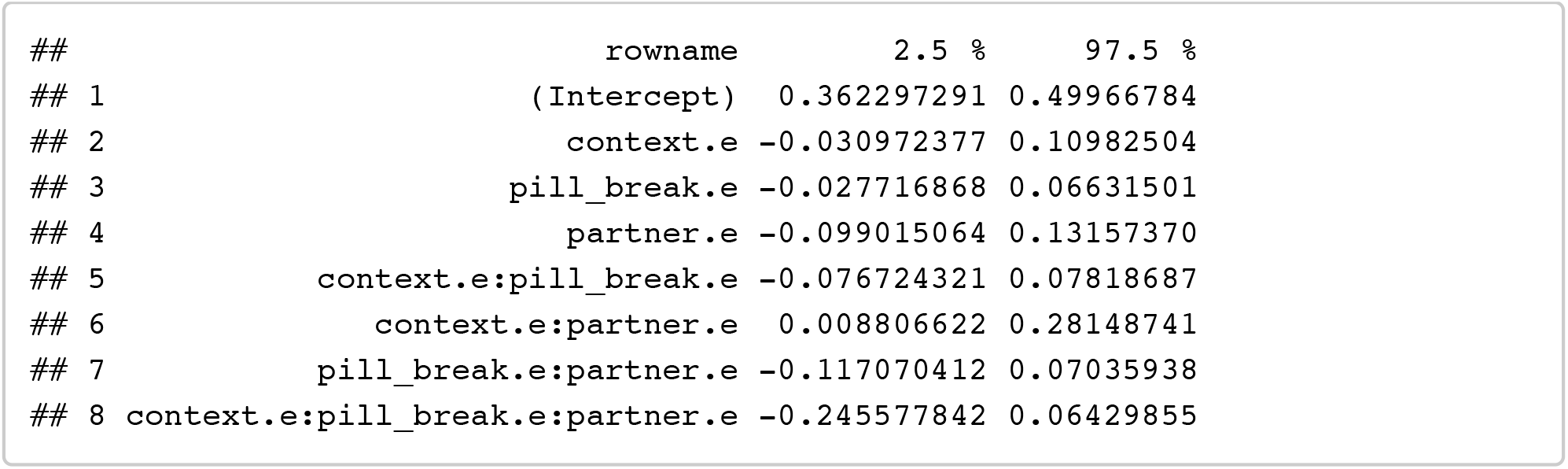

#### Pill break (single women only to interpret interaction)

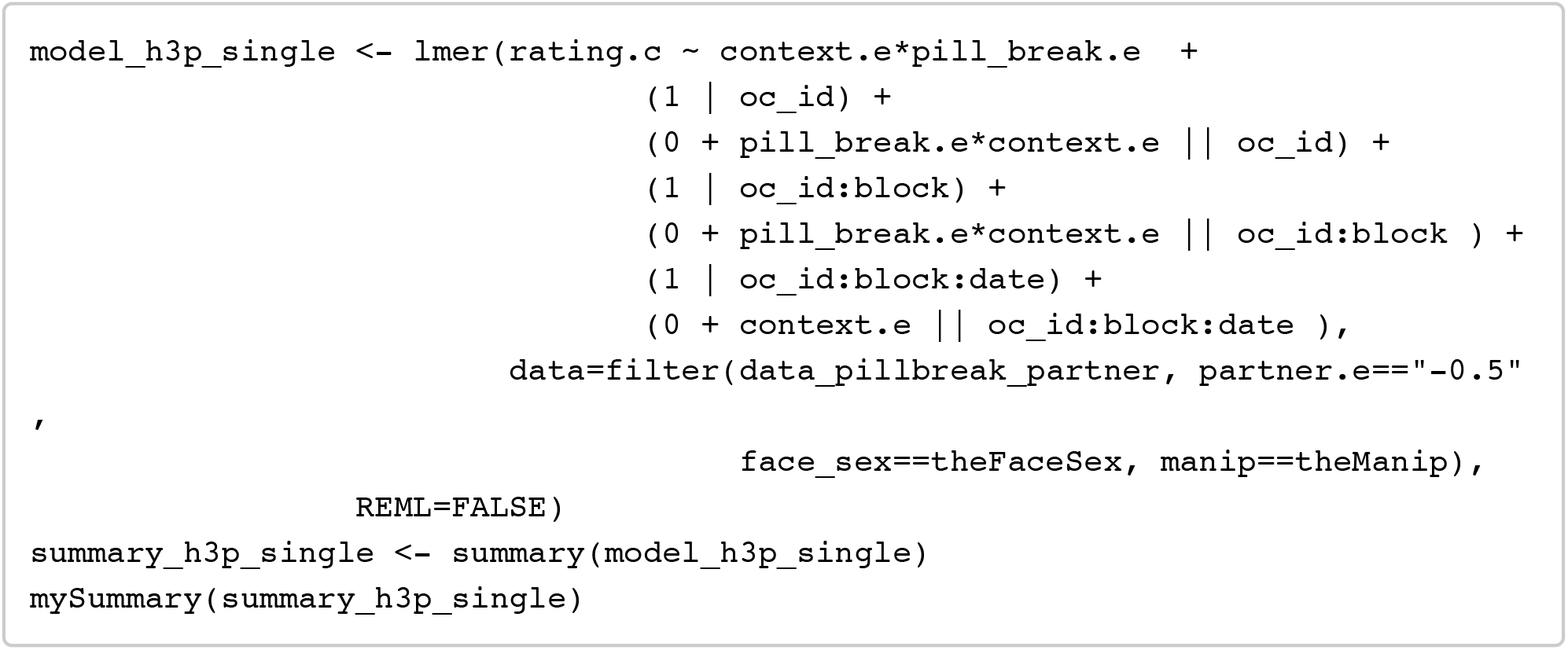

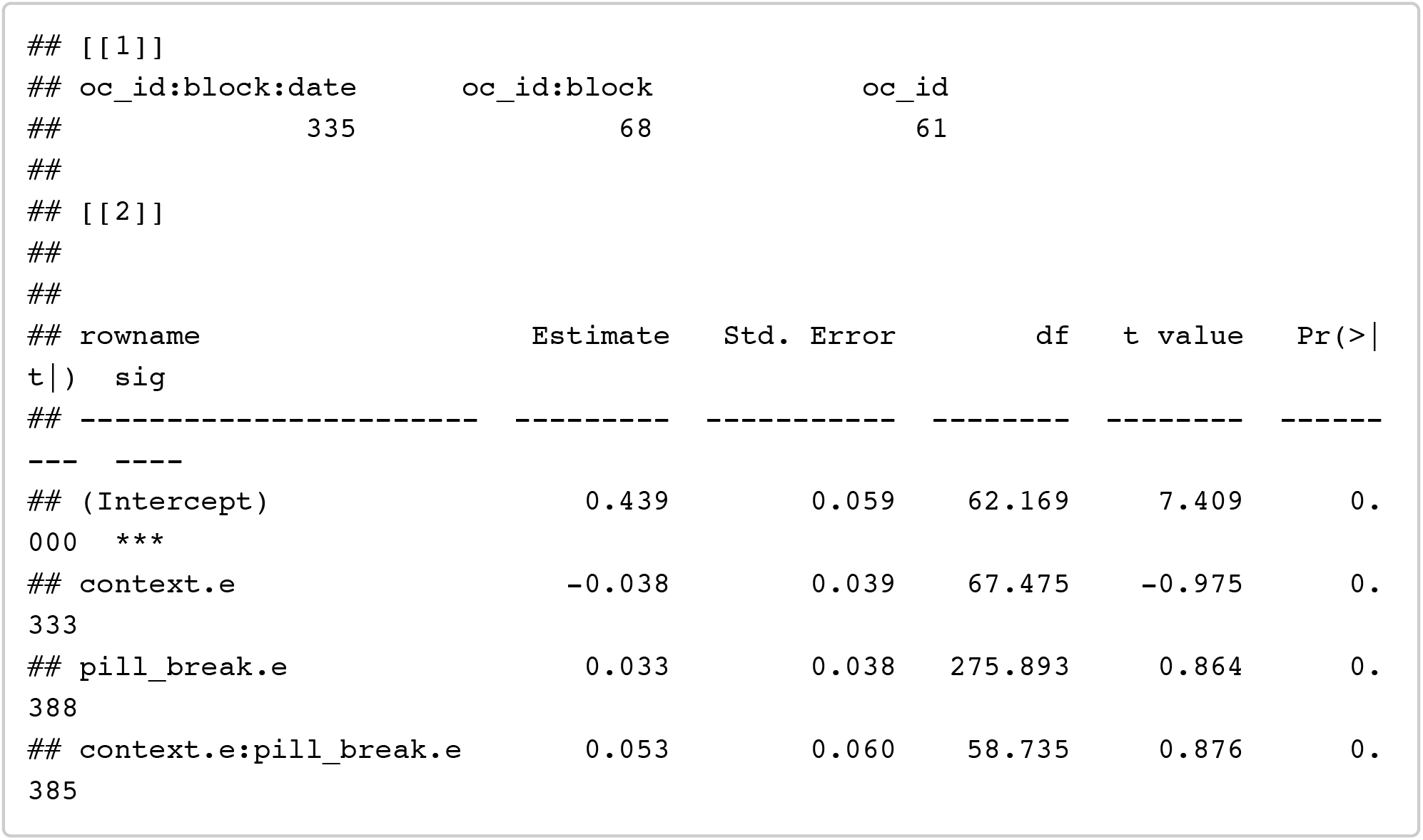

~~~
confint(model_h3p_single, method = confint_method) %>% as.data.frame() %>% row names_to_column() %>% filter(!is.na(‘2.5 %’))
~~~

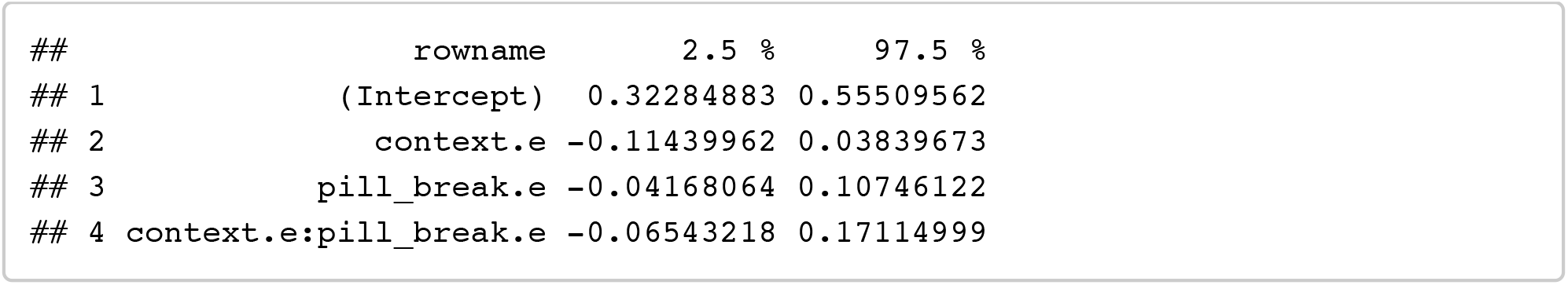

#### Pill break (partnered women only to interpret interaction)

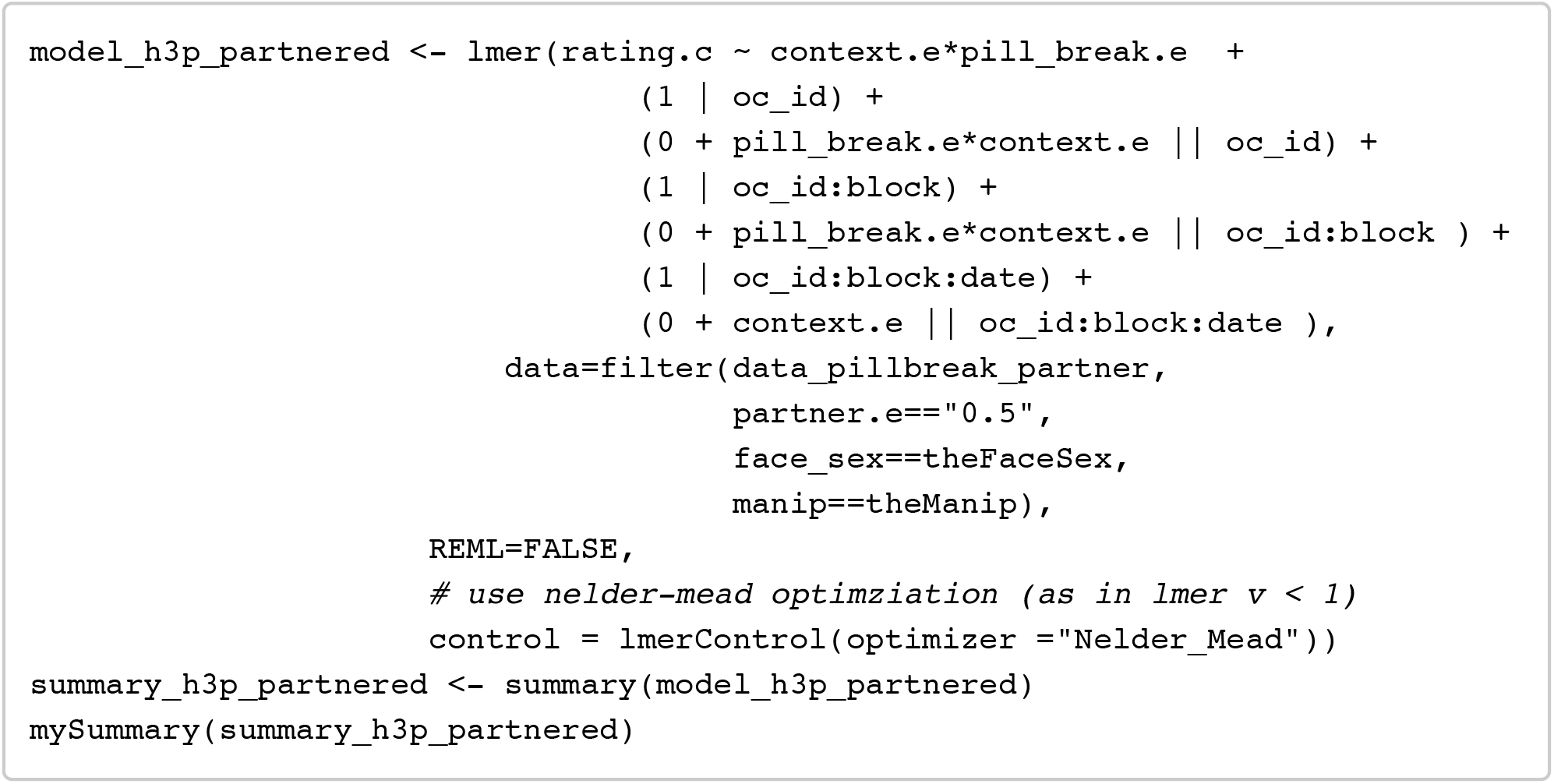

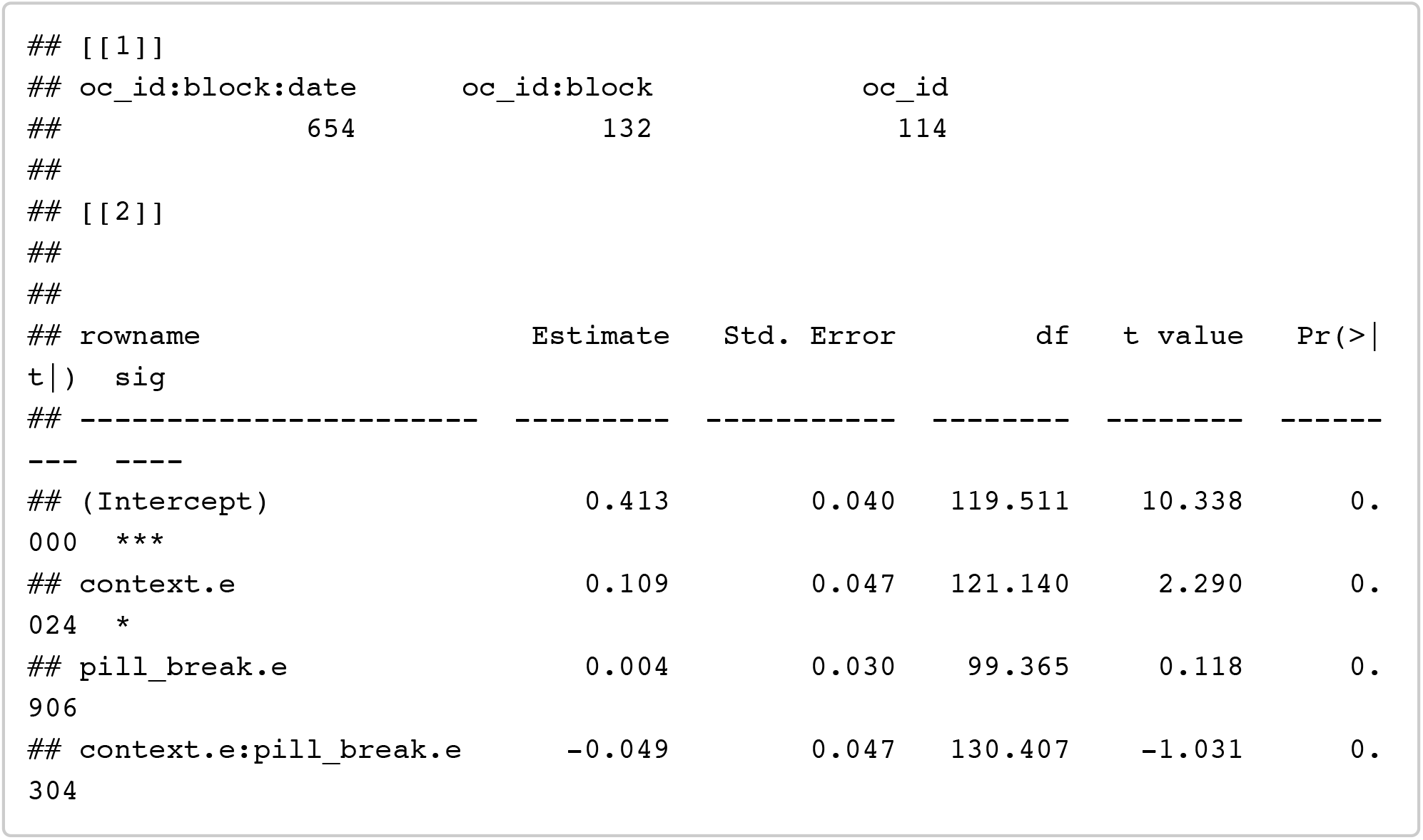

~~~
confint(model_h3p_partnered, method = confint_method) %>% as.data.frame() %>% rownames_to_column() %>% filter(!is.na(‘2.5 %’))
~~~

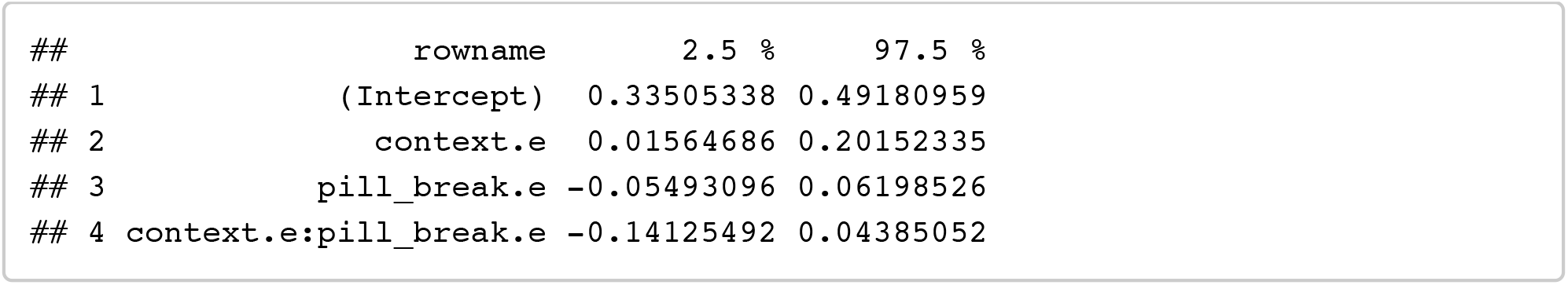

### Hypothesis 4

Do preferences change when women start or stop using the combined oral contraceptive pill?

#### Descriptive stats: data_pill_switchers

~~~
*# create mean DV for all ratings by oc_id*
stats_overall <- filter(data_pill_switchers, face_sex==theFaceSex, manip==theM anip) %>%
 group_by(oc_id) %>%
 summarise(
    overall_rating.c = mean(rating.c)
 ) %>%
 ungroup() %>%
 group_by() %>%
 summarise(
    context = “overall”,
    n= n_distinct(oc_id),
    mean_dv = mean(overall_rating.c),
    sd_dv = sd(overall_rating.c),
    se_dv = se(overall_rating.c)
 )
*# create mean DV splitting by context*
stats_context <- filter(data_pill_switchers, face_sex==theFaceSex, manip==theM anip) %>%
 group_by(oc_id, context) %>%
 summarise(
    context_rating.c = mean(rating.c)
 ) %>%
 group_by(context) %>%
 summarise(
    n= n_distinct(oc_id),
    mean_dv = mean(context_rating.c),
    sd_dv = sd(context_rating.c),
    se_dv = se(context_rating.c)
 )
rbind(stats_overall, stats_context)
~~~

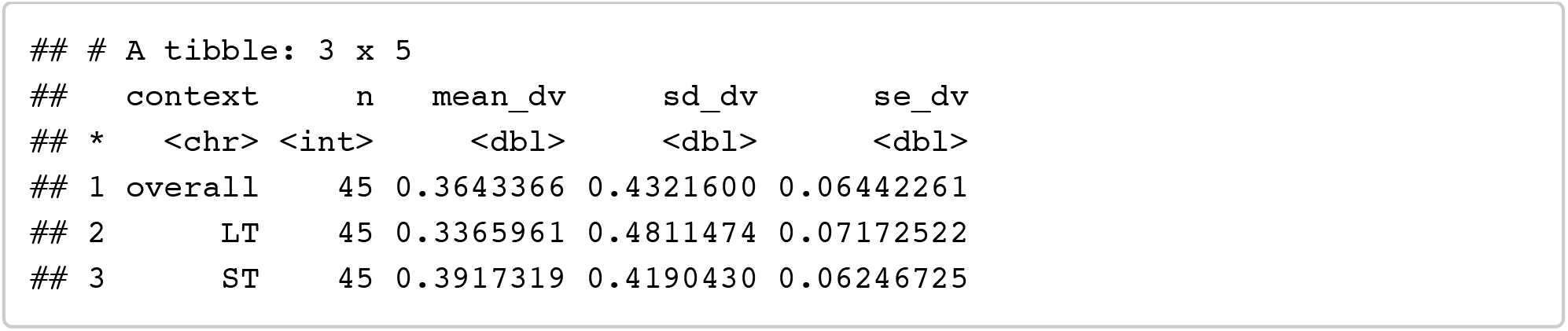

#### Interval between pill use and non-use testing blocks

~~~
switchers_date_diffs <- filter(data_pill_switchers, face_sex==theFaceSex, mani p==theManip) %>%
 group_by(oc_id, block_pill, direction.e) %>%
 summarise(
    min_date = min(date),
    max_date = max(date),
    the_date = ifelse(
        mean(direction.e) == -.5,
        ifelse(block_pill == 0, max_date, min_date),
        ifelse(block_pill == 1, max_date, min_date)
       )
 ) %>%
 ungroup() %>%
 mutate(block_pill = paste0(“hc”, block_pill)) %>%
 select(oc_id, block_pill, the_date) %>%
 spread(block_pill, the_date) %>%
 mutate(date_diff = abs(interval(ymd(hc0), ymd(hc1)) / ddays(1)))
switchers_date_diffs %>%
 group_by() %>%
 summarise(
    mean = mean(date_diff),
    sd = sd(date_diff),
    se = se(date_diff),
    min = min(date_diff),
    max = max(date_diff)
 ) %>% gather(“date_diff”, “value”, 1:length(.)) %>%
 mutate(value = round(value, 4))
~~~

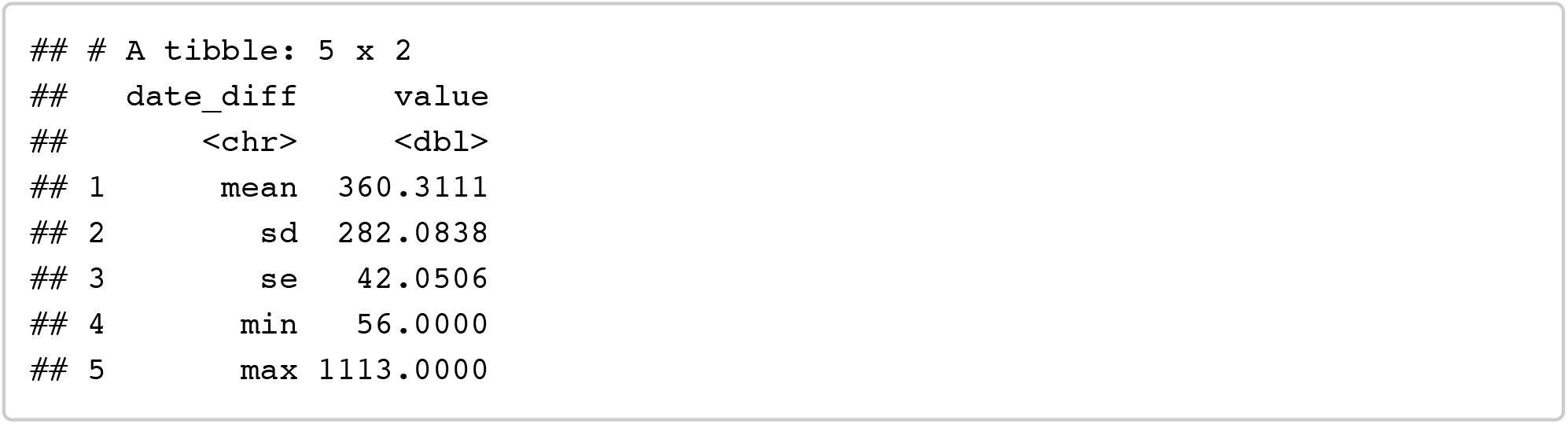

#### Analyses H4: Pill-switch

Within-subject change in pill use: (not considering partnership status)

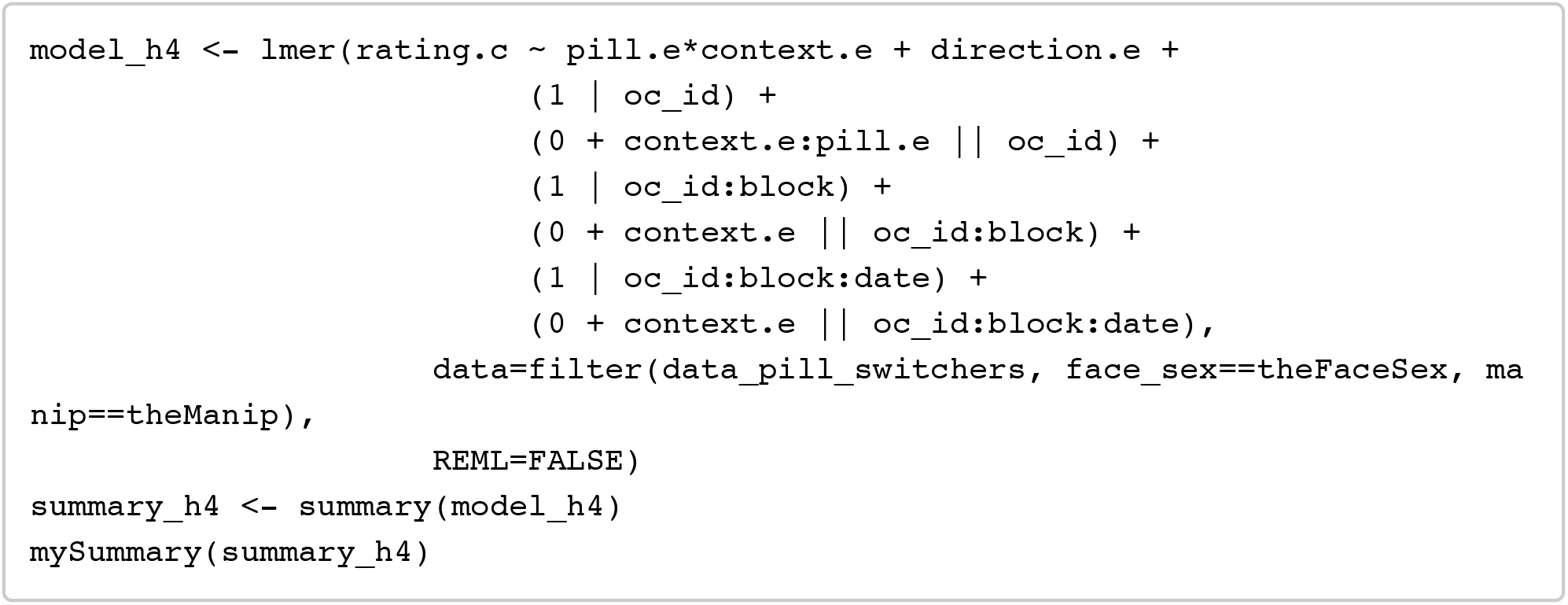

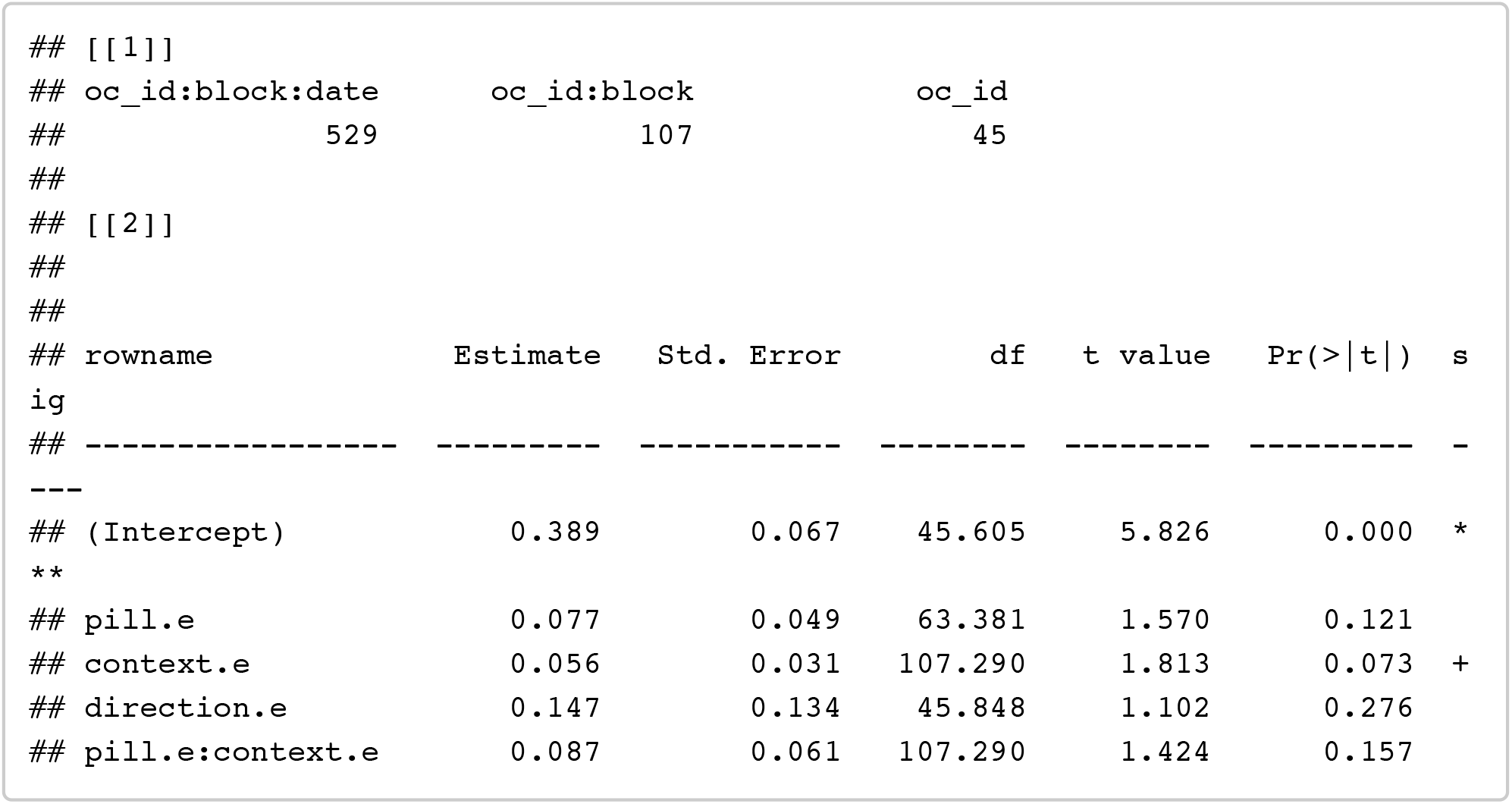

~~~
confint(model_h4, method = confint_method) %>% as.data.frame() %>% rownames_to _column() %>% filter(!is.na(‘2.5 %’))
~~~

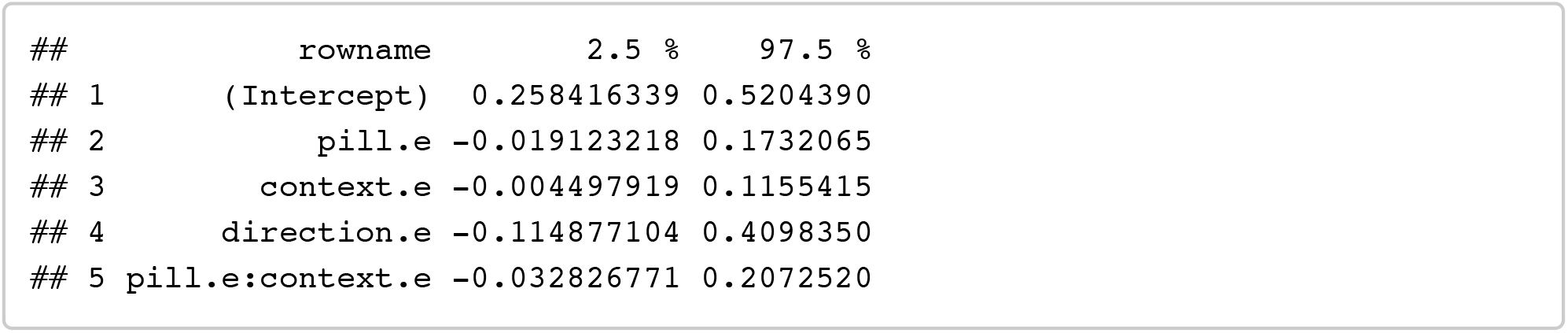

#### Analyses H4p: Pill-switch (+ partnership status change)

Within-subject change in pill use: (considering possible effects of change in partnership status)

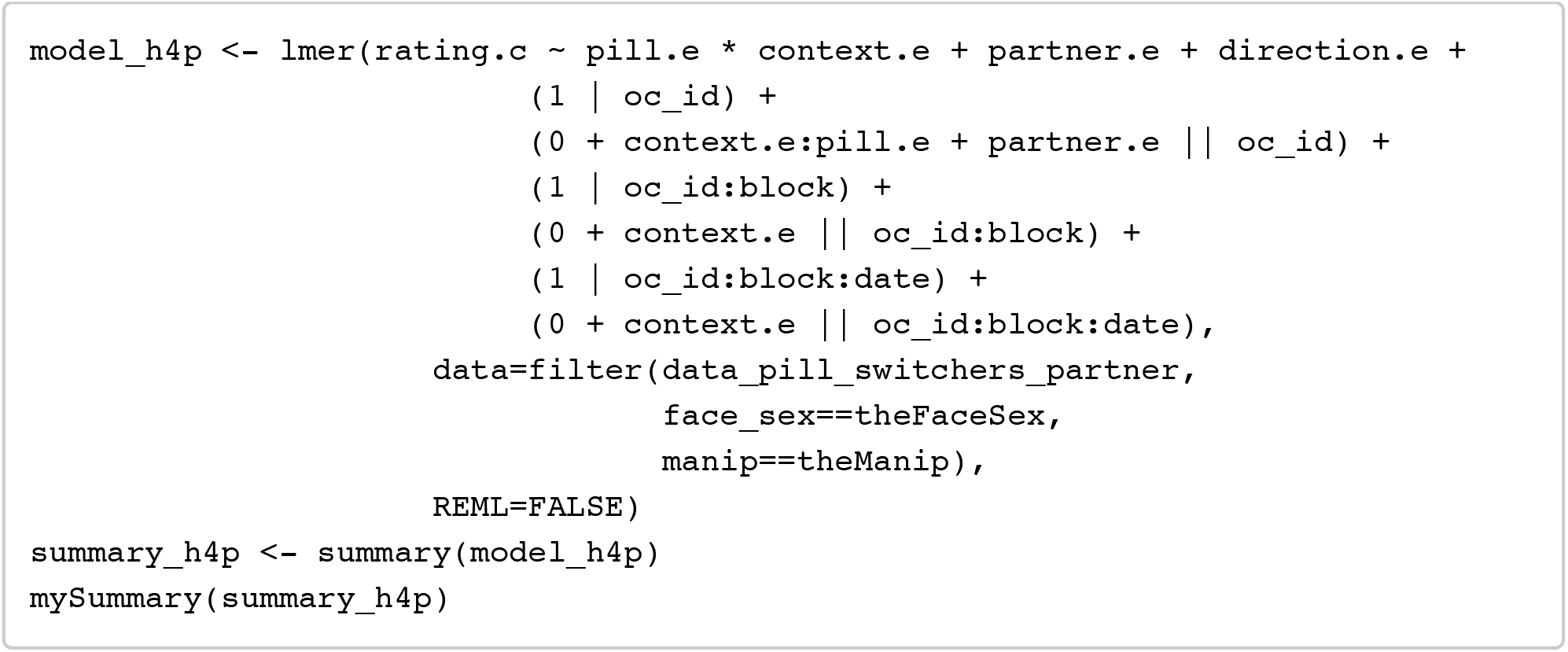

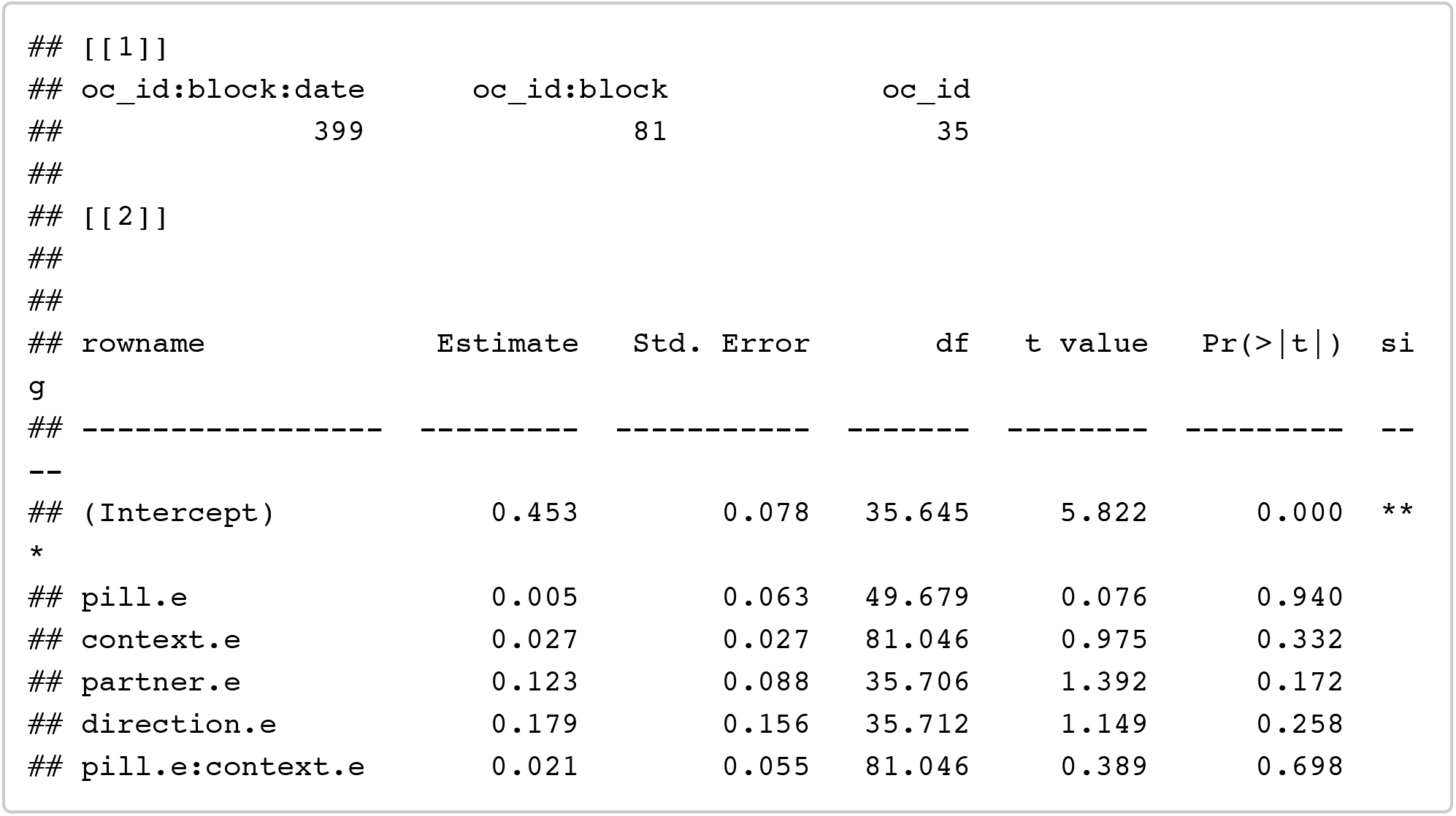

~~~
confint(model_h4p, method = confint_method) %>% as.data.frame() %>% rownames_t o_column() %>% filter(!is.na(‘2.5 %’))
~~~

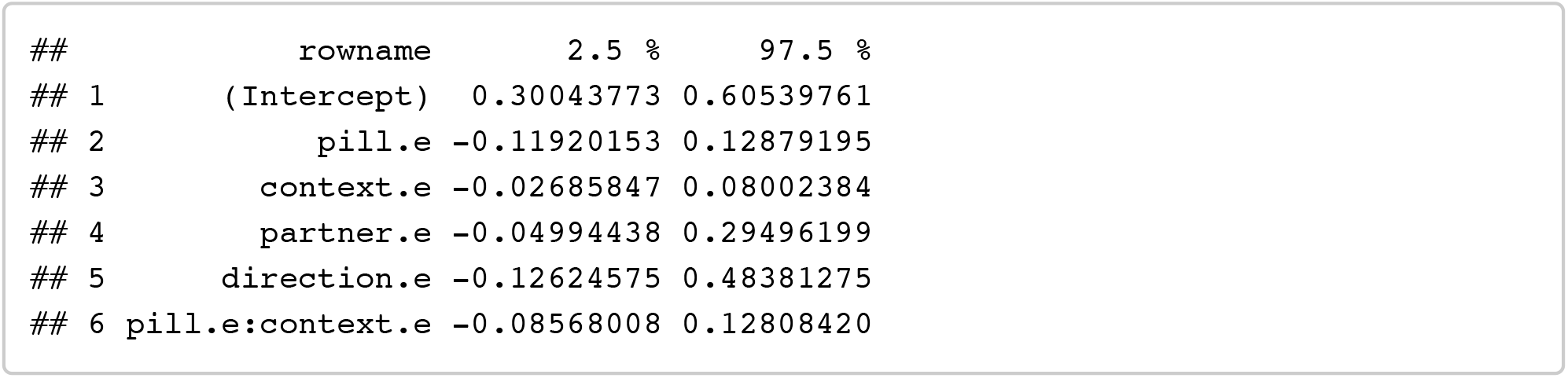

## Footnotes

1 The only significant effect of salivary hormones on masculinity preferences that we observed was an interaction between testosterone and cortisol. This was not an a priori prediction and we suggest is a false positive. This result is also not a prediction of the hypothesis that women’s masculinity preferences are affected by changes in fertility.

